# Osmotic Coupling of Mechanics and Epigenetics Regulates Chromatin Phase Organization

**DOI:** 10.64898/2026.10.29.762828

**Authors:** Aayush Kant, Monika Dhankhar, Rohit Joshi, Zixian Guo, Vinayak Vinayak, Diana I. Cruz, Kedi Wang, Ivan V. Surovtsev, Megan King, Abhijit Majumder, Jung-Hwan Lee, Hae-Won Kim, Elizabeth Hinde, Guilherme PF Nader, Vivek B. Shenoy

## Abstract

Cells experience epigenetic, osmotic, and mechanical perturbations that reorganize chromatin, yet how these distinct cues are integrated into a common physical response of the nucleus remains unclear. Here, we identify nuclear hydration as a central state variable that links microenvironmental perturbations to chromatin phase organization and nuclear mechanics. We develop a non-equilibrium mesoscale model that couples chromatin elasticity, water transport, and epigenetic reaction-diffusion kinetics. In this framework, chromatin behaves as a deformable polymer that segregates into coexisting water-rich euchromatic and water-poor heterochromatic phases, whose compositions and morphologies are regulated by hydration-dependent swelling, elastic strain, and histone-mark interconversion. The model predicts that epigenetic perturbations, osmotic shocks, and volume-changing mechanical loads reorganize chromatin by shifting intranuclear water content, nuclear volume, and euchromatin-heterochromatin phase balance. It further predicts that anisotropic forces impose directional strain, driving heterochromatin elongation, alignment, and lamellar transitions. Quantitative comparisons with super-resolution imaging, AFM and chromatin-accessibility measurement - across pharmacological, osmotic, and mechanical perturbations including confinement and micropatterning - support these predictions. Together, our results establish a chemo-mechano-osmotic framework in which nuclear hydration integrates biochemical and biophysical cues to regulate chromatin organization across scales.

## 1 INTRODUCTION

Cells continuously sense mechanical, osmotic, and biochemical cues from their microenvironment, and these signals influence proliferation, migration, metabolism, differentiation, and tissue morphogenesis ^1^. The nucleus is a central organelle in this response because it is both a mechanical force transducer and a regulatory compartment that organizes the genome ^2^. A key determinant of nuclear function is the spatial organization of chromatin into transcriptionally permissive, relatively open euchromatin (EC) and repressive, compact heterochromatin (HC). These chromatin states are regulated in part by post-translational histone modifications, including methylation marks such as H3K9me3 and H3K27me3, and acetylation marks such as H3K9ac ^3,4^. However, EC and HC are not fixed structural compartments, but are dynamically reorganized by the physical and chemical environment of the cell.

Environmental cues reach the nucleus through multiple coupled pathways. Mechanical forces generated at the cell-matrix interface are transmitted through the cytoskeleton and the linker of nucleoskeleton and cytoskeleton (LINC) complex, deforming the nuclear envelope and chromatin ^2^. Mechanical cues also regulate nucleo-cytoplasmic transport of transcriptional and epigenetic regulators, including YAP/TAZ and HDAC3, providing a mechano-chemical route for chromatin regulation ^5^. In parallel, the nucleus exchanges water and ions with the surrounding cytoplasm, allowing osmotic and ionic perturbations to alter nuclear volume, chromatin compaction, and molecular transport ^6,7^. Consistent with these pathways, mechanical deformation, osmotic stress, confined migration and altered epigenetic regulation have all been shown to reorganize chromatin across multiple length and timescales, including heterochromatin condensation, altered chromatin accessibility, changes in nuclear stiffness and persistent mechanical memory ^8–11^. Yet it remains unclear when these diverse perturbations converge on shared physical state variables, such as nuclear hydration, and when they reorganize chromatin through hydration-independent mechanical pathways.

Polymer models across length-scales have explained how epigenetic modification of histone marks and chromatin interactions shape genome organization ^12^, while phase-separation models have provided insight into the coexistence of transcriptionally distinct chromatin states ^13–15^. Separately, nuclear mechanics and osmotic models have described how forces and water exchange regulate nuclear volume and deformation ^5,6,8,10^. However, mesoscale chromatin is not simply a passive liquid-like mixture: compact heterochromatin and open euchromatin differ in hydration, stiffness and swelling behavior. Thus, chromatin phase organization should depend not only on histone-mark-dependent interactions, but also nucleoplasmic water transport and the elastic cost of maintaining water-rich and water-poor phases within a deformable polymeric material.

Here, we develop a thermodynamically consistent non-equilibrium mesoscale model that couples chromatin elasticity, nucleoplasmic water transport, and epigenetic reaction-diffusion kinetics. In this framework, EC and HC emerge as coexisting phases of a deformable chromatin polymer, with EC forming a water-rich swollen phase and HC forming a water-poor compact phase. Nuclear hydration thereby provides a common route by which epigenetic, osmotic and volume-changing mechanical perturbations regulate EC-HC phase balance, HC domain size, nuclear volume, chromatin accessibility, and nuclear stiffness. At the same time, the model predicts that anisotropic stresses can act through an additional route by imposing directional strain that elongates and aligns HC domains, even when overall nuclear hydration or phase abundance is preserved. We test these numerical predictions using complementary perturbations that alter epigenetic state, osmotic potential, nuclear volume, and anisotropic strain, with quantitative experimental comparisons in chromatin reorganization via multiple imaging and sequencing modalities. Together, this theory-experiment framework explains how biochemical and biophysical cues regulate chromatin phase organization through coupled hydration-dependent and strain-dependent mechanisms.

## 2 RESULTS

### 2.1 Osmotically partitioning establishes EC-HC phase coexistence

To determine how nuclear-scale chromatin architecture emerges from coupled biochemical and biophysical regulation, we developed a mesoscale continuum model that couples nucleoplasm transport, epigenetic reaction–diffusion dynamics, and large-deformation chromatin mechanics (Figure 1a; Section 4, Methods; Supplementary Section S1.1-S1.2). Chromatin organization is described by two spatiotemporally evolving fields: the local <u>n</u>ucleoplasm volume fraction (*φ_n_*) and an epigenetic order parameter representing the relative abundance of <u>m</u>ethylated and <u>a</u>cetylated histones (*φ_d_* = *φ_m_* − *φ_a_*). Together with the deformation field of the chromatin polymer, these variables determine the energetic and kinetic evolution of chromatin in the nucleus.

**Figure 1:**
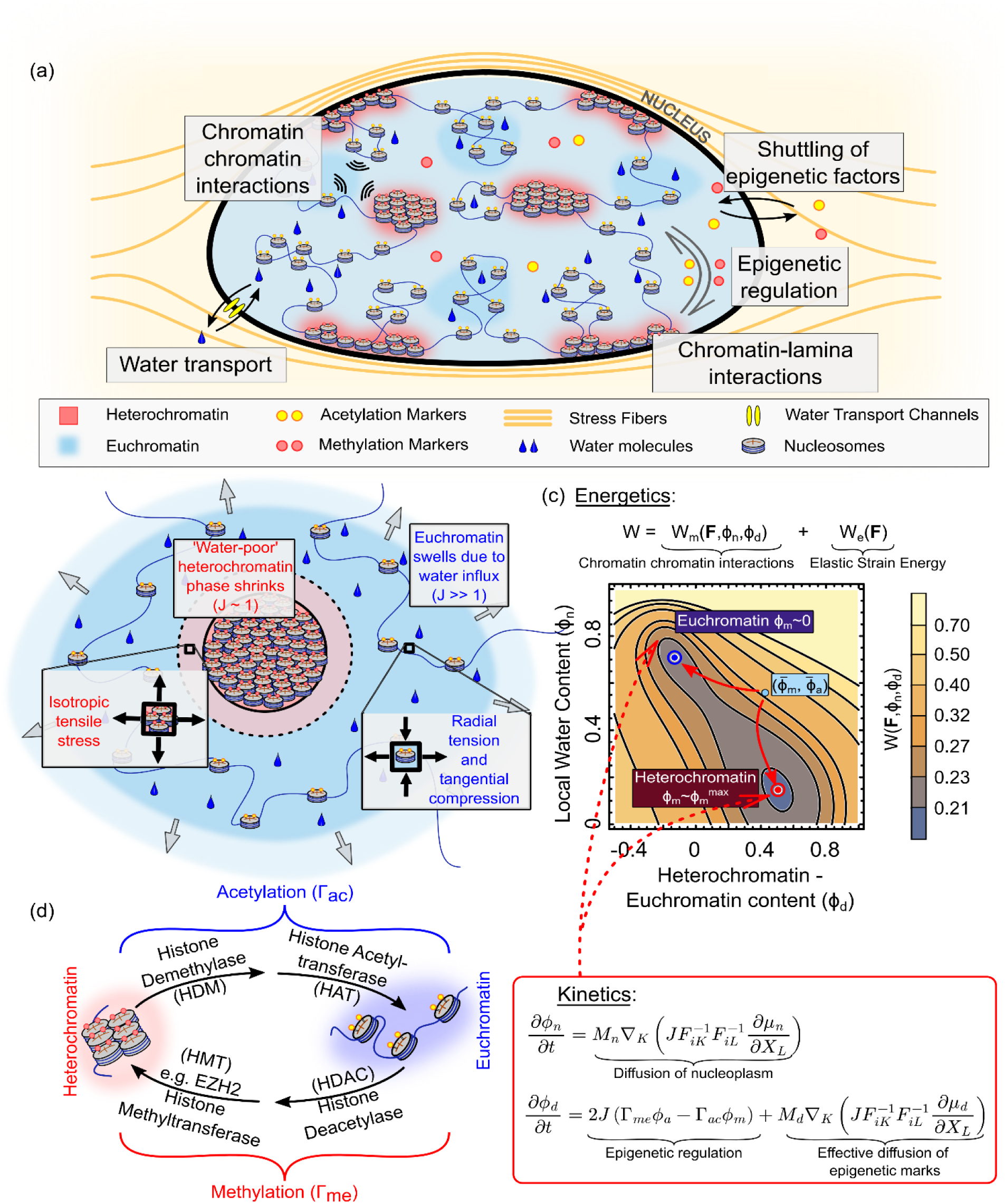
Schematic representation of the model ingredients. (a) A schematic diagram showing the components of mathematical model of chromatin organization. We consider the energetics ingredients such as chromatin-chromatin interactions and the chromatin lamina interactions, in addition to the kinetics of nucleoplasm diffusion, and epigenetic regulation. The nucleus can exchange water with the cytoplasm in response to environmental cues, while nucleo-cytoplasmic shuttling of epigenetic factors is also incorporated. Chromatin is treated as a non-linear visco-hyperelastic polymer (b) Local presence of space-occupying nucleoplasm increases the local volumetric swelling. Thus, euchromatic phase swells more than heterochromatin. The differential swelling in chromatin phases causes volumetric strain mismatch resulting in shear strains especially in the softer euchromatin phase. Note that *J* = det(***F***) denotes the local volume of each point (Eq. S1, SI), where ***F*** denotes the deformation gradient which tracks the deformation of all points in the nucleus at all times. (c) The contour plot of the free energy density (Eq 1) in the bulk of the nucleus shows a characteristic double well feature. Compared to the free energy density in the absence of elasticity (Figure S2), the plot shows the distinctive contributions of the strain energy. The two wells correspond to the euchromatin (blue) and the heterochromatin (red) phases. Any initial configuration (light blue circle) will spontaneously phase separate into these wells so as to reduce the free energy of the nucleus. The dynamics of the organization is governed by the diffusion of nucleoplasm, epigenetic marks and the active reaction kinetics of the epigenetic regulation. (d) Epigenetic factors catalyze the reactions leading to interconversion of EC and HC. The acetylation reaction occurs at an overall rate of Γ*_ac_* and involves the removal of methylation marks (via histone demethylase, HDMs) and addition of acetylation marks (via histone acetyltransferases, HATs). Conversely the methylation reaction occurs at an overall rate Γ*_me_* and involves removal of acetylation marks (via histone deacetylase, HDACs) and addition of methylation marks (histone methyltransferases, HDMs).

The chromatin free-energy landscape arises from several competing physical contributions (Figure 1c; Section 4, Methods; Section S1.4). Histone-mark-dependent chromatin-chromatin interactions favor segregation of compositionally distinct chromatin states and generate the underlying double-well structure of the free energy. Nucleoplasm content is coupled to local chromatin volume through the incompressibility constraint (derived in Section S1.3), such that water uptake causes local swelling whereas water loss produces compaction (Figure 1b). Because chromatin is treated as a deformable polymer, these hydration-dependent volume changes incur an elastic strain-energy cost. In addition, interfacial energy due to dissimilar neighboring nucleosomes penalizes boundaries between compositionally dissimilar EC and HC regions, while preferential interactions between HC and the nuclear lamina favor peripheral heterochromatin organization. The competition among these energetic terms produces two stable bulk minima corresponding to a water-rich, swollen EC state and a water-poor, compact HC state (Figure 1c).

Starting from a nearly homogeneous distribution of nucleoplasm and histone marks with small spatial perturbations, the system evolves toward these two preferred states. The parameter choice used for the simulations is described in detail in Section S1.8 (Table 1). Nucleoplasm preferentially partitions into the EC phase, causing it to swell as it takes up space-occupying water molecules, whereas HC expels nucleoplasm and becomes compact. This coupled epigenetic and osmotic segregation produces HC domains within a continuous EC-rich background in the nuclear interior (Figure 2a). At the nuclear boundary, the additional preferential interaction between HC and lamina-associated proteins stabilizes peripheral HC, giving rise to lamina-associated domains ^16^. Thus, interior HC domains emerge from the bulk phase-separation mechanism, whereas peripheral HC is additionally stabilized by chromatin–lamina interactions.

**Figure 2:**
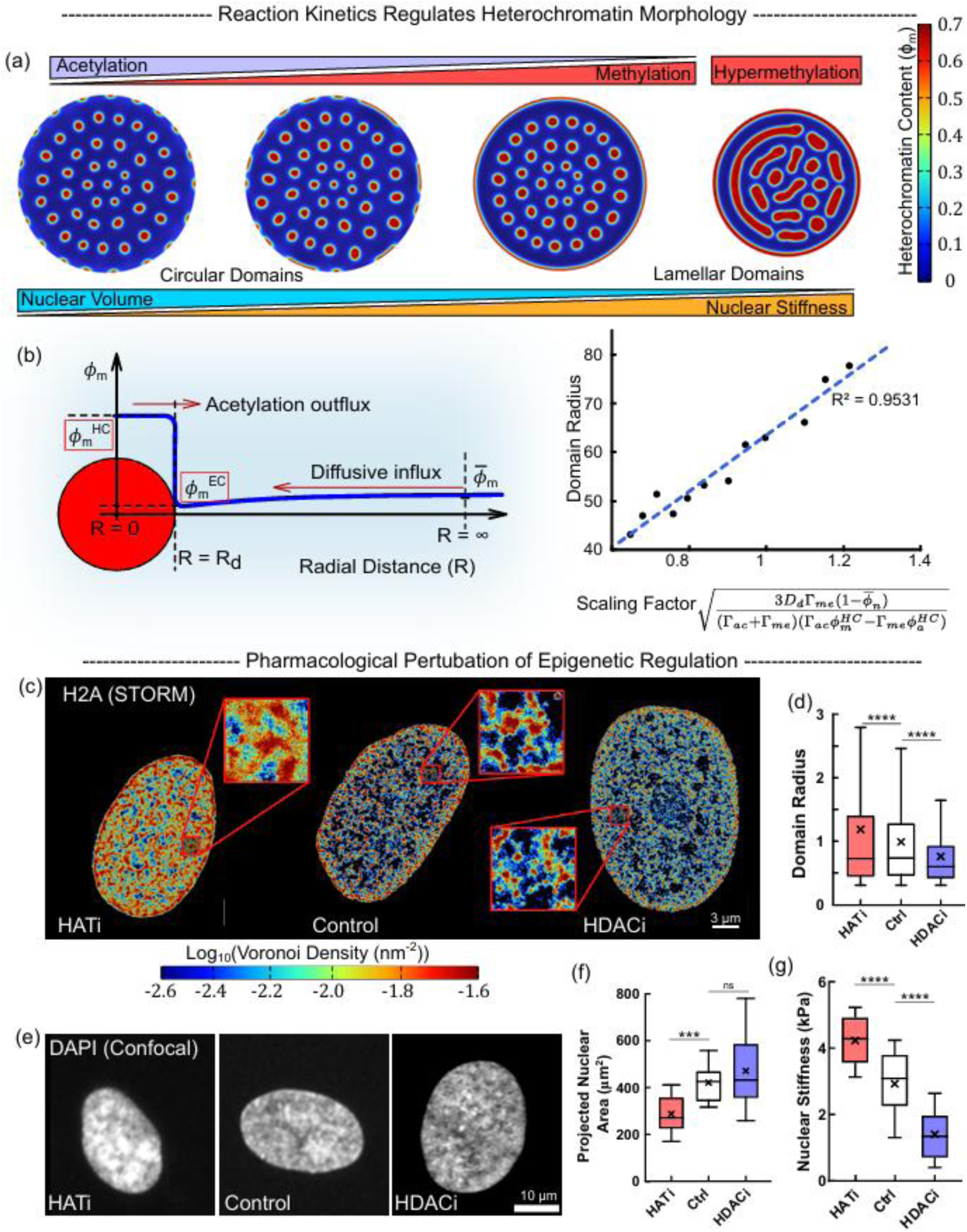
Epigenetic reaction kinetics determines the size of heterochromatin domains. (a) Numerical predictions of steady-state chromatin organization as the epigenetic reaction rates of histone methylation and acetylation are altered. Increasing acetylation reduces the size of heterochromatin domains, while elevated methylation increases their sizes, forming more non-circular lamellar domains. (b) The size of the heterochromatin domains observed via numerical simulations (red dots) follows the theoretically predicted scaling relationship (Eq 2, blue dashed line). (c) Super-resolution STORM images of hMSC nuclei under HAT inhibition, control, and HDAC inhibition treatment conditions. The images are color-coded according to the density of Voronoi tessellations of H2A localizations (n=15 nuclei for each condition). Scale bar: 3 *μ*m. (d) Distributions of interior heterochromatin domain radii observed experimentally in nuclei under HAT inhibition, control, and HDAC inhibition treatment conditions (n=16593 loci for control, 18563 loci for HDACi, 13102 loc for HATi; unpaired two-tailed t-test, p<1e-10 for control vs HATi, and control vs HDACi). (e) Confocal images of nuclei under HAT inhibition, control, and HDAC inhibition treatment conditions. Scale bar: 10*μ*m (f) Quantification of two-dimensional nuclear area on the maximum intensity projected (MIP) after HAT inhibition, control, and HDAC inhibition treatment conditions (n = number of nuclei: 31 HDACi, 15 control and 24 HATi; unpaired two-tailed t-test, p=7e-6 for control vs HATi, p=0.13 for control vs HDACi). (g) Quantification of nuclear stiffness after HAT inhibition, control, and HDAC inhibition treatment conditions (n = number of nuclei: 27 control, 30 HATi, 26 HDACi; unpaired two-tailed t-test, p=2e-6 control vs HATI, p=2e-7 control vs HDACi).

A defining feature of the resulting coexistence is the large difference in hydration between the coexisting chromatin phases. The water-rich EC phase undergoes substantially greater volumetric swelling than the relatively dehydrated HC phase (Figure 1b). Because the two phases are mechanically connected within the chromatin polymer, this differential swelling necessarily generates an elastic mismatch. Specifically, the expansion of EC region exerts isotropic tensile stress on the HC domain opposing its compaction and resisting expulsion of water from HC. Conversely, the contraction of HC deforms the surrounding EC. The EC phase experiences a radial tension (inward pull) and tangential compression (inward motion of points in EC brings them closer to each other), leading to the emergence of deviatoric (shear) stresses in the EC region particularly near the EC-HC interface. Simulations (Figure S17) and the simplified analysis of an HC domain embedded in EC (Supplementary Section S4-S5) show that these stresses feed back onto the effective free-energy landscape. The elastic strain energy due to elastic mismatch of the two chromatin phases effectively competes with chromatin-chromatin interactions which favors distinct EC and HC compositions. Consequently, elasticity pushes the coexisting chromatin phases to be compositionally closer, i.e. the EC phase becomes less dilute, while the HC phase becomes less compact. Thus, elastic strain does not merely arise as a passive consequence of phase separation, but directly modifies the compositions and stability of the coexisting phases.

Thus, EC–HC organization is not determined by histone-dependent interactions alone. Osmotic partitioning establishes distinct hydration states, while the elastic cost of accommodating their differential swelling frustrates complete segregation and modifies the compositions and stability of the coexisting phases (as derived in Section S7). Chromatin–lamina interactions further bias HC toward the nuclear periphery without changing this underlying bulk coexistence mechanism. Hydration and elasticity therefore establish the physical landscape on which EC and HC organize. They do not, however, explain why HC domains remain at a finite characteristic size. In Section 2.2, we show that this length scale emerges from the non-equilibrium balance between epigenetic interconversion and diffusive transport.

### 2.2 Epigenetic flux balance selects a hydration-dependent heterochromatin length scale

#### Epigenetic reaction-diffusion arrests HC domain coarsening

Building on Section 2.1, we next ask how the characteristic size of HC domains emerges once water-rich EC and water-poor HC phases have formed. In a passive phase-separating system, compact HD domains are expected to coarsen progressively to reduce interfacial energy. In our model, however, chromatin organization is maintained away from thermodynamic equilibrium by active epigenetic interconversions between acetylated and methylated histone states (Section S3). These reactions continuously reshape the local balance between EC-like and HC-like chromatin, thereby arresting, coarsening and selecting a steady-state HC domain size (Section S5.3). As HC domains nucleate, gradients in the methylated histone concentration drive an inward diffusive flux of methylated histones promoting the growth of the compact, water-poor HC domain (Figure 2b). This growth is opposed by acetylation-driven conversion and outward redistribution of acetylated histones into the surrounding water-rich EC phase (Figure 2b). We find that the steady-state domain radius is therefore determined by a balance of opposing fluxes across the HC–EC interface, as derived in Section S5.3,

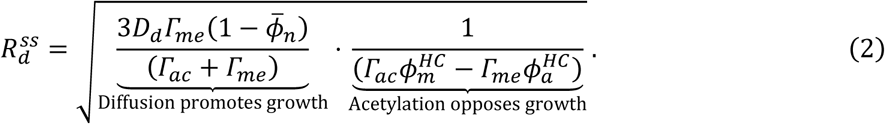

where, 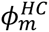 and 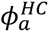 are the volume fractions of methylated histones and acetylated histones in the HC phase, ϕ̅*_n_* is the overall water content in the nucleus (Section S1.3), Γ*_me_* and Γ*_ac_* are the overall reaction rates of conversion of methylated histones into acetylated histones and vice versa (Figure 1d). Importantly, the phase compositions entering this scaling relation are not fixed constants (Section S4). They are set by osmotic partitioning of nucleoplasm and by the elastic cost of differential swelling between EC and HC, established in Section 2.1. Thus, HC domain size is not purely reaction-diffusion length scale; it is a hydration-dependent chemo-mechano-osmotic length scale.

The scaling relationship (Eq. 2) predicts that increasing *Γ_ac_* (*Γ_me_*) leads to smaller (larger) HC domains. Starting from a steady-state chromatin distribution (Figure 2a, center), simulations show that higher methylation or acetylation rates correspondingly increase or decrease HC domain size respectively (Figure 2b, right). Thus, our model predicts that the characteristic size of HC domains is selected by epigenetic flux balance operating on an osmotically and mechanically regulated phase landscape; any perturbation that changes nuclear hydration should therefore shift HC domain size.

#### Pharmacological perturbations validate predicted HC domain-size shifts

To experimentally validate our model predictions, we pharmacologically perturbed epigenetic reactions in bone marrow-derived human mesenchymal stem cells (hMSCs) by treating them with histone deacetylase inhibitors (HDACi: trichostatin A (TSA) and valproic acid (VPA)) and histone acetyltransferase inhibitor (HATi: anacardic acid, ANA). HDACi treatment reduces the rate of histone deacetylation, contributing to a decrease in the histone methylation rate *Γ_me_* in our model (Figure 1d), while HATi inhibits histone acetylation rate *Γ_ac_*. To visualise and quantify the in-vitro nucleus-wide chromatin reorganization in response to these epigenetic inhibitors, we used stochastic optical reconstruction microscopy (STORM) to image histone H2A density in hMSC nuclei after HDACi and HATi treatment (Figure 2c). High H2A density regions correspond to tightly packed HC (red region in Figure 2c), while the low-density regions correspond to EC ^15–17^. The STORM images clearly show that the HATi increases the overall HC content (Figure 2c), while HDACi reduces them.

We next quantified the radius of HC domains from the STORM density maps. In agreement with the predicted scaling of HC domain size with epigenetic reaction kinetics, HATi treatment increased the mean HC domain radius (∼1.2 times), whereas HDACi treatment reduced it (∼0.8 times, Figure 2d). Together, these results support the pred iction that HC domains size is selected by epigenetic reaction-diffusion operating on a hydration- and elasticity-regulated EC-HC phase landscape.

#### Hypermethylation leads to morphological transition of heterochromatin into lamellar domains

In addition to altering HC domain size, our model predicts that epigenetic reactions can reshape domain morphology. Specifically, under conditions of very high methylation rates, the abundance of methylated histones in the nucleus reduces effective nuclear hydration. Thus, hypermethylation promotes the progressive enrichment of densely packed chromatin, leading to the crowding of enlarged water-poor HC domains in the nuclear interior. The favorable energetic interactions between methylated histones promotes the coalescence of the HC domains – inducing a morphological transition from circular to lamellar HC domains (Figure 2a).

To test this prediction, we examined chromatin organization in tenocytes from tendinosis patient samples, a biologically relevant setting in which epigenetic dysregulation is well-documented ^9^. Diseased tenocytes exhibit elevated levels of histone methyltransferase (e.g., enhancer of zeste homolog 2, EZH2) which is incorporated in our model as an increased histone methylation rate. STORM imaging of nuclei under these hypermethylated conditions reveals HC domains with elongated lamellar morphology ^9^, consistent with the model prediction that strong bias toward the water-poor HC phase can drive both domain growth and shape transitions (Figure S18).

### 2.3 Nuclear hydration couples epigenetic chromatin phase transitions to nuclear mechanics

#### Epigenetic regulation redistributes water to control nuclear volume

In Sections 2.1 and 2.2, we predicted that epigenetic reactions directly regulate the relative abundance and mesoscale organization of the water-rich EC and water-poor HC phases. We next ask whether this hydration-dependent reorganization also controls macroscopic nuclear mechanics. By shifting the chromatin phase balance, epigenetic perturbations in our framework directly drive water efflux or influx across the permeable nuclear envelope to maintain osmotic equilibrium with the cytosol. The changes in the nuclear volume and mechanics then emerge as a consequence of this osmotic redistribution, as we discuss below.

Epigenetic kinetics-driven conversion of water-rich EC into water-poor HC generates an effective excess of intranuclear water, which is expelled through the nuclear envelope pores. Thus, the promotion of HC reduces the overall nuclear hydration. Because water occupies space within the nucleus, while the total chromatin content remains fixed, this loss of intranuclear water leads to a reduction in nuclear volume, under the incompressibility constraint (second term in Eq. 1; Section S1.3). Accordingly, our model predicts that an increase in histone methylation rates reduces nuclear volume. Conversely, epigenetic conversion towards EC draws water into the nucleus, increasing the nuclear hydration and volume. The change in nucleus volume driven by altered histone acetylation and methylation rates, as predicted by our simulations, is illustrated Figure 2a and quantified in Figure S19.

We validate these predictions by pharmacologically perturbing epigenetic kinetics in hMSCs followed by DAPI staining and Airyscan confocal imaging (Figure 2e) to quantify the nuclear size and HC content. For a quantitative comparison with the 2D computational analysis the 3D image stacks were converted to 2D via maximum intensity projection (MIP) in ImageJ. The flattened 2D projection was used to calculate the projected nuclear area. Regions of high DAPI intensity, which correspond to locally increased DNA concentration were used to identify HC phase, and HC content was calculated as percent of projected nuclear area occupied by HC. Cells treated with the histone acetyltransferase inhibitor (HATi) anacardic acid (ANA), which promotes water-poor HC phase, exhibit a reduction in nuclear area to approximately 0.68 times that of control cells (Figure 2e,f). Consistent with this hydration loss mechanism, the HC content (percent of projected nuclear area occupied by HC) increases by nearly 2.61-fold (Figure S20). These data were used to set up the simulation parameters (Table 1, Section S1.8) calibrated to the experimentally seen relationship between nuclear volume and HC fraction (Figure 2f).

We then quantitatively validate the model parameters using an independent perturbation. Treatment with the histone deacetylase inhibitor (HDACi) valproic acid (VPA), which promotes water-rich EC phase, results in an increase in nuclear volume to approximately 1.12 times that of control cells. Our model predicts that such swelling corresponds to a reduction of HC fraction to approximately 0.68 times control. Quantifying the HC fraction from the DAPI-stained nuclei shows that the HC content is reduced to 0.69 times that in control nuclei (Figure S20), closely matching the simulation-predicted trend. (Figure 2f). As a further confirmation of global chromatin condensation, we evaluate the chromatin condensation parameter (CCP) which is an image-based quantitative metric to quantify the extent of chromatin condensation in the nucleus ^18^. We observe that chromatin condensation increases with ANA (HAT inhibition) and decreases with VPA (HDAC inhibition) treatments (Figure S20).

Together, these results support the model prediction that epigenetic regulation controls nuclear volume through hydration-mediated chromatin phase transitions. By shifting the balance between the water-rich EC and water-poor HC phases, epigenetic reactions drive nucleo-cytoplasmic osmotic redistribution, making nuclear volume a physical readout of chromatin phase balance.

#### Hydration-dependent chromatin reorganization governs nuclear stiffness

Chromatin has been previously shown to be a key contributor to the mechanical stiffness of the nucleus ^11,19^ with HP1*α*-crosslinked HC regions demonstrating higher local stiffness than the water-rich EC phase ^20^. Hence, epigenetic driven changes in HC-EC proportions are expected to alter nuclear mechanical properties.

Applying a constant uniaxial load (as described in Section S8), we compute the effective nuclear stiffness from the slope of the stress-strain response (Figure S12). Our model predicts a monotonic increase (decrease) in nuclear stiffness with increasing HC (EC) content. Mechanistically, this stiffening arises because epigenetic conversion towards HC increases the fraction of condensed, stiffer HC, while conversion towards EC enriches the nucleus in more hydrated softer chromatin. We validated these predictions experimentally using atomic force microscopy (AFM) to measure the stiffness of control, HDACi- and HATi-treated nucleus. Consistent with the model, HDACi treatment softens the nucleus, whereas HATi treatment increases overall nuclear stiffness (Figure 2g). These observations support the view that changes in the nuclear mechanics upon changes in epigenetic regulation can be prominently attributed to hydration-dependent chromatin reconfiguration.

We have previously shown that biophysically relevant changes in the cell microenvironment, which affects the cell contractility, can alter the nucleo-cytoplasmic partitioning of epigenetic regulators such as HDAC3 ^5,16,21^. Thus, beyond pharmacological epigenetic perturbations, we also tested our model predictions under biophysically relevant epigenetic regulation mediated by substrate stiffness ^5,16^ as described in the SI (Section S10, Figure S16). Consistent with our predictions, substrate stiffness-dependent changes in contractility drive chromatin reorganization leading to corresponding changes in nuclear volume analogous to those observed under pharmacological epigenetic perturbations. Together, these results demonstrate that pharmacological and mechanical perturbations, convergently via epigenetic reactions, can control macroscale nuclear properties – nuclear volume and stiffness – through a common osmotic pathway. Changes in histone methylation or acetylation alters the relative abundance of EC and HC phases, thereby regulating intranuclear water content, nuclear volume and stiffness. Osmotic regulation, thus, emerges as a direct mechanistic link between chromatin organization and nuclear mechanics.

### 2.4 Osmotic perturbations directly alter chromatin condensation and HC domain sizes

In Section 2.3, we have shown that intranuclear water content plays a central role in regulating the chromatin reorganization and nuclear-scale properties under epigenetic perturbations. Since osmotic perturbations directly modulate water exchange between the nucleus and the cytosol ^6,7,22,23^, we next examine chromatin reorganization under osmotic stress and compare our model predictions with experimental observations.

#### Osmotic loading alters the composition of coexisting chromatin phases and HC domain size

Osmotic perturbations, such as those caused by changes in osmolyte concentrations, are incorporated in our model via the cytosolic chemical potential of water, 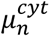, prescribed along the boundary of the steady state nucleus (Section S1.8). Higher cytosolic water potential relative to that within the nucleus, results in an influx of water into the nucleus from the cell, and vice versa.

To simulate osmotic perturbations, we alter 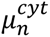 and allow the chromatin to reorganize, reaching a new steady state. Our results (Figure 3a) demonstrate that osmotic forces can directly regulate the overall nucleus volume as well as the hydration of the coexisting HC and EC phases. In an iso-osmotic environment (i.e., 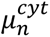 unchanged) no net water exchange occurs, and the chromatin organization stays unaltered (Figure 3a, center panel). In a hypo-osmotic environment, the cytosol becomes water-rich driving an influx of water into the nucleus (Figure S21) leading to dynamic reorganization of the HC domains (Figure 3a, left). The HC phase becomes less favored, and the fraction of area occupied by the HC relative to the nucleus decreases as shown in Figure S21. This is accompanied by a decrease in the size of the HC domains (Figure 3b). Further the nuclear volume increases due to the water influx as compared to that under isotonic (unaltered) conditions (Figure S21). Conversely, in a hyper-osmotic environment (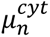 decreases), we observe efflux of water (Figure S21), reduction of nuclear volume (Figure 3a, Figure S21), an increase in the fraction of area occupied by the HC phase (Figure S21) and an increase in size of the HC domains (Figure 3b).

**Figure 3:**
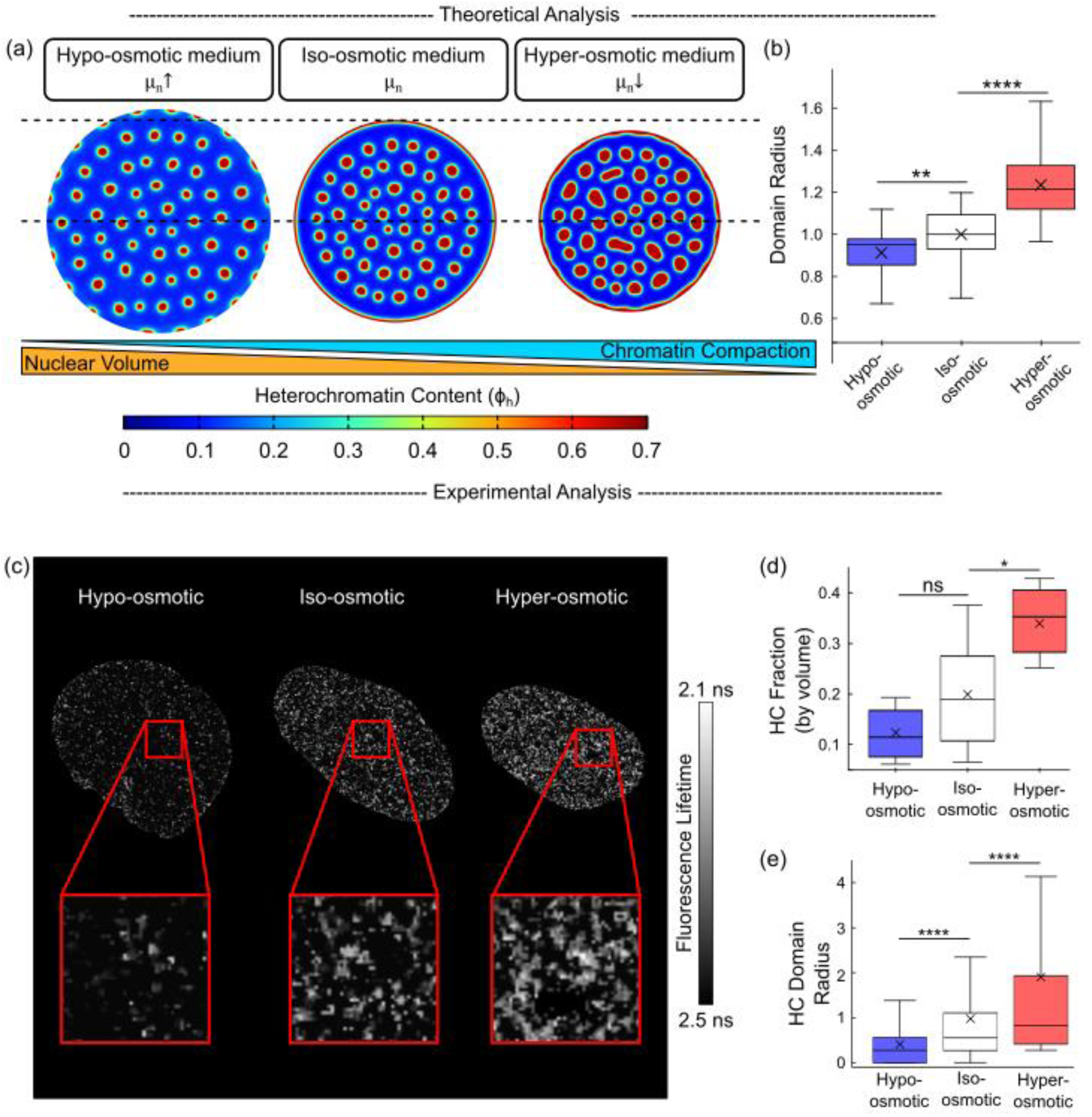
Osmotic stress on nucleus results in chromatin reorganization. (a) Simulations show the chromatin compaction and nuclear volume is affected by osmotic levels of surrounding medium. Increasing osmotic stress increases the chromatin compaction and heterochromatin domain size, while reducing the euchromatin region (blue) and the overall nuclear volume. (b) Quantification of HC domain sizes obtained from simulations (n = 49 domains in hypo-osmotic, 51 domains in iso-osmotic, and 41 domains in hyper-osmotic case; unpaired two-tailed t-test, p=0.002 for hypo- vs iso-osmotic, p=1e-13 for hyper- vs iso-osmotic). (c) FLIM-FRET images of HeLa nuclei subjected to hypo-osmotic, control (iso-osmotic) and hyper-osmotic loading. Brighter intensities indicate more efficient FRET efficiency and closer histones, representing compacted HC phase. (d) The fraction of nuclear area occupied by HC increases under hyper-osmotic loads and reduces under hypo-osmotic loads (n=7 nuclei for each case; unpaired two-tailed t-test, p=0.2 for hypo- vs iso-osmotic, p=0.02 for hyper- vs iso-osmotic), and (e) the HC domain size follows a similar trend (n = 4002 domains in hypo-osmotic, 4327 domains in iso-osmotic, and 4400 domains in hyper-osmotic case; unpaired two-tailed t-test, p=0 for hypo- vs iso-osmotic, p=0 for hyper- vs iso-osmotic).

#### FLIM-FRET of in-vivo hyper- and hypo-osmotic treated nuclei validates theoretical and numerical predictions

To experimentally validate these predictions, we analyzed chromatin reorganization in HeLa nuclei under osmotic stress using FLIM-FRET imaging (Figure 3c). FLIM-FRET reports the nanoscale chromatin compaction by measuring the energy transfer fluorescently labeled histones, with a higher FRET efficiency implying more densely packed histones indicative of HC phase ^24–26^. Consistent with model predictions, hyperosmotic shock increases chromatin compaction, resulting in a ∼1.7-fold increase in the fraction of nuclear area occupied by HC (Figure 3d), while hypo-osmotic conditions reduce HC region (∼0.8-fold, Figure 3d). Further, we quantify the sizes of the heterochromatin domains observed via the FLIM-FRET imaging (Figure 3e). We observe that the heterochromatin domains formed in nuclei undergoing hyper(hypo)-osmotic shocks are significantly larger by ∼1.8 times (smaller by ∼0.5 times) than those in nuclei experiencing iso-osmotic loads.

Together, these results demonstrate that osmotic perturbations directly regulate chromatin organization via nuclear hydration, by altering the balance between the coexisting HC and EC phases and regulating the HC domain size. Importantly, these findings establish nuclear water content as a central regulator of chromatin organization.

### 2.5 Tensile and compressive loads modify nuclear-wide chromatin compaction

#### Mechanical forces modulate the nuclear volume and composition of coexisting chromatin phases

In sections 2.1-2.4, we showed that nuclear hydration regulates the composition and relative abundance of the coexisting water-rich EC and water-poor HC phases, thereby controlling chromatin organization across scales. However, comparable changes in nuclear volume and thus hydration can be enforced via direct deformation of nucleus upon application of mechanical forces ^5,8^. Hence, we next ask whether external mechanical loading can influence chromatin reorganization via the nuclear hydration pathway.

To elucidate how mechanical forces influence chromatin organization, we subject the nucleus to equi-biaxial tensile and compressive loading (represented by blue and red arrows respectively in Figure 4a) and analyze chromatin organization once steady state is attained (Figure 4a, center panel). During these simulations we permit a free exchange of water across the nuclear membrane by maintaining a constant chemical potential 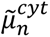 at the boundaries. In the absence of mechanical stress, the nucleus remains undeformed with no net water flux (Figure 4a, center; Figure S22), and no chromatin reorganization. Under compressive loading, the nuclear volume decreases (Figure 4a, left), reducing the space available for intranuclear constituents and thereby expelling water from the nucleus (Figure S22). Conversely, tensile loading increases nuclear volume (Figure 4a, right) and facilitates water influx (Figure S22). Thus, our model predicts that mechanical forces directly modulate intranuclear hydration by regulating nuclear volume.

**Figure 4:**
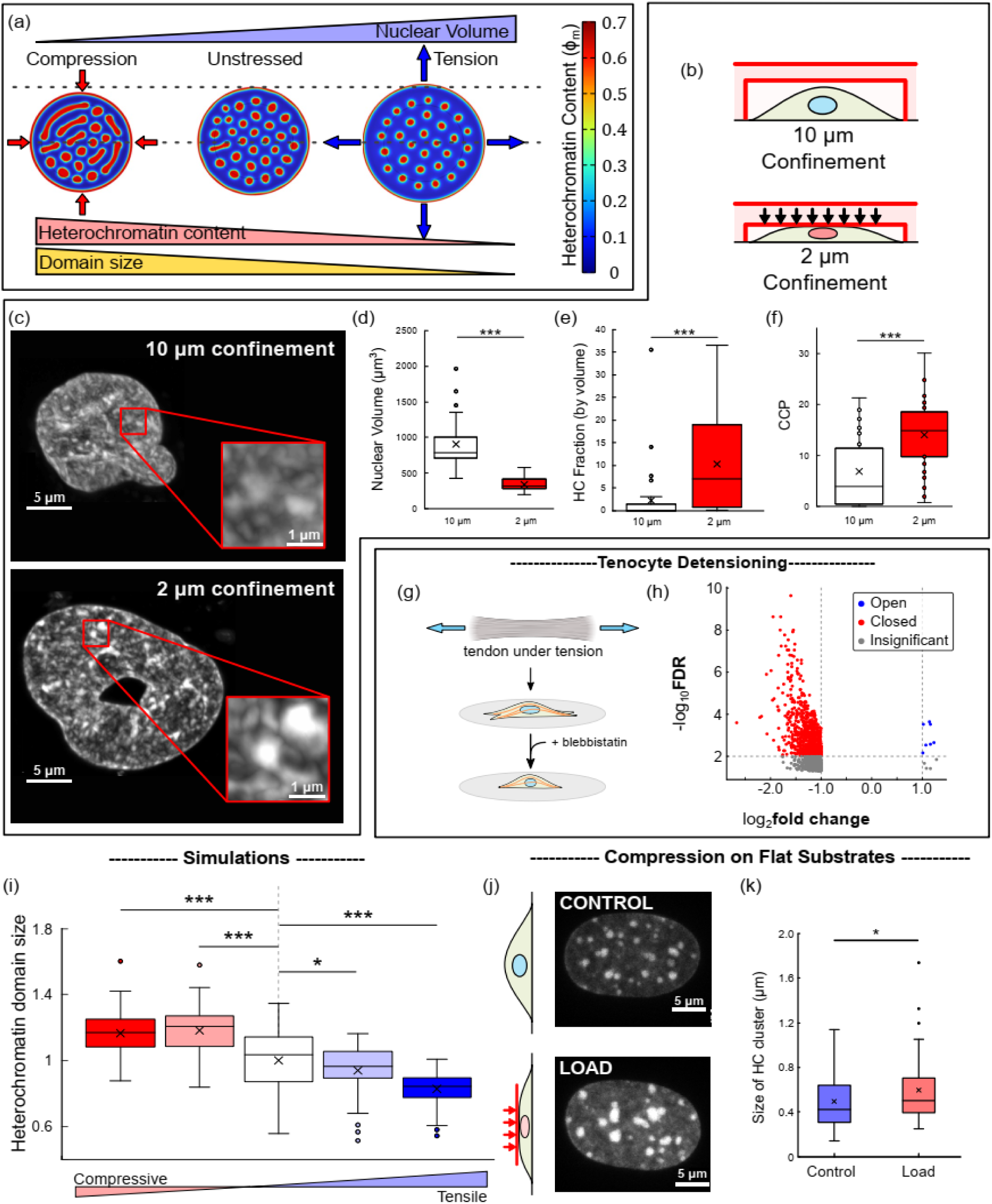
Mechanical compressive and tensile forces alter global chromatin reorganization. (a) Numerical modeling shows that the compressive forces increase the heterochromatin content while the nuclear volume reduces. Tensile forces reduce the heterochromatin content while the nuclear volume increases. (b) Schematic demonstrating the application of compressive loads on MDA-MB-231 cells cultured on flat with confinements of 10 *μm* and 2 *μm*, (c) with corresponding confocal images of DAPI-stained nuclei. Increasing confinement of the nucleus (d) reduces the nuclear volume (n=35 and 34 cells for 2*μm* and 10 *μm* confinement respectively; unpaired two-tailed t-test, p=1e-11), (e) increases the fraction of nuclear volume occupied by HC (unpaired two-tailed t-test, p=3e-4), and (f) increases the chromatin compaction quantified via CCP (unpaired two-tailed t-test, p=1e-5). (g) Schematic of ex-vivo tenocyte de-tensioning ^27^ via blebbistatin treatment. (h) Nuclei of de-tensioned tenocytes exhibit a reduced chromatin accessibility, suggesting that homeostatically tensed nuclei have a relatively open chromatin arrangement. (i) Quantifications of domain sizes observed in numerically predicted chromatin reorganization in nuclei after compressive (red) and tensile (blue) loads. Compression increases HC domain sizes, while tension results in formation of smaller domains. (j) Application of compressive forces (approximately 2.5 *μN* ^8^) alters the chromatin organization observed via confocal imaging (bottom panel) after DAPI staining in NIH3T3 fibroblasts. The chromatin reorganization after compressive loads and is accompanied by (k) an increase in size-scales of the dense HC clusters seen in the confocal images (n=75 and 76 HC clusters for control and load case respectively; unpaired two-tailed t-test, p=0.03).

Because nucleoplasm is predominantly associated with the diluted EC region, these hydration changes primarily alter the composition of EC. Compression reduces EC hydration making the EC phase denser (darker blue EC in Figure 4a, left panel), whereas tension increases EC hydration (lighter blue EC in Figure 4a, right panel). The HC phase, already relatively water-poor, undergoes comparatively smaller compositional changes. The non-equilibrium compositional reconfiguration of the coexisting chromatin phases is also derived analytically in the SI (Section S4-S5). Our model, thus, predicts that compression shifts the nucleus toward a greater HC fraction (more red in Figure 4a), whereas tension shifts it toward a greater EC fraction (more blue, Figure 4a). The predicted change in chromatin phase balance is magnitude-dependent, reflecting the extent to which applied stress alters nuclear volume and water content. Collectively, these results show that isotropic mechanical loads reorganize chromatin by changing intranuclear hydration and thereby shifting the balance between coexisting HC and EC phases.

#### Compressive load induced chromatin compaction and nuclear volume changes confirm model predictions

To validate the predicted effects of compressive loads, we cultured MDA-MB-231 breast cancer cells and IMR90 fibroblasts on glass substrates and applied compressive loads using a microfabricated confinement device with piston-controlled and displacement-regulated PDMS micro-pillars (Methods, Section S2.7) confining the cells to heights of 10 *μm*, 6 *μm* and 2 *μm* (Figure 4b,c). The 10 *μm* confinement is chosen as a control height which does not deform the nucleus. Conversely, the 2 *μm* confinement severely compresses the nucleus beyond its original height significantly reducing the nuclear volume (Figure 4d for MDA-MB-231 cells, Figure S23 for IMR90 cells). In MDA-MB-231 cells, confinement to 2 *μm* – smaller than the typical nucleus diameter – reduced nuclear volume by approximately 52.2% compared to 10 *μm* conditions. Upon applying a similar volume reduction in the simulations, we observed that the fraction of nuclear space occupied by HC increased by nearly 4.8-fold.

To quantify the nucleus-wide chromatin reorganization observed experimentally, we stained the nuclei with DAPI and measured the fraction of nuclear volume occupied by high-intensity DAPI regions as a proxy for HC. In quantitative agreement with the model predictions, nuclear volume reduction after compression at 2 *μm* increased the HC fraction by nearly 4.6-fold (Figure 4e) compared to 10 *μm* confinement. As further validation, the chromatin condensation parameter (CCP ^18^) increased by nearly 2.05-fold after compression at 2 *μm* confinement relative to 10 *μm* condition (Figure 4f). At intermediate confinement (6 *μm*), nuclear volume, HC fraction and CCP exhibited intermediate values confirming the magnitude-dependence observed via the simulations (Figure S23). Similar trends were also observed in IMR90 cells (Figure S23) for the nuclear volume, HC fraction and CCP. These findings demonstrate that, across two distinct cell lines, compression-driven reduction in nuclear volume is accompanied by increased chromatin compaction and HC enrichment, in agreement with our model predictions.

To determine whether dynamic compressive loading produces similar chromatin reorganization, we reanalyzed the previously published data ^10^ on MDA-MB-231, HT1080 and SV40 human fibroblast cells migrating through constrictive pores (Figure S24a). Across all cell types, confined migration was associated with increased HC markers (Figure S24b-d) and reduced chromatin accessibility (Figure S24e), indicating that compression-induced chromatin compaction is not restricted to a single cell type. These results further support our prediction that both dynamic and steady state compression promote HC formation by reducing nuclear volume and intranuclear hydration.

#### Tensile forces increase chromatin accessibility

Our model predicts that tensile mechanical forces, which increase nuclear volume and promote water influx, favor the formation of EC phase (Figure 4a), which is associated with increase in chromatin accessibility. To confirm this, we examined the ATAC-seq data (collected from GEO repository, accession: GSE207896) comparing chromatin accessibility in tenocytes from mouse tendons maintained under homeostatic tensile load versus tenocytes de-tensed ex-vivo via myosin contractility inhibition blebbistatin treatment ^27^ (Figure 4g). Consistent with model predictions, chromatin accessibility was higher under tensile loading than in the de-tensioned state (Figure 4h). Specifically, compared to homeostatic tenocyte, de-tensioned nuclei exhibited 691 regions of reduced chromatin accessibility and 7 regions with opened chromatin, indicating a shift toward HC upon loss of tension. Thus, sustained tension promotes EC formation and enhances chromatin accessibility, in line with our model predictions (Figure 4a).

Overall, these findings confirm that mechanical forces reorganize chromatin by altering nuclear volume and thereby intranuclear water content, thereby shifting the balance between HC and EC phases and altering chromatin accessibility and compaction. Importantly, these results place mechanical loading within the same hydration-mediated framework that underlies epigenetic and osmotic perturbations described in the previous sections, with nuclear hydration emerging as the common physical pathway linking diverse microenvironmental cues to chromatin reorganization.

### 2.6 Mechanical forces alter the size-scale of heterochromatin domains

We next ask whether, beyond changing nucleus-wide chromatin compaction and the balance between EC and HC phases, mechanical forces also locally regulate the mesoscale sizes of individual HC domains. In Section 2.2, we showed that HC domain size is selected by the balance between epigenetic reaction kinetics and diffusive transport of histone marks, yielding a characteristic size scale described by Eq. 2. Because this scaling relation depends on nuclear hydration and on the compositions of the coexisting chromatin phases, the HC domain size is expected to change under mechanical loading, through the same hydration mediated mechanism established in Section 2.5.

Consistent with this, HC domain size, our simulations show that compressive (tensile) forces increase (decrease) HC domain sizes (Figure 4a). Mechanistically, compression reduces nuclear volume, and the EC hydration which in turn drives a stronger histone diffusive influx resulting in selection of larger HC domains. Tension produces the opposite effect by promoting EC hydration, reducing the histone diffusive influx, thus selecting smaller HC domains. We quantify the mean HC domain size over two simulations (with varying spatial heterogeneities in epigenetic reactions and initial conditions; n ∼ 45 domains) and found that domain size changes systematically with the magnitude of biaxial load (Figure 4i), consistent with the stress-dependent changes in nuclear volume and the intranuclear water content.

To experimentally test these predictions, we reanalyzed the previously published DAPI-stained confocal images of NIH3T3 mouse fibroblast nuclei subject to compressive mechanical load (∼2.5 *μN*) ^8^(Figure 4j). These images reveal DNA-dense HC clusters ^8^ that can be used as a proxy for HC domains. Although these confocal measurements do not fully resolve nanoscale structures accessible by super-resolution methods, they nonetheless exhibit characteristic size distribution that permits comparison with the model predictions.

Consistent with the simulations, compressive loading increased global chromatin compaction, as reflected by an increased fraction of nuclear volume occupied by HC and an approximately 1.13-fold increase in CCP (Figure S25). Moreover, when we quantified the dense HC clusters by measuring the mean radius of the circumscribing ellipse, we observed a ∼ 1.2-fold increase (Figure 4k) after compression, which falls within the range predicted by our simulations (Figure 4i). Thus, the experimental data support the model prediction that compression regulates not only global chromatin state, but also the characteristic size scale of condensed HC domains.

Overall, these results show that mechanical perturbations regulate HC domain size through the same chemo-mechano-osmotic pathway that governs phase balance at the nuclear scale. Compression enlarges HC domains by reducing nuclear hydration and favoring the compact HC phase, whereas tension is predicted to shrink them by promoting the water-rich EC phase. These results, therefore, extend the mechanisms described in Section 2.5 from global chromatin compaction to mesoscale domain architecture, reinforcing the central conclusion that nuclear hydration is the common mediator linking mechanical loading to chromatin phase organization across scales.

### 2.7 Anisotropic forces on the nucleus realign the heterochromatin domains

In preceding sections, we showed that isotropic mechanical loading reorganizes chromatin by changing nuclear volume, hydration and chromatin phase balance. We next ask how directionally constrained mechanical loads, such as uniaxial compression and shear, influence chromatin reorganization – as is observed in a range of biological contexts such as extravasation and intravasation ^2^. Unlike isotropic loading, anisotropic stresses can not only modify phase balance through osmotic effects, but also impose a directional bias in domain morphology and orientation. We therefore use the same chemo-mechano-osmotic framework to examine how uniaxial and shear stresses reorganize heterochromatin domains in space.

#### Uniaxial forces result in elongation of HC domains along the principal strain orientation

To examine the effects of anisotropic loading, we first apply uniaxial tension or compression to the nucleus (Figure 5a). As in the biaxial loading case, these perturbations can alter nuclear volume and therefore drive water influx or efflux across nuclear envelope in tension or compression respectively. Consequently, uniaxial compression (tension) shifts the chromatin landscape water-poor HC (water-rich EC) phase. Consistent with the isotropic results, our simulations predict that uniaxial compression increases overall HC content and HC domain size (Figure 5a), while uniaxial tension favors EC and yields smaller HC domains (Figure 5a).

**Figure 5:**
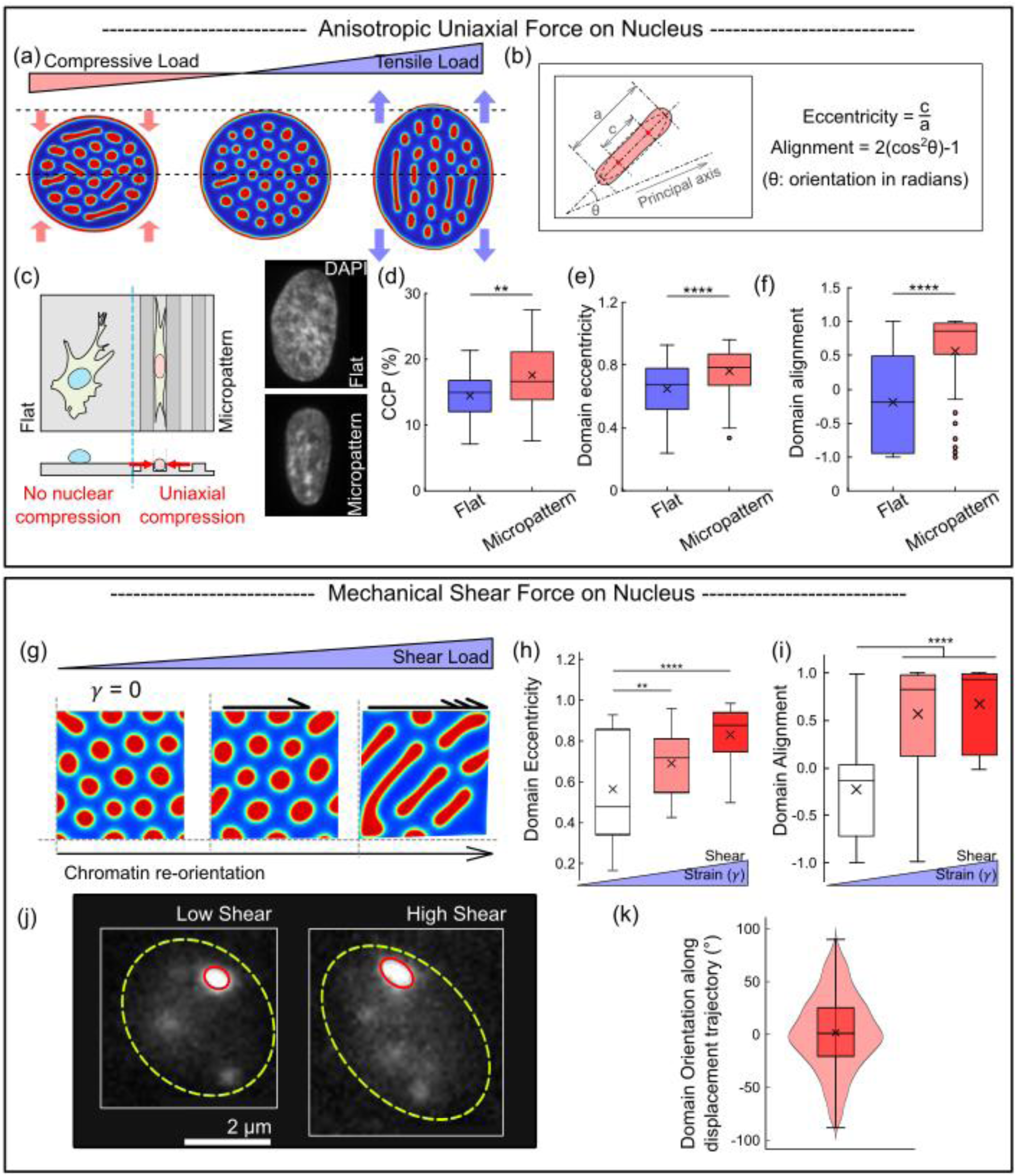
Anisotropic mechanical forces can reorient heterochromatin domains while spatially reorganizing chromatin. (a) Numerical model shows that uniaxial tensile and compressive forces alter the chromatin organization by not only changing the size of the heterochromatin domains, but also introducing a directional bias by forming lamellar domains. The orientation of the domains depends on the type of mechanical load applied. Under tension the domains reorient along the applied load while in compression the domains orient perpendicular to the compression force. (b) The elongation and orientation of the heterochromatin domains is quantified respectively via eccentricity (1: straight line, 0: circle) and the alignment parameter (1: along, 0: perpendicular) of an ellipse circumscribing the domain, as shown in the schematic. (c) Schematic depicting hMSCs cultured on flat and micropatterned glass substrates with ridges narrower than nucleus diameter, with DAPI stained nucleus on right. The chromatin reorganization after anisotropic uniaxial compressive loads and is accompanied by (d) an increase in CCP (n = 29 and 30 nuclei on flat and micropatterned substrates respectively; unpaired, two-tailed t-test, p=0.008), with heterochromatin domains showing (e) an increased eccentricity (unpaired, two-tailed t-test, p=0.01) and (f) a preferential realignment perpendicular to the anisotropic compressive force (unpaired, two-tailed t-test, p=1e-9). (g) Application of anisotropic shear loads on the region in the interior of the nucleus results in a reorientation of heterochromatin domains. Simulations predict that application of shear loads will result in (h) increased domain eccentricity as well as (i) increased domain realignment in a magnitude-dependent manner. (j) Confocal images showing fluorescently labelled Swi6-GFP (yeast orthologue of HP1) in S. pombe nucleus ^11^ with the heterochromatin domain (bright intensity) in the vicinity of spindle pole body (SPB) outlined in red. The microtubule dynamics apply varying levels of shear loads on the nucleus, and specifically on the chromatin region in the SPB vicinity. Quantifying the domain morphologies, we observe (k) alignment of the domains preferentially along the SPB displacement due to shear forces from the nucleus (n=11973 domains observed over 164 cells).

Beyond these hydration-mediated changes in phase balance, our model predicts that anisotropic loading introduces a directional bias in chromatin organization (Figure 5a). In the absence of external forces, HC domains are approximately circular (Figure 5a, center panel). However, under uniaxial loads the domains preferentially adopt elongated, lamellar morphologies (Figure 5a). We quantify this transition using the eccentricity (ratio of interfocal distance to the length of major axis; 0: circle, 1: straight line, Figure 5b) and aspect ratio (ratio of major to minor axes; 1 indicates a circle) of an ellipse circumscribing each domain (Figure 5b), both of which increase with the magnitude of the applied uniaxial load. Thus, anisotropic loading does not simply change the amount of HC present in the nucleus; it also reshapes the domains themselves.

The simulations further predict a load-dependent orientation adopted by the lamellar HC domains. Under uniaxial tension, the lamellar domains align parallel to the tensile forces applied (i.e. along the principal strain direction). Under uniaxial compression (Figure 5a, left panel) they align perpendicular to the applied compression, which is again along the principal strain direction. Quantifying orientation via the alignment parameter (2(*cos θ*)^2^ − 1, *θ*: angle between domain major axis and principal direction, Figure 5b), shows that increasing uniaxial load drives progressively stronger domain alignment with respect to each other (alignment parameter close to 1) as well as to the principal direction (*θ* → 0). Mechanistically, this collective realignment reduces the strain energy associated with deforming the relatively stiffer HC phase and therefore represents a mechanically favored chromatin configuration under anisotropic stress.

#### Theoretically predicted chromatin realignment is observed experimentally

We next tested these predictions experimentally by culturing hMSCs on micropatterned substrates with parallel ridges of width 400 nm (Figure 5c), a geometry that constrains the nucleus anisotropically. Relative to cells on flat substrates, nuclei on micropatterned ridges showed increased chromatin condensation as quantified via CCP (Figure 5d) consistent with the prediction that compressive loading promotes HC enrichment.

We then quantified the morphology and orientation of HC domains from DAPI stained images (Figure 5b). On micropatterned substrates, HC domains exhibited significantly higher eccentricity (Figure 5e) indicating the formation of more lamellar structures, as predicted by our model. In addition, the domains displayed preferential alignment along the micropatterned ridges (Figure 5f, value of 1 denotes the alignment along the ridges), which is perpendicular to the direction of compression imposed by the ridge walls and matches the model prediction for uniaxial compression. This realignment was reflected not only in an increased mean alignment parameter, but also by a narrower alignment distribution, indicating a more coherent orientation across domains (Figure 5f). Together, these results support our predictions that anisotropic compressive loads reorganize chromatin not only by increasing HC compaction, but also by elongating and reorienting HC domains along a mechanically favored direction.

#### Volume-preserving shear loads realign HC domains

We next isolate the role of anisotropic stress from the hydration-mediated effects associated with nuclear volume change by applying volume-preserving shear stresses. Unlike anisotropic compression or tension, shear loads can deform the nucleus while preserving its volume.

Starting with a steady state chromatin organization in the nucleus, we applied volume preserving shear strains on an interior nucleus region (Figure 5g). Our simulations confirmed that the shear forces did not alter the nuclear volume (Figure S26a), the overall water content in the nucleus (Figure S26a) or the fraction of total area occupied by the HC phase (HC area/nucleus area, Figure S26a). Despite this, the morphology and orientation of HC domains change substantially. Under increasing shear, the HC domains become progressively elongated (Figure 5g), as reflected by the increase in domain eccentricity (Figure 5h) and aspect ratio (Figure S26c). A theoretical analysis of a simplified HC domain embedded in EC confirms the physical basis of this result: shear stress does not alter nuclear volume or the composition of the coexisting HC and EC phases, but it does generate deviatoric strain (Section S9.2, Eq S80, last equation in red) within the stiffer HC phase – which changes its shape but not its volume (distortion without dilation).

Additionally, shear loads also produce a strong magnitude-dependent HC domain realignment (Figure 5i). Quantifying the angle between the major axis of each domain and the first principal strain direction (45° for this loading condition) shows that in the absence of shear, domain orientations are broadly distributed, whereas increasing shear causes the domain alignment parameter to approach 1. Moreover, the distribution of alignment parameter of all HC domains becomes narrower in a shear magnitude-dependent fashion (Figure 5i). To confirm the local realignment of HC with principal direction, we overlay the principal directions denoted via arrows above the chromatin organization under the high shear condition (Figure S26b). We observe that the orientation of HC domains at each grid point is exactly along the first principal direction.

To test these predictions in a biological setting, we reanalyzed a previously reported chromatin reorganization in S. pombe nuclei experiencing in-vivo shear due to the microtubule spindle-pole body (SPB) coupling (Figure 5j) ^11^. The orientation of HC domains in the SPB-vicinity is quantified as the angle between the major axis of HC domains in SPB vicinity and the trajectory of domain deformation as principal-strain direction readout (Figure 5k). The quantification confirms that the mean orientation of the heterochromatin domains is 1.5° with respect to the principal deformation direction. Moreover, about 45% of the domains are oriented within ±20° of the principal deformation direction.

Altogether, these results demonstrate that anisotropic mechanical forces reorganize chromatin in a direction-dependent manner. Uniaxial stresses combine hydration-mediated shifts in HC/EC balance with directional changes in domain morphology and alignment, whereas volume-preserving shear isolates the latter effect by reorienting HC domains without changing nuclear water content or phase abundance. Taken together, these observations establish a unified framework in which chemo-mechano-osmotic cues jointly regulate chromatin reorganization – from macroscale phase balance to mesoscale HC domain morphology.

## 3 DISCUSSION

The compartmentalized organization of chromatin observed across super-resolution imaging modalities ^17,28,29^ has often been interpreted through the lens of liquid-liquid phase separation (LLPS) between transcriptionally distinct chromatin states ^30,31^. Existing models have provided important insights into chromatin folding and organization across states ^12,32,33^, but they do not fully account for a central feature of chromatin at the mesoscale – its solid-like mechanical response ^30,34–36^. Here, we develop a non-equilibrium continuum framework that incorporates chromatin elasticity, nucleoplasmic water transport and epigenetic reaction-diffusion kinetics to explain how chromatin organization emerges and reorganizes under epigenetic, osmotic and mechanical perturbations.

A central hypothesis of this work is that nuclear hydration provides the physical link between mesoscale chromatin phase behavior and macroscale nuclear mechanics. In our framework, chromatin segregates into coexisting water-rich euchromatic and water-poor heterochromatic phases, but this segregation is not governed by epigenetic regulation alone. Because the two phases differ in hydration, phase separation necessarily generates differential swelling and hence an elastic mismatch between the two phases. The resulting strain energy opposes complete segregation, reshapes the free-energy landscape, and shifts the steady-state compositions of the coexisting phases. Thus, elasticity is not merely an added material property of chromatin; it directly regulates the composition, morphology, and stability of mesoscale chromatin domains.

This perspective also clarifies how our framework relates to liquid-like descriptions of chromatin organization. At short length and time scales, chromatin can exhibit liquid-like behavior driven by thermal fluctuations and constrained sub-diffusion ^37^. At the mesoscale relevant to HC domains, however, multiple experiments point to solid-like behavior, including resistance to deformation, distinct phase-dependent stiffness, and limited fluorescence recovery after photobleaching ^19,30,34–36,38,39^. Our results therefore argue that mesoscale chromatin reorganization under the perturbations studied here cannot be understood without accounting for elasticity. Rather than framing chromatin organization as strictly liquid-liquid or strictly solid-solid, our model identifies the regime in which elastic stresses and hydration-dependent swelling are essential determinants of phase organization and its mechanical consequences.

A second key result is that the characteristic size of heterochromatin domains emerges from a non-equilibrium balance between diffusion-like transport and epigenetic interconversion. In the absence of reactions, diffusion would drive continued coarsening of compact heterochromatin domains. In contrast, our model shows that acetylation- and methylation-mediated interconversion competes with diffusive influx at the HC-EC interface and arrests growth at a finite domain size, which scales via Eq 2. Because the coexisting phase compositions entering this flux balance are themselves set by hydration and elastic stress, the selected domain size is not purely reaction-diffusion controlled, but intrinsically chemo-mechano-osmotic. This prediction is supported by pharmacologic perturbations of histone acetylation and methylation, which shift heterochromatin domain size in the directions predicted by the model (Figure 6).

**Figure 6:**
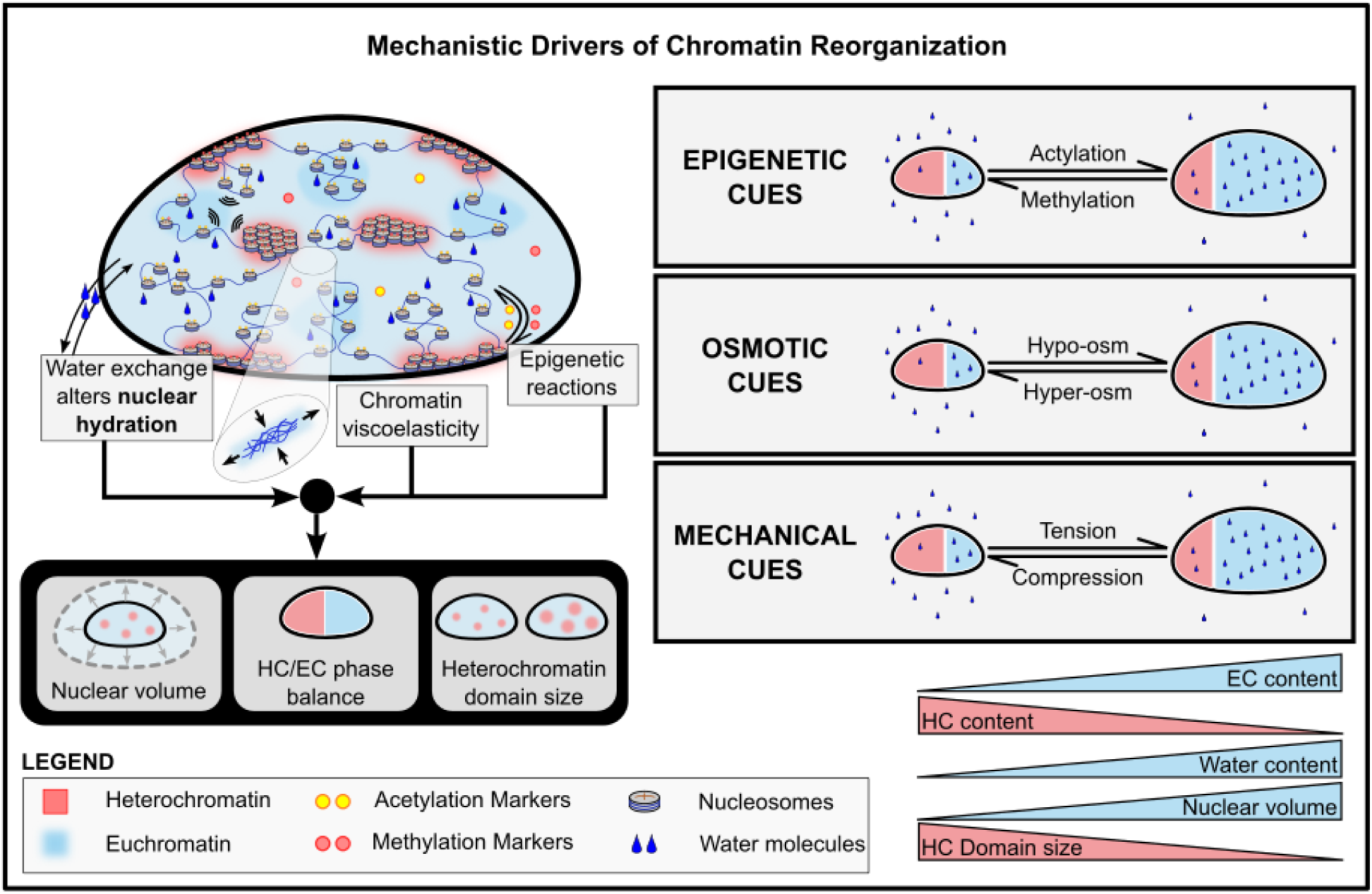
Mechanistic framework for chromatin reorganization under chemo-mechano-osmotic perturbations. (Left) Schematic of the proposed model showing the interplay between nuclear hydration, epigenetic histone modifications and chromatin viscoelasticity. Chromatin is treated as a viscoelastic polymer that deforms while storing mechanical strain energy. Epigenetic reactions are governed by the availability of epigenetic factors such as HDACs/HATs/HMTs/HDMs, while the nuclear hydration is dictated by the water exchange between the nucleus and the cytosol. Together, these factors regulate the chromatin phase balance in terms of the chromatin phase balance (HC versus EC), the mean size of the HC domains as well as the nuclear volume and stiffness. (Right) External signals including epigenetic (change in rates of histone acetylation and methylation), osmotic (hypo- and hyper-osmotic) and mechanical (tensile, compressive and shear) perturbations of the nucleus are summarized as predicted by our model.

Our mesoscale chromatin modeling framework explains how distinct classes of perturbation produce apparently diverse chromatin phenotypes through a common mechanism. Epigenetic perturbations alter the relative abundance of the water-rich EC and water-poor HC phases, thereby driving water influx or efflux across the nuclear envelope and changing nuclear volume and stiffness. Osmotic perturbations act more directly on the same variable by changing water exchange with the cytosol, and accordingly reorganize chromatin compaction and domain size. Isotropic mechanical loading accesses this pathway through deformation: compression reduces nuclear volume and hydration, favoring HC enrichment and larger domains, whereas tension promotes water influx, EC enrichment, and greater chromatin accessibility. Across these perturbation classes, nuclear hydration emerges as the common mediator linking external cues to chromatin phase balance and to the macroscopic mechanical state of the nucleus. At the same time, our results show that not all mechanically induced chromatin reorganization is reducible to changes in phase balance alone. Anisotropic forces introduce a directional bias in chromatin organization. Under uniaxial loading, domain size changes are accompanied by elongation and collective alignment of heterochromatin domains along principal strain directions. Under volume-preserving shear, by contrast, the overall water content and HC-EC phase balance remain essentially unchanged, yet the domains still become elongated and reoriented. This distinction is important because it shows that hydration-mediated changes in phase composition and strain-mediated changes in domain shape and orientation are coupled but separable consequences of mechanical loading. In this sense, isotropic and anisotropic loads probe different aspects of the same chemo-mechano-osmotic framework.

More broadly, the coupling between nuclear mechanics and chromatin organization predicted here is consistent with the emerging evidence of the nucleus as an active mechanosensor ^2,23^. Moreover, during T-cell activation, nuclear enlargement is accompanied by a reduction in DNA nanodomain size, with nuclear area inversely correlated with nanodomain size ^40^ (relevant results reproduced in Figure S27, SI). This observation is consistent with our prediction that increased nuclear hydration favors a more open chromatin organization. Studies of B-cell activation have similarly shown that histone acetylation accompanies chromatin decompaction and reorganization of nuclear architecture ^41^. By linking chromatin phase balance to nuclear volume, stiffness, accessibility, and domain morphology, our model provides a mechanistic basis for how environmental cues may be integrated at the level of nuclear architecture. It offers a physical interpretation for observations that cyclic loading, confinement, and mechano-osmotic perturbations remodel chromatin accessibility and condensation in epithelial, mesenchymal, and stem cell contexts ^42–44^. Within this framework, chromatin is not simply a passive substrate of gene regulation, but a mechanically responsive material system whose hydration state helps determine how the nucleus adapts to its microenvironment.

The limitations of the present model point to natural directions for future work. Although we incorporate chromatin-lamina interactions that generate lamina-associated domains, the current framework does not resolve epigenetic-state-specific modes of chromatin anchoring at the nuclear periphery ^45^ or the active mechanical remodeling of the lamina itself ^46^. In addition, while the model captures time-dependent chromatin reorganization under the perturbations considered here, it does not yet address how temporally varying forces couple to slower biochemical programs, transcriptional feedback, or cell-fate decisions. Extending the framework to include lamina mechanics, explicit mechanosensitive transport of epigenetic regulators, and longer-timescale transcriptional responses should help connect mesoscale chromatin physics more directly to nuclear function.

Overall, this work establishes a unified non-equilibrium chemo-mechano-osmotic framework for chromatin organization. By coupling histone-mark-dependent interactions, hydration-driven deformation, and epigenetic reaction-diffusion kinetics, the model explains how chromatin domains emerge, acquire a characteristic size, reorganize under environmental perturbation, and contribute to the emergent mechanics of the nucleus. These results position nuclear hydration as a central physical variable through which biochemical and biophysical cues are integrated to regulate chromatin organization across scales.

## 4 METHODS

### Overview of mathematical model of mesoscale chromatin reorganization

To capture the nucleus-wide mesoscale spatiotemporal organization of chromatin, we model three nuclear constituents: the nucleoplasm, and the two main epigenetic states of histones – acetylated and methylated. At each point in the nucleus, we define the local volume fraction (i.e., volume of a constituent per unit volume of the infinitesimal point prior to any deformations) denoted as *φ_i_*(***x***, *t*) with *i* = n (nucleoplasm), a, m (acetylated or methylated histones). Note that volume fractions of the three nuclear constituents at any infinitesimal point will add up to unity i.e., *φ_a_* + *φ_m_* + *φ_n_* = 1 (see SI, Section S1.1 for derivation). In the HC phase, the histones are predominantly methylated (*φ_m_* > *φ_a_*), while in the EC phase they are prominently acetylated (*φ_a_* > *φ_m_*). Additionally, variable *φ_d_*(= *φ_m_* − *φ_a_*) acts as an epigenetic order parameter measuring the local preference for epigenetic state and is positive (negative) for HC (EC).

We consider a nucleoplasm-devoid nucleus as a reference configuration, with respect to which all mechanical deformations, strains and free-energy are measured. With space-occupying water molecules present, as well as due to chromatin-chromatin interactions, chromatin deforms with changes to its shape and volume – which we characterize by the deformation gradient tensor ***F***(***x***, *t*), a generalization of the stretch tensor applied to large deformations. At any point in the nucleus, the ratio of volume at time *t* with respect to that in reference configuration, is given as *J*(***x***, *t*) = det ***F***, and is related to the local nucleoplasm content as 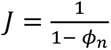.

In the following, we express the free energy of the nucleus in terms of these variables, which allows us to determine chromatin evolution under chemo-mechanical cues.

### The energy landscape governing chromatin organization

At any point in the nucleus, we can define the free energy of chromatin organization per unit reference volume in terms of the variables ***F***, *φ_n_*, and *φ_d_*, as

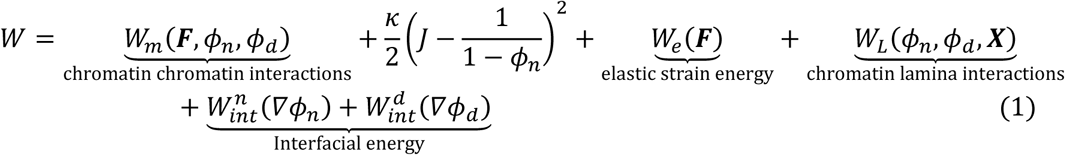

Here, each term denotes a distinct contribution to the free energy density, as follows:

i. The first term is Flory-Huggins type free-energy density and denotes *chromatin-chromatin interactions* capturing the interactions between differentially marked nucleosomes and gives rise to the double-well energy landscape with the thermodynamically preferred HC and EC phases. These interactions may be mediated by cross-bridging of methylated histones via HP1*α* ^47^, internucleosome ionic interactions, or transcription-induced entropic variations in HC and EC phases. Mobile ions and electrostatic fields are not resolved as independent fields at the mesoscale considered here; their rapidly equilibrating contributions are incorporated into this effective interaction free energy (Section S1.5, SI).
ii. While change *φ_d_* only alters the local epigenetic state, i.e. number of acetylated versus methylated histones, changes in the water content *φ_n_* alters the number of space occupying water molecules and thus the local volume. The second term enforces this constraint enabling the water-rich EC to swell more than the water-poor HC.
iii. The third term incorporates the additional elastic strain energy cost arising from deformations in the solid-like chromatin polymer as it swells ^48^. We capture the elastic strain energy via the neo-Hookean model which approximates the entropic contributions to the change in the free energy due to mechanical deformation (shear and swelling) of chromatin from the reference configuration ^49^. The shear modulus, which quantifies chromatin stiffness, is a measure of the cross-linking of polymer. In accordance with observations from nuclear elastography ^20^, the shear modulus depends on the epigenetic state, such that heterochromatin is stiffer than euchromatin.
iv. Next, the chromatin-lamina interactions capture the role of inner nuclear membrane proteins such as LAP2*β*, emerin, and MAN1 ^32^ in anchoring heterochromatin to the nuclear lamina. We consider these interactions to be strongly localized to the nuclear lamina such that the strength of such interactions decays exponentially away from the lamina ^15,16^. Since the chromatin anchoring at the lamina are preferentially enriched in methylated histones associated with heterochromatin ^50^, the chromatin-lamina interactions are captured specifically towards HC phase.
v. The separation of the dissimilar chromatin phases leads to the formation of an interface between them. Since methylated histones preferentially interact with other methylated histones rather than that with acetylated histones, the presence of heterochromatin-euchromatin interface incurs an additional energy cost. The last two terms explicitly account for this interfacial energy cost penalizing interface formation.

The free-energy landscape defined by Eq (1) (described explicitly in SI Section S1.4) produces two favorable phases of chromatin – a water-rich, prominently acetylated EC phase, and a water-poor prominently methylated, compacted HC phase. At each point in the nucleus, the local nucleoplasm fraction *φ_n_* and the epigenetic order parameter *φ_d_* evolve towards one of these stable phases. This evolution is propelled by the driving forces originating from the gradients in the energy landscape called chemical potential and defined as 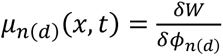. Here, the operator *δ* denotes the change in free energy density due to change in volume fraction of nucleoplasm *φ_n_* or the order parameter *φ_d_*, respectively defining the chemical potential of nucleoplasm (*μ_n_*) or that of the order parameter (*μ_d_*). The evolution occurs such that all physical forces within the nucleus are balanced at all times, which gives, *∇_k_s_iK_* = 0, where *s_iK_* = *δW*/*δF_iK_* denotes the nominal stress (defined as the force in the current configuration per unit reference area).

### Kinetic evolution of nucleoplasm and epigenetic marks

The spatiotemporal evolution of nucleoplasm *φ_n_*(***x***, *t*) is governed by its diffusion along the gradients in nucleoplasm chemical potential as

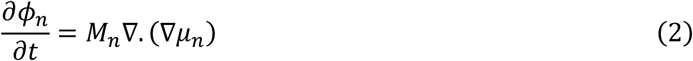

where *M_n_* is the mobility of nucleoplasm in the nucleus and ∇ denotes spatial gradients. As discussed in Section S1.2, this evolution is conservative, the total amount of nucleoplasm in the nucleus is conserved unless water is explicitly allowed to enter or exit the nucleus and is governed by the energy landscape.

In contrast, the evolution of the epigenetic order parameter *φ_d_* is non-conservative (Section S1.2) because epigenetic marks can interconvert through reactions catalyzed by epigenetic factors such as histone deacetylase (HDAC), histone methyltransferase (HMT), histone acetyltransferase (HAT), and histone demethylase (HDM, Figure 1d, Eq. S11). Since like-marked histones prefer to accumulate, the conversion of acetylated to methylated histones is energetically more favorable in the HC region than in the EC region, and vice versa. Consequently, the effective reaction rates depend on (i) availability of epigenetic factors, and (ii) the epigenetic marks on the neighboring histones. In the SI Section S1.7, we theoretically describe how such spatially heterogeneous, neighborhood-dependent reaction kinetics can be effectively represented as a combination of conservative diffusion-like evolution of epigenetic marks and non-conservative reaction-like dynamics –

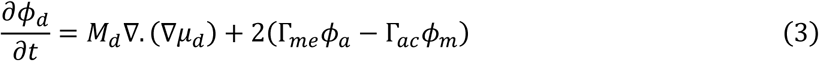

where, *M_d_* denotes the effective mobility of epigenetic marks due to the diffusion-like evolution and depends on neighbourhood-dependent reaction rates. Moreover, the spatial diffusion-like evolution of epigenetic marks can occur along the chromatin polymer, referred to as cis-diffusion (Figure S1), or between histones that are spatially proximal but not sequentially adjacent along the polymer chain, referred to as trans-diffusion (Figure S1). Both cis- and trans-diffusion of epigenetic marks are effectively captured in our model through the diffusive kinetic term in Eq. S11. Thus, the overall evolution of epigenetic marks arises from a combination of non-conservative reaction kinetics and conservative diffusion-like dynamics, as described by Eq. S11 (Figure 1d).

The combined energetic and kinetic contributions yield a fully coupled reaction-diffusion-mechanical system that governs chromatin organization over space and time (Section S1.1-1.5). Prior to solving these, we rescale the governing equations with respect to the intrinsic length and timescales of the system, as described in Section S1.6. We solve these equations numerically, subjected to initial and boundary conditions described in Section S1.8, to explore how the intranuclear chromatin architecture adapts to mechanical, osmotic and biochemical perturbations arising from the cellular microenvironment.

### Cell culture, immunostaining and confocal imaging

Bone marrow-derived human mesenchymal stem cells (hMSCs; Lonza, PT-2501) were cultured in low-glucose DMEM supplemented with 10% fetal bovine serum, 1% antibiotic–antimycotic and 1% GlutaMAX and seeded at a density of 2000cells/cm². For immunostaining cells were fixed, permeabilized then blocked with bovine serum albumin (BSA) and immunolabelled for histone H2A and counterstained DAPI. Confocal z-stacks were acquired using a 63× oil-immersion objective and images were analysed in Fiji to quantify nuclear morphology and volume. Detailed cell-culture, immunostaining and image analysis procedures are provided in the Supplementary Section S2.

### Chromatin condensation analysis

Chromatin condensation parameter (CCP) was quantified from DAPI stained nucleus using a MATLAB script, as previously described ^18^. In brief, individual nuclei were analysed using a Sobel gradient-based edge-detection algorithm that identifies local intensity transitions within the nuclear DAPI signal. CCP was calculated from the density of detected chromatin edges within the nuclear area, with higher CCP values corresponding to a more condensed chromatin organization. Additional details of image preprocessing and CCP calculation are provided in the Supplementary Section S2.

### Atomic force microscopy

Nuclear stiffness was measured in live cells by atomic force microscopy (AFM). The AFM probe was positioned above the central nuclear region using phase contrast optics and force indentation curves were acquired under controlled indentation conditions. Curves were fit in AtomicJ using a Hertzian contact model for a pyramidal indenter, and the apparent nuclear stiffness was reported as the Young’s modulus obtained from the fitted force-indentation curves. Details of cantilever properties, calibration and indentation parameters are provided in the Supplementary Section S2.

### Super-resolution imaging of chromatin organization

Nanoscale chromatin organization was visualized by stochastic optical reconstruction microscopy (STORM) of immunolabelled histone H2A using an Oxford NanoImager. Alexa Fluor 647-labelled samples were imaged under photoswitching conditions and single-molecule localizations were reconstructed to generate super-resolution maps of H2A organization. Localization data were subsequently analysed using custom MATLAB codes to quantify nanoscale chromatin organization. Detailed imaging conditions and analysis procedures are described in the Supplementary Section S2.

### FLIM–FRET analysis of live-cell chromatin compaction

Live-cell chromatin compaction was independently quantified in HeLa cells co-expressing the histone FRET pair H2B–eGFP and H2B–mCherry. FLIM–FRET measurements were performed under iso-osmotic (control), hypo-osmotic, or hyper-osmotic conditions. Chromatin decompaction was induced by hypotonic treatment, in which imaging medium was diluted with distilled water to a final ratio of 35:65 (v/v), whereas chromatin compaction was induced by supplementing the imaging medium with 160 mM sucrose. Fluorescence-lifetime data were analysed using a phasor-based approach to generate nuclear FRET-efficiency maps, in which regions of higher FRET efficiency correspond to more compact chromatin. These maps were used to quantify the fraction of the nucleus occupied by compact heterochromatin and the corresponding heterochromatin-domain size distributions. Complete acquisition and analysis procedures are provided in the Supplementary Section S2.

### Mechanical confinement of cells

To examine chromatin and nuclear responses to mechanical deformation, MDA-MB-231 breast cancer cells and IMR90 fibroblasts were confined to defined heights using non-adhesive microfabricated PDMS spacer coverslips. Confined cells were imaged live by Airyscan confocal microscopy, and z-stacks were used to characterize deformation-associated changes in nuclear and chromatin organization. Fabrication of the confinement devices, confinement conditions and imaging procedures are described in details in the Supplementary Section S2.

### Micropatterned substrates

For experiments examining responses to anisotropic microenvironmental geometry, hMSCs were cultured on fibronectin coated PDMS substrates containing parallel microgrooves or on flat PDMS controls. Cells were allowed to attach and align along the microgrooves before subsequent experimental analysis. Detailed PDMS fabrication, surface treatment and cell seeding are provided in the Supplementary Section S2.

## DATA AVAILABILITY STATEMENT

The authors declare that the data supporting the findings of this study are available within the paper and its supplementary information files. The data generated in this study are provided in the Source Data file.

## ACKNOWLEDGEMENT

This work was supported by NIH Award U54CA261694 (V.B.S.); NSF CEMB Grant CMMI-154857 (V.B.S.); NSF Grant DMS-2347834 (V.B.S.); National Institute of Biomedical Imaging and Bioengineering (NIBIB) Awards R01EB017753 (V.B.S.) and R01EB030876 (V.B.S.); National Institute of General Medical Sciences award R01GM155943 (V.B.S.) and National Institute of Diabetes and Digestive and Kidney Diseases grant R01DK144619 (V.B.S.). We gratefully acknowledge the valuable feedback and suggestions from Dr. G.K. Shivashankar. We are also grateful to Dr. Jan Lammerding for sharing the data on changes in epigenetic markers after confined migration.

