## Supplementary Information for "Osmotic Coupling of Mechanics and Epigenetics Regulates Chromatin Phase Organization"

### S1 Extended description of mathematical model for nucleus-wide spatial chromatin reorganization

#### *S1.1 Model for Solid-solid Chromatin Phase Separation*

To investigate the effect of chemo-mechanical stresses upon chromatin organization, we develop a mathematical model for phase separation of viscoelastic solid-like chromatin by coupling chromatin and water transport with associated local structural reconfigurations within the nucleus. To track the local structural reconfigurations, we define a theoretical reference state that is free of nucleoplasm[1] with reference coordinates $\boldsymbol{X}$. The current state of the nucleus at any time $t$ can be written as material coordinates $\boldsymbol{x}(\boldsymbol{X},t)$. The deformation of the nucleus in the current configuration relative to the reference configuration is tracked by the deformation gradient $F_{iK}(\boldsymbol{X},t)$,

$$F_{iK}(\boldsymbol{X},t) = \frac{\partial x_i(\boldsymbol{X},t)}{\partial X_K} = \nabla_K x_i, \qquad J(\boldsymbol{X},t) = \det \boldsymbol{F}(\boldsymbol{X},t) \tag{S1}$$

Here, $J(\boldsymbol{X},t)$ denotes the local volumetric swelling, i.e. volume of an infinitesimal region at a point $\boldsymbol{X}$ and time $t$ relative to its volume at the reference state.

At any material point $\boldsymbol{x}(\boldsymbol{X},t)$, at time $t$, consider an infinitesimal observation window of with a volume (in current configuration) $V(\boldsymbol{X},t)$. The three nuclear constituents – nucleoplasm, acetylated and methylated histones are present in this observation window, with respective volumes (in current configuration) $V_n(\boldsymbol{X},t), V_a(\boldsymbol{X},t)$, and $V_m(\boldsymbol{X},t)$. Then,

$$V_n + V_a + V_m = V,$$

$$\Rightarrow \frac{V_n}{V} + \frac{V_a}{V} + \frac{V_m}{V} = 1$$

The ratio of the current volumes of the individual components to the total current volume of the observation window is defined as the material volume fraction (i.e., volume of a constituent per unit current volume of an infinitesimal point in the nucleus) denoted as $\phi_i(\boldsymbol{X},t)$ with $i = a, m, n$. Hence, note that the volume fractions of the three nuclear constituents at any infinitesimal point, when measured in the reference configuration, will add up to unity, such that, $\phi_a + \phi_m + \phi_n = 1$.

The volume fraction can also be defined with respect to the reference configuration (i.e. volume per unit reference volume) for nucleoplasm $\Phi_n(\boldsymbol{X},t)$, and acetylated $\Phi_a(\boldsymbol{X},t)$ and methylated $\Phi_m(\boldsymbol{X},t)$, where the corresponding material and reference volume fractions are related via the equation $\Phi = \phi J$. Thus, the physical state of any material point in the nucleus can be fully described in terms of the three independent field variables $J, \Phi_m$, and $\Phi_a$.

Equivalently, we can redefine the physical state in terms of field variables $J, \Phi_n$, and $\Phi_d$, which respectively denote the deformation gradient (Eq (S1)), the reference volume fraction of the nucleoplasm and the difference between the reference volume fractions of methylated and acetylated histones ($\Phi_d = \Phi_m - \Phi_a$). While equivalent, the latter description permits a natural

[1] The nucleoplasm-free reference state of the nucleus is a hypothetical one where it is devoid of nucleus, and via Eq S1, $J = 1$. A reference state need not be a real, physically observable state (1. Larché, F. and J.W. Cahn, *A linear theory of thermochemical equilibrium of solids under stress.* Acta metallurgica, 1973. **21**(8): p. 1051-1063.)

definition of the movement of the mobile species in the nucleus i.e. water in the nucleoplasm in terms of $\Phi_n$ and epigenetic marks on the histones in terms of preference of methyl marks over acetyl marks $\Phi_d$. As discussed at length in the main text $\Phi_d$ acts as an order parameter such that it is positive (negative) in heterochromatin (euchromatin) region.

The free energy per unit reference volume at any point in the nucleus can thus be expressed as $W(F, \Phi_n, \Phi_d, \boldsymbol{\nabla\Phi_n}, \boldsymbol{\nabla\Phi_d})$. Here the effects of the spatial gradient of the volume fractions with respect to the reference configurations are incorporated to capture the interface between the two phases (as discussed further in Section S1.4, Eq S17). The presence of osmotic and mechanical forces acting on the nucleus (at its outer surface) from its environment can also alter the chromatin distribution, thereby expending thermodynamic work on the nucleus. The mechanical forces acting on the outer surface of the nucleus are captured as a traction vector $\boldsymbol{t}$, doing a mechanical work expressed as $\boldsymbol{t} \cdot \boldsymbol{x}$. Similarly, osmotic forces which drive the exchange of water between the nucleus and its surroundings is captured as extranuclear chemical potential of water $\mu_n^{ext}$. The work done by the external chemical potential of water at the boundary is $\mu_n^{ext} I^n$ where $I^n$ gives the volume of water entering per unit reference area normal to the boundary. The total free energy of the nucleus including the work done on it can be written as,

$$\Pi[\boldsymbol{x}, \Phi_n, \Phi_d] = \int_V W(F, \Phi_n, \Phi_d, \boldsymbol{\nabla\Phi_n}, \boldsymbol{\nabla\Phi_d})\, dV - \underbrace{\int_{\partial V_t} t_i x_i dA}_{\text{Work done by mechanical forces}} - \underbrace{\int_{\partial V_p} \mu_n^{ext} I^n dA}_{\text{Work done by osmotic forces}} \tag{S2}$$

where $\partial V_t$ is the reference surface where external traction acts, while $\partial V_p$ denotes the reference surface where the chemical potential of water acts. At the steady state, the total fre e energy of the nucleus is minimized. Minimizing the total free energy ($\delta\Pi = 0$) using variational principles, and subsequently integrating by parts and applying the divergence theorem gives,

$$\begin{aligned} 0 = &\int_V -\nabla_K \overbrace{\left(\frac{\partial W}{\partial F_{iK}}\right)}^{s_{iK}} \delta u_i dV + \int_V \overbrace{\left[\frac{\partial W}{\partial \Phi_n} - \nabla_K \left(\frac{\partial W}{\partial \nabla_K \Phi_n}\right)\right]}^{\mu_n} \delta\Phi_n dV + \int_{\partial V} \frac{\partial W}{\partial \nabla_K \Phi_n} \widehat{N}_K \delta\Phi_n dA \\ &+ \int_V \overbrace{\left[\frac{\partial W}{\partial \Phi_d} - \nabla_K \left(\frac{\partial W}{\partial \nabla_K \Phi_d}\right)\right]}^{\mu_d} \delta\Phi_d dV + \int_{\partial V} \frac{\partial W}{\partial \nabla_K \Phi_d} \widehat{N}_K \delta\Phi_d dA \\ &+ \int_{\partial V_t} \left(\frac{\partial W}{\partial F_{iK}} \widehat{N}_K - t_i\right) \delta u_i dA - \int_{\partial V_p} \mu_n^{ext} \delta I^n dA \end{aligned} \tag{S3}$$

For this to be true for any arbitrary changes in $\boldsymbol{u}$, $\Phi_n$, and $\Phi_d$, the following equations must hold true at all points in the nucleus,

$$\begin{aligned} \nabla_K s_{iK} &= 0, \\ \mu_n(\boldsymbol{x}, t) &= \frac{\partial W}{\partial \Phi_n} - \nabla_K \left(\frac{\partial W}{\partial \nabla_K \Phi_n}\right), \\ \mu_d(\boldsymbol{x}, t) &= \frac{\partial W}{\partial \Phi_d} - \nabla_K \left(\frac{\partial W}{\partial \nabla_K \Phi_d}\right) \end{aligned} \tag{S4}$$

Here, $s_{iK} = \frac{\partial W}{\partial F_{iK}}$ denotes the nominal stress (also called the first Piola-Kirchoff stress) such that the first equation in Eq (S4) accounts for the balance of forces at equilibrium. The next two equations define the chemical potentials of nucleoplasm and the epigenetic marks respectively which denote the thermodynamic driving force for their evolution in space and time, as discussed next in Section S1.2. In addition to the governing equations Eq (S4) which hold inside the nucleus, the following boundary conditions (emerging from the last two terms in Eq (S3)) hold at the surface of the nucleus,

$$s_{iK}N_K = t_i, \text{ (Emerges from Eq S3)}$$
$$\mu_n = \mu_n^{ext} \text{ (Specified at the boundary).} \tag{S5}$$

S1.2 Diffusion and reaction kinetics

The dynamic evolution of the nucleus towards the steady state is driven in a time-dependent fashion by a combination of diffusion as well as reaction kinetics. The local conservation of reactively inert nucleoplasm content relates the time evolution of local nucleoplasm volume fraction with its nominal volumetric flux in reference configuration $\boldsymbol{\mathcal{J}}^n$ as,

$$\frac{\partial \Phi_n}{\partial t} = -\nabla_K \mathcal{J}_K^n \tag{S6}$$

Here the nominal volumetric flux $\boldsymbol{\mathcal{J}}^n$ is the volume fraction of nucleoplasm moving across a reference surface of unit area. Physically, such flux involves the spatial movement of water that occurs to allow nucleosomes in the chromatin polymer to be redistributed driven by their self-interactions. For instance, methylated histones in vicinity coalesce together, with nucleoplasm moving away, driven by chromatin-chromatin interactions mediated by cross-linkers such as HP1$\alpha$ (Fig S1a). Such evolution leads to a reduction in energy, driving the local chromatin organization towards one of the energy wells in the landscape depicted in Fig 2b (main text). Thus, the diffusive flux of nucleoplasm is dictated by the energetic driving force, the chemical potential of nucleoplasm $\mu_n$ described in Eq (S4). In the current configuration, we can define the true volumetric flux of nucleoplasm $\boldsymbol{j}^n$, i.e. the volume fraction of nucleoplasm moving across per unit area in the current configuration as,

$$j_i^n = -\frac{D_n \Omega}{k_B T}\frac{\partial \mu_n}{\partial x_i} = -M_n \frac{\partial \mu_n}{\partial x_i} \tag{S7}$$

where, $M_n$ is the mobility of nucleoplasm in the nucleus, $D_n$ is its diffusivity, $\Omega$ is the molecular volume of water and $k_B T$ is the temperature in the unit of energy. The mobility $M_n$ is a measure of energy dissipated as water moves a unit length through the nuclear architecture and is isotropic in the current configuration. The true flux in the current state can be related to the nominal reference flux via the relation $\mathcal{J}_K = j_i H_{iK} J$ where $\boldsymbol{H} = \boldsymbol{F}^{-1}$, such that considering nominal variables we can rewrite Eq (S6) as,

$$\frac{\partial \Phi_n}{\partial t} = \nabla_K \left( M_n J H_{iK} H_{iL} \frac{\partial \mu_n}{\partial X_L} \right) \tag{S8}$$

On the other hand, the spatio-temporal evolution of epigenetic marks, i.e. the methyl or acetyl marks on the histones are prominently governed by their interconversion which occurs due to epigenetic reactions (Fig S1b). Methylation involves removal of acetyl group and addition of methyl group thereby forming heterochromatin phase, whereas acetylation involves removal of methyl and addition of acetyl group resulting in euchromatin phase (Fig 1c, Main text). Note that, by themselves, these reaction terms do not alter the total number of histones, only their epigenetic flavors. The acetylation and methylation reactions thus incorporated, however, occur at all points in the nucleus, i.e. at any histone with an equal rate $\Gamma_{ac}$ or $\Gamma_{me}$ respectively. The contribution of reactions to the evolution of epigenetic marks is captured via first order reaction kinetics as,

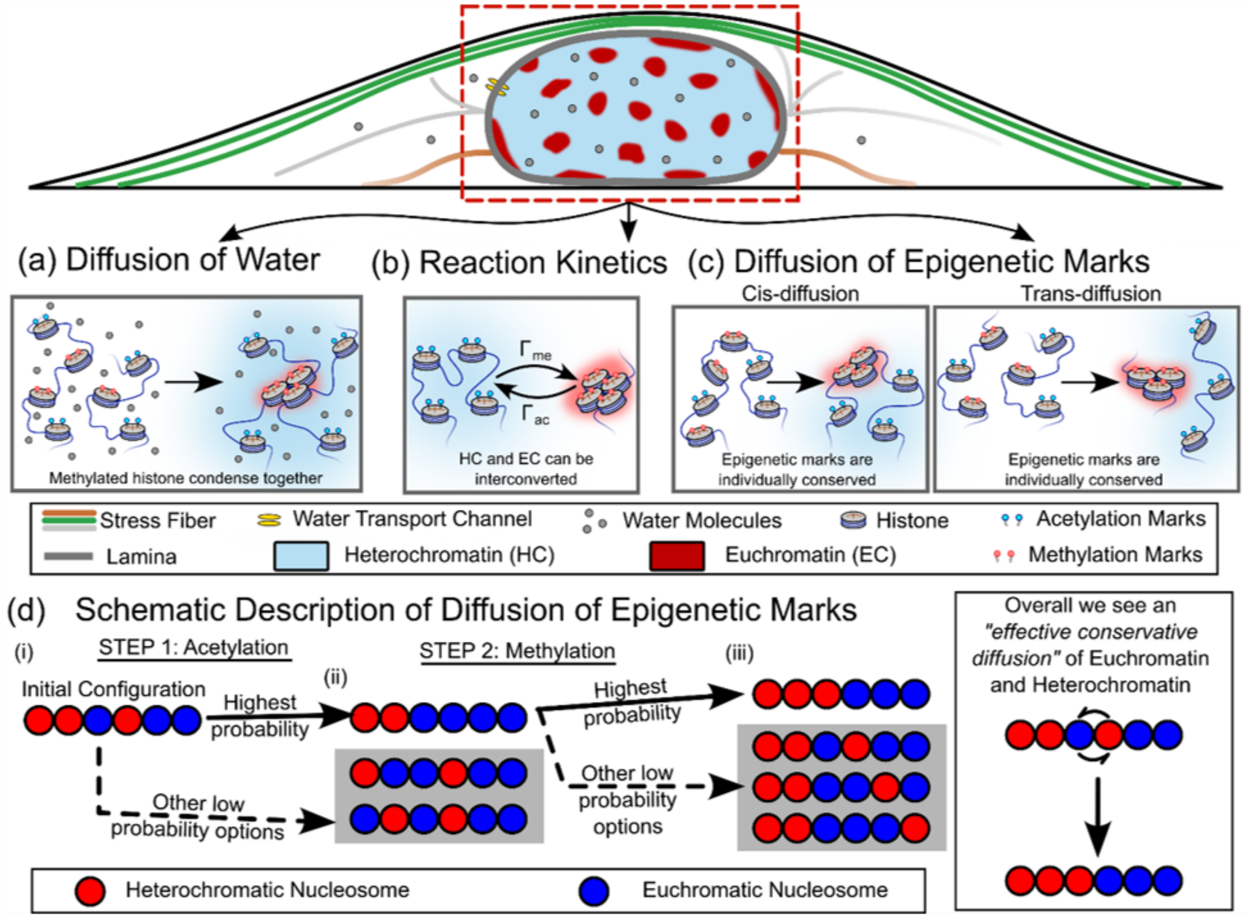

Fig S1: A schematic depicting the individual roles of the diffusion and reactions kinetics incorporated into the heterochromatin organization model. (a) Conservative diffusion of water which can redistribute molecules of water within the nucleus without changing the total amount of water, or total amount of hetero- or euchromatin in the nucleus. (b) Non-conservative reaction kinetics of histone acetylation and methylation, which allows an interconversion of chromatin phases. This changes the individual amounts of heterochromatin and euchromatin in the nucleus without changing the total amount of DNA. The reaction rates determine the ratio of heterochromatin to euchromatin at steady state. (c) The chromatin-chromatin interactions contribute to the reaction kinetics effectively driving a conservative evolution of epigenetic marks, which we call ‘diffusion of epigenetic marks’. (d) The diffusion of epigenetic marks is an overall result of reaction kinetics coupled with preference of like-like neighbor over unlike ones, thereby effectively rendering reactions ate certain sites more probable than others.

$$\frac{\partial \Phi_d}{\partial t} = \underbrace{2J(\Gamma_{me}\phi_a - \Gamma_{ac}\phi_m)}_{\text{Epigenetic Reactions}} \tag{S9}$$

Here we have assumed a first order reaction kinetics for acetylation of heterochromatin (with reaction rate $\Gamma_{ac}$) and methylation of euchromatin ($\Gamma_{me}$), although a more complicated reactions kinetics can be as easily used. Also, it must be noted that the reaction kinetics is written for the material volume fraction of eu- and heterochromatin (in the current configuration) and is then transformed into the reference configuration via the multiplication by $J$.

Due to the energetic interactions between neighboring histones, captured as chromatin-chromatin interactions, their epigenetic marks can also be influenced. For example, methylated histones can be cross-linked to neighboring methylated histones via proteins such as HP1. As a result, energetics dictates that a histone would prefer to stay methylated when its neighbors are prominently methylated, such in the heterochromatin phase. Thus, the rates of acetylation and methylation are not the same at all points in the nucleus, and instead depend on the epigenetic states of the neighbors.

To illustrate this better consider a section of chromatin polymer as the one depicted in Fig S1d(i), where nucleosomes with methylated histones are shown in red while those with acetylated histone tails are in blue. Let us say that acetylation occurs (this is without loss of generality, methylation could occur first, and the following statements will still hold, albeit reversed). Since acetylated histones prefer acetylated histone neighbors, the site on the chromatin polymer which is acetylated with the highest probability is the one which maximizes the likelihood of like-marked histones forming neighbors. Thus, the chromatin polymer configuration shown in Fig S1d(ii) is the most likely to occur, i.e. occurs with the highest reaction rates. While other sites could be acetylated, their probability is low, i.e. the acetylation reaction rate associated with the conversion of other sites is low. As a next step, if methylation occurs the site most likely to get converted is the one resulting in maximal red-red neighbors, thus resulting in a chromatin configuration shown in Fig S1d(iii). Thus, over the course of these two steps, we observe that the outcome with highest occurrence probability is where there is an effective switching of blue and red beads, i.e. a conservative evolution in time where the acetyl and methyl groups have effectively been interchanged with each other over time. Other possible outcomes which are non-conservative are effectively captured by the 'non-conservative' reaction terms in Eq (S9). The effective conservative evolution is captured in our model as the 'diffusion of epigenetic marks' (Fig S1c) as

$$\frac{\partial \Phi_d}{\partial t} = \underbrace{\nabla_K \left( M_d J H_{iK} H_{iL} \frac{\partial \mu_d}{\partial X_L} \right)}_{\text{Diffusion-like evolution}} \tag{S10}$$

where $M_d$ is the mobility of epigenetic marks. The diffusivity of epigenetic marks can be equivalently written as $\mathrm{D_d} = Mk_BT/\Omega$. To support the qualitative description of events which can give rise to the 'diffusion-like evolution' in Eq (S10), in Section S1.7 we present a quantitative proof of how such evolution is effectively emulated by the diffusion of epigenetic marks. The overall time-evolution of the epigenetic marks is given as,

$$\frac{\partial \Phi_d}{\partial t} = \underbrace{\nabla_K \left( M_d J H_{iK} H_{iL} \frac{\partial \mu_d}{\partial X_L} \right)}_{\text{Diffusion-like}} + \underbrace{2J(\Gamma_{me}\phi_a - \Gamma_{ac}\phi_m)}_{\text{Epigenetic Reactions}} \tag{S11}$$

Equations (S4), (S8) and (S11) together govern the evolution of the nucleus towards a steady state under boundary conditions prescribed by Eq (S5).

#### *S1.3 Molecular Incompressibility Enables Differential Swelling*

Local flux of nucleoplasm at any infinitesimal region in the nucleus, via Eq (S8), alters the number of space-occupying water molecules, and thus changes the local volume of the region with time. Note, however, that flux of epigenetic marks only alters the histone-tail posttranslational modification, without changing the total number of histones, and thus the local volume. This relation between the volumetric deformation and the local volume fraction of nucleoplasm is called the molecular incompressibility constraint.

Say a region of nucleus has a reference volume (reference is defined as free of nucleoplasm) of $V_0$. After an influx of nucleoplasm of volume $V_n$, the current volume of the region increases to $V_c$. Molecular incompressibility dictates,

$$V_c - V_0 = V_n \tag{S12}$$

Dividing both sides by $V_0$ and noting that $\frac{V_n}{V_0} = \Phi_n$ and $\frac{V_c}{V_0} = J$,

$$J - 1 - \Phi_n = 0 \tag{S13}$$

Thus, the constraint given by Eq (S13) relates the local swelling with the local nucleoplasm content. In current configuration, we can rewrite Eq (S13) as, $J = \frac{1}{1-\phi_n}$. Thus, dictated by the molecular incompressibility constraint, local expulsion of nucleoplasm would result in chromatin compaction (decrease in $J$) resulting in formation of genetically silenced heterochromatin domain. On the other hand, euchromatin would swell ($J$ increases) due to nucleoplasm influx resulting in a loose chromatin polymer network.

Note that Eq (S13) can be averaged over the entire nucleus (with total volume of the nucleus in reference and current configurations being $V_{nuc}^R$ and $V_{nuc}^c$ respectively) as:

$$\frac{1}{V_{nuc}^R} \int_{V_{nuc}^R} (J - 1 - \phi_n)\, dV = 0$$

$$\Rightarrow V_{nuc}^c = (1 + \bar{\phi}_n) V_{nuc}^R$$

which gives the global relation between the nuclear volume and the overall water content in the nucleus $\bar{\phi}_n = \frac{1}{V_{nuc}^R} \int_{V_{nuc}^R} \phi_n dV$. Note that $\bar{\phi}_n$ is effectively $\phi_n$ averaged over the entire reference nuclear volume. When the overall water content in the nucleus increases, the nuclear volume increases.

#### *S1.4 Specific Energy Description to Enable Phase Separation*

In this section, we invoke the explicit form of free energy density $W(\boldsymbol{F}, \Phi_n, \Phi_d, \boldsymbol{\nabla\Phi_n}, \boldsymbol{\nabla\Phi_d})$ used in Eq (S2) that enables the separation of euchromatic and heterochromatic phases in the nucleus. We begin incorporating the following contributions to the free energy density,

$$W(\boldsymbol{F}, \Phi_n, \Phi_d, \boldsymbol{\nabla\Phi_n}, \boldsymbol{\nabla\Phi_d}) = W_m(\boldsymbol{F}, \Phi_n, \Phi_d) + W_L(\Phi_m, \boldsymbol{F}) + W_e(\boldsymbol{F}) + W_{int}^n(\boldsymbol{\nabla\Phi_n}) + W_{int}^d(\boldsymbol{\nabla\Phi_d}) \tag{S14}$$

where, all free energy density contributions are written in the reference configuration.

The first term in Eq (S14) $W_m(\boldsymbol{F}, \Phi_n, \Phi_d)$ denotes the energy associated with the entropic and enthalpic interactions between nucleosomes with acetylated or methylated histones. It is equivalent to the Flory-Huggins type free energy description [2, 3] in which energetic interactions between histones drive their segregation and entropic contributions favor the mixing of chromatin constituents. Over a short-range, nucleosomes interact with each other via ionic interactions or physical tethering mediated by nucleosome bridging proteins such as HP1$\alpha$. These interactions bring like-marked nucleosomes together, while such segregation is unfavorable from an entropic perspective which rather prefers a more homogenized distribution of acetylated and methylated histones. In addition, locally driven energetic perturbations, such as motor activity or transcription, or ATP consumption, can further drive an 'activity-induced' segregation of the chromatin phases. These 'chromatin-chromatin interactions' are captured by constructing a double-well energy landscape which gives rise to two stable phases of chromatin (Fig S2) – a water-rich euchromatin phase ($\phi_a > \phi_m$, and $\phi_n$ high), and a water-poor heterochromatin phase ($\phi_m > \phi_n$ and $\phi_n$ very small) such that,

$$W_m(\boldsymbol{F}, \Phi_n, \Phi_d) = \frac{k_B T}{\Omega}[J\phi_a^2 + J\phi_m^2(\phi_m^{max} - \phi_m)^2] \tag{S15}$$

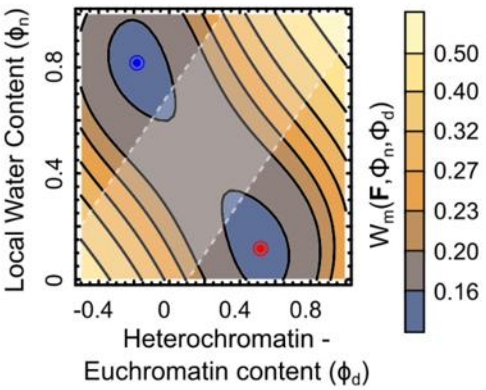

Fig S2: Contour plot of energetics of chromatin-chromatin interaction, the first term in Eq S12

where, $\phi_m^{max}$ determines the extent of compaction of methylated histones in the heterochromatin phase and the coefficient $k_B T/\Omega$ emerges as the scale of energy density.

Some of the heterochromatin regions formed as a result of the energetic segregation can be sequestered to the nuclear periphery. At the periphery, methylated histones can be tethered to chromatin-anchoring proteins such as LAP2$\beta$, emerin, MAN1, etc. [4-6] present in the nuclear lamina forming lamina-associated domains (LADs). Such chromatin-lamina interactions are captured in our model via the second term in Eq (S14), as $W_L(\Phi_m, \boldsymbol{F}) = V_L \Phi_m e^{-d/d_0}$. Here $V_L$ denotes the strength of anchoring proteins tethering the methylated histones to the lamina. The interactions decay exponentially away from the lamina over a length-scale $d_0$.

The changes in the local volume fractions of acetylated and methylated histones, resulting in local compaction or swelling result in the deformation of the chromatin polymer which is modelled as a neo-Hookean solid incorporating the non-linearities associated with finite strains. The elastic energy, penalizing such strains associated with the deformations give rise to the third term in Eq (S14), as,

$$\begin{aligned} W_e(\boldsymbol{F}) &= \frac{G(\phi_h)}{2}[F_{iK}F_{iK} - 3 - 2\ln J] \\ G(\phi_m) &= G_{EC}\left[1 + \frac{\delta G}{\phi_m^{max}}\phi_m\right] \end{aligned} \tag{S16}$$

where, $G$ is the shear modulus of the chromatin polymer. It has been previously [7] shown via nuclear elastography that in its compacted phase heterochromatin is relatively stiffer as compared to euchromatin. Consistent with the experimental observations, we allow the shear modulus to be dependent on the levels of local histone methylation via Eq (S16) such that the heterochromatin phase is twice as stiffer than the euchromatin phase, i.e., $G_{HC} = 2G_{EC}$. The ratio of stiffness can be adjusted according to the stiffness of chromatin phases in specific cell-lines. The energy landscape (shown in Fig S2) gets altered due to contribution of strain energy as shown in Fig 2b. The quantitative role of strain energy in altering the energy landscape is further discussed in Section S7.1.

The last two terms in Eq (S14) account for the energy associated with formation of boundaries between the phase separating heterochromatin and euchromatin region. The phase boundaries are defined in terms of the spatial gradients in the volume fractions $\Phi_n$ and $\Phi_d$ such that we define,

$$
\begin{aligned}
W_{int}^{n}(\boldsymbol{\nabla\Phi_n}) &= \frac{\eta_n}{2} J \frac{\partial \Phi_n}{\partial x_i} \frac{\partial \Phi_n}{\partial x_i} = \frac{\eta_n}{2} J H_{Mj} \frac{\partial \Phi_n}{\partial X_M} \frac{\partial \Phi_n}{\partial X_L} H_{Lj} \\
W_{int}^{d}(\boldsymbol{\nabla\Phi_d}) &= \frac{\eta_d}{2} J \frac{\partial \Phi_d}{\partial x_i} \frac{\partial \Phi_d}{\partial x_i} = \frac{\eta_d}{2} J H_{Mj} \frac{\partial \Phi_d}{\partial X_M} \frac{\partial \Phi_d}{\partial X_L} H_{Lj}
\end{aligned} \tag{S17}
$$

where, $\eta_{n,d}$ measures the increase in energy due to formation of interface of unit width in the current configuration. As $\eta_{n,d}$ increases there is a greater penalty for the formation of interfaces, resulting in formation of smoother and wider interfaces. In our simulations we choose $\eta_n = \eta_d = \eta$.

Additionally, the constraint of molecular incompressibility (Eq (S13)) is implemented numerically by adding an energy penalty $W_{inc}$ to the free energy density given as,

$$W_{inc} = \frac{\kappa}{2}(J - 1 - \Phi_n)^2 \tag{S18}$$

where $\kappa$ is a parameter that penalizes a deviation in the volume from that imposed by the incompressibility constraint. Physically $\kappa$ corresponds to the bulk modulus. Thus, the free energy density in Eq (S14) is given as,

$$
\begin{aligned}
&W(\boldsymbol{F}, \Phi_n, \Phi_d, \boldsymbol{\nabla\Phi_n}, \boldsymbol{\nabla\Phi_d}) \\
&\quad = \frac{G}{2}[F_{iK}F_{iK} - 3 - 2\ln J] + J\phi_e^2 + J\phi_h^2(1-\phi_h)^2 \\
&\quad + \frac{\eta_n}{2} J H_{Mj} \frac{\partial \Phi_n}{\partial X_M} \frac{\partial \Phi_n}{\partial X_L} H_{Lj} + \frac{\eta_d}{2} J H_{Mj} \frac{\partial \Phi_d}{\partial X_M} \frac{\partial \Phi_d}{\partial X_L} H_{Lj} + \frac{\kappa}{2}(J - 1 - \Phi_n)^2 \\
&\quad + \text{viscous term}
\end{aligned} \tag{S19}
$$

where the last term accounts for the energy dissipation due to the viscous deformation of chromatin. Considering the specific form of free energy defined in Eq (S19), the nominal stress defined using Eq (S3) can be written as,

$$
\begin{aligned}
s_{iK} = \underbrace{\frac{\partial W_e}{\partial F_{iK}}}_{\text{neo-Hookean Elastic Stress}} &+ \underbrace{\left\{\frac{\partial W_m}{\partial J} + \kappa(J - 1 - \Phi_n)\right\}}_{\text{osmotic pressure}} J H_{Ki} + \underbrace{\frac{\partial\left(W_{int}^{n} + W_{int}^{d}\right)}{\partial F_{iK}}}_{\text{interfacial surface tension}} \\
&+ \underbrace{\theta J H_{Lj}\left(\frac{1}{2} H_{Kj} \frac{\partial v_i}{\partial X_L} + \frac{1}{2} H_{Ki} \frac{\partial v_j}{\partial X_L} - \frac{1}{3} H_{Ki} \frac{\partial v_j}{\partial X_L}\right)}_{\text{Distortional stress due to viscous dissipation}}
\end{aligned} \tag{S20}
$$

where, $\theta$ is the viscosity of chromatin and $\boldsymbol{v} = \frac{\partial \boldsymbol{x}(\boldsymbol{X},t)}{\partial t}$ denotes the velocity vector in the current configuration. Similarly, the chemical potential driving the kinetics of nucleoplasm and difference between hetero and euchromatin can be written as,

$$\mu_n = \frac{\partial W_m}{\partial \Phi_n} - \nabla_K\left(\frac{\partial W_{int}^{n}}{\partial \nabla_K \Phi_n}\right) - \kappa(J - 1 - \Phi_n), \tag{S21}$$

$$\mu_d = \frac{\partial W_m}{\partial \Phi_d} - \nabla_K \left( \frac{\partial W_{int}^d}{\partial \nabla_K \Phi_d} \right)$$

Eq S4, S10, S11, S20 and S21 together form the time-dependent governing equations which describe the spatiotemporal evolution of chromatin organization in the nucleus. In Table S1, we list the mathematical symbols used in our model, their physical meaning and units.

*S1.5* *Coarse-grained treatment of ionic and electrostatic interactions:*

Chromatin is a highly charged polymeric material, and electrostatic interactions between DNA, histones, and other nuclear constituents contribute to chromatin compaction, hydration, and intermolecular interactions. Our framework does not explicitly resolve mobile ionic species or the electrostatic potential as independent fields. Instead, their collective effects are coarse-grained into the effective free-energy landscape governing chromatin organization. Specifically, ionic contributions to interactions between chromatin constituents are incorporated into the chromatin-chromatin interaction term in Eq. (S14), whose explicit form is given in Eq. (S15). The EC and HC phases are consequently distinguished by their effective thermodynamic interactions, nucleoplasm content, and volumetric compaction, while osmotic exchange between the nucleus and cytoplasm is described through the chemical potential of nucleoplasm prescribed at the nuclear boundary. This coarse-grained description allows us to focus on the coupled mesoscale evolution of chromatin mechanics, epigenetic modifications, and nucleoplasm transport without resolving the molecular degrees of freedom underlying the effective thermodynamic interactions.

More detailed continuum descriptions of charged polymer networks and polyelectrolyte gels treat the concentrations of fixed polymer charges, mobile ionic species, and solvent as separate fields and explicitly couple them to the electrostatic potential through Maxwell electrostatics. Such models clarify when these additional fields need to be retained. In particular, Celora et al. [8] showed that the behavior depends on the relative magnitude of the Debye screening length and the diffuse-interface length scale associated with phase separation. When the Debye length is much smaller than the interfacial length scale, the bulk gel remains approximately electroneutral and electric double layers do not control the large-scale phase-separation dynamics. Nevertheless, solvent transport continues to produce swelling and deswelling, coexistence of differently hydrated phases, propagating phase-transition fronts, and spinodal decomposition. In contrast, when the Debye and interfacial length scales become comparable, local electroneutrality can break down and explicit electrostatics can generate additional spatially localized charged structures [8]. Thus, explicit charge resolution is essential for describing Debye-layer structure and charge-separated patterns, but is not required for the longer-wavelength swelling and phase-separation phenomena that are the focus of the present framework.

The characteristic length scale of electrostatic screening under physiological conditions is substantially smaller than the mesoscale chromatin structures considered here. At physiological ionic strengths of approximately 0.1−0.15 M, the Debye length of the order 1 nm or smaller $\left(\text{Debye length} = \lambda_D \approx \frac{0.3}{\sqrt{0.1\,M}} < 1\text{nm}\right)$, whereas the HC domains resolved in our framework are of order $10^2$ nm. The model also contains an intrinsic diffuse-interface length scale associated with the gradient-energy terms in Eq. (S17), as discussed in Section S1.6. We do not attempt to resolve the electrostatic structure within this interface. Rather, the present model is intended to describe chromatin organization on length scales much larger than the screening length, for which the effects of rapidly screened electrostatic interactions can be represented through effective thermodynamic parameters.

A corresponding separation exists in time. Mobile ions redistribute over submicron distances much faster than the epigenetic reactions, nucleoplasm redistribution, and mesoscale chromatin reorganization described by our model. For example, for a characteristic distance of 100 nm, and an ionic diffusivity of order $10^{-10}\mathrm{m}^2/s$, the diffusive redistribution time $L^2/D$ is of the order $10^{-4}$ s. This is many orders of magnitude shorter than the characteristic epigenetic reaction times used in our model, for which the methylation rate is of order $10^3$ s. Ionic degrees of freedom can therefore be regarded as rapidly relaxing relative to the slow variables explicitly retained in the theory.

Our treatment can thus be viewed as a further coarse-graining of the electroneutral description used in polyelectrolyte-gel theories. Rather than retaining locally electroneutral ionic concentrations as additional dynamical variables, we incorporate their equilibrated contributions into the effective chromatin-chromatin free energy. This approximation preserves the mesoscale competition among chromatin interactions, nucleoplasm partitioning, differential swelling, elasticity, interfacial energy, and epigenetic reaction kinetics that govern the phenomena studied here. It is not intended to describe salt-specific ion partitioning, electric-double-layer structure, local charge-density variations, or regimes in which the Debye length becomes comparable to the chromatin interfacial width. Within this scope, the coarse-grained treatment provides a minimal description of the chemo-mechano-osmotic coupling responsible for EC-HC phase organization while avoiding additional fast microscopic degrees of freedom that are not explicitly resolved by the present mesoscale model.

#### *S1.6 Rescaling the Governing Equations*

The governing equations derived in the previous subsections can be rescaled with respect to the intrinsic length and time scales arising from the model. The coefficient of the chromatin-chromatin interactions in Eq (S15), $k_B T/\Omega$ is chosen as the scale of energy density. All energy densities are then normalized such that $\widetilde{W} = W\Omega/k_B T$ and the rescaled chemical potentials are $\tilde{\mu} = \mu\Omega/k_B T$. Consequently, the normalized shear and bulk modulus are obtained as $\tilde{G} = G\Omega/k_B T$, and $\tilde{\kappa} = \kappa\Omega/k_B T$ respectively. Similarly, the strength of chromatin-lamina interactions is normalized to obtain $\tilde{V}_L = V_L\Omega/k_B T$. The normalized free energy density can thus be written as,

$$\widetilde{W} = \underbrace{\widetilde{W}_m(\boldsymbol{F}, \Phi_n, \Phi_d)}_{\substack{\text{chromatin-chromatin} \\ \text{interactions}}} + \underbrace{\widetilde{W}_e(\boldsymbol{F})}_{\substack{\text{elastic strain} \\ \text{energy}}} + \underbrace{\tilde{V}_L \Phi_m e^{-d/d_0}}_{\substack{\text{chromatin-lamina} \\ \text{interactions}}} + \underbrace{\widetilde{W}_{int}^n(\boldsymbol{\nabla\Phi_n}) + \widetilde{W}_{int}^d(\boldsymbol{\nabla\Phi_d})}_{\text{Interfacial energy}} + \frac{\tilde{\kappa}}{2}(J - 1 - \Phi_n)^2 \quad \text{(S22)}$$

Note that the interfacial energies $\widetilde{W}_{int}^{n,d}$ in Eq (S22) are also normalized with respect to the scale of energy densities. As a consequence of energy density normalization, the first Piola-Kirchoff stress is also normalized in a similar fashion such that $\tilde{\boldsymbol{s}} = \frac{\partial \widetilde{W}}{\partial \boldsymbol{F}} = \frac{s\Omega}{k_B T}$. The competition between the interfacial energies, which scale as $\eta$ and the bulk energy densities which scale as $k_B T/\Omega$ gives rise to an intrinsic length scale $\ell_{int} = \sqrt{\eta\Omega/k_B T}$. Physically, $\ell_{int}$ corresponds to the width of the interface formed between the heterochromatin and the euchromatin phases. Although $\ell_{int}$ can be used as the rescaling parameter for all lengths in our model, a mesoscopic length scale that emerges from the non-equilibrium kinetic considerations is more appropriate. The evolution of epigenetic marks given by Eq (S11) is a reaction-diffusion equation which gives rise to a characteristic length-scale from the balance between the reaction and diffusion kinetics called the reaction-diffusion length-scale $\ell_{RD} = \sqrt{M_D k_B T/\Omega\Gamma_{ac}} = \sqrt{D_d/\Gamma_{ac}}$. All lengths in our model are non-dimensionalized with respect to $\ell_{RD}$ such that $\tilde{\boldsymbol{x}} = \boldsymbol{x}/\ell_{RD}$. Note that the rescaled width of the interface is $\delta = \ell_{int}/\ell_{RD}$. Lastly, the reaction kinetics offer an intrinsic time scale for our equations such that all times are rescaled as $\tilde{t} = t\Gamma_{ac}$. The governing equations of our model, Eq (S4), (S8) and (S11) can thus be rescaled as,

$$\begin{aligned}
\nabla_K \tilde{s}_{iK} &= 0, \\
\tilde{\mu}_n(\tilde{\boldsymbol{x}}, t) &= \frac{\partial \widetilde{W}}{\partial \Phi_n} - \nabla_K \left( \frac{\partial \widetilde{W}}{\partial \nabla_K \Phi_n} \right), \\
\mu_d(\tilde{\boldsymbol{x}}, t) &= \frac{\partial \widetilde{W}}{\partial \Phi_d} - \nabla_K \left( \frac{\partial \widetilde{W}}{\partial \nabla_K \Phi_d} \right) \\
\frac{\partial \Phi_n}{\partial \tilde{t}} &= \nabla_K \left( J H_{iK} H_{iL} \frac{\partial \tilde{\mu}_n}{\partial \tilde{X}_L} \right) \\
\frac{\partial \Phi_d}{\partial \tilde{t}} &= \underbrace{\nabla_K \left( J H_{iK} H_{iL} \frac{\partial \tilde{\mu}_d}{\partial \tilde{X}_L} \right)}_{\text{Diffusion-like}} + \underbrace{2J\left(\tilde{\Gamma}_{me}\phi_a - \phi_m\right)}_{\text{Epigenetic Reactions}}
\end{aligned} \tag{S23}$$

### *S1.7 Neighborhood dependent reaction kinetics are captured by diffusion-like evolution*

In Section S1.2 we have presented a qualitative scenario where the effect of chromatin-chromatin interactions on the reaction kinetics of acetylation and methylation can lead to an effective evolution of epigenetic marks such that the like-marked nucleosomes gather together resulting in coarsening of heterochromatin. Here we theoretically prove that such evolution follows diffusion-like kinetics, such that purely by asserting the neighborhood-dependence of epigenetic reactions the spatio-temporal evolution of epigenetic marks described by Eq S11 is recovered.

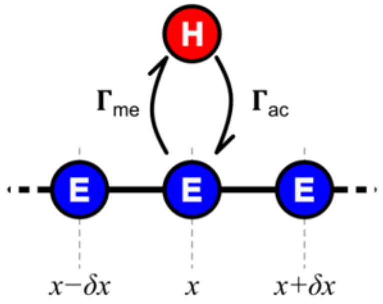

Fig S3: A representation of a section of chromatin polymer.

To simulate the chromatin polymer consider a series of nucleosomes as shown in Fig S3. Consider a nucleosome at location $x$ with left and right neighbors at locations $x - \delta x$ and $x + \delta x$ respectively. Let $P_H(x)$ denote the probability that the nucleosome at location $x$ is methylated. Thus the probability that the nucleosome at location $x$ is acetylated is $P_E(x) = 1 - P_H(x)$. The nucleosome can be interconverted from acetylated into methylated forms and vice-versa at a rate $\Gamma_{me}$ and $\Gamma_{ac}$ respectively. Focusing only at location $x$, the probability that the nucleosome at this location is methylation evolves in time as,

$$\frac{\partial P_H(x,t)}{\partial t} = -\Gamma_{ac} P_H + \Gamma_{me} P_E \tag{S24}$$

Now let us consider a scenario where at a location $x$ the rates of acetylation and methylation are not constant but dependent on the epigenetic states of the left and right neighbors. This is because any interconversion of the histone marks also requires breaking the bonds between neighbors. Taking the example depicted in Fig S3, say a nucleosome is acetyl marked and thus

E (euchromatic), and needs to be converted into H (heterochromatic), depending on the left and right neighbors there are four possible options for such a methylation step:

| | Configuration | | Change in energy | Methylation rate |
|---|---|---|---|---|
| 1 | EEE | Need to break two eu-eu bonds | $\Delta\mathrm{E} = 2\mathrm{E}_{EE}$ | $\Gamma_{me}$ |
| 2 | EEH | Need to break one eu-eu and one eu-hetero bond | $\Delta\mathrm{E} = \mathrm{E}_{EE} + E_{EH}$ | $\sim \Gamma_{me} + \Delta\Gamma_{\mathrm{me}}$ |
| 3 | HEE | | | |
| 4 | HEH | Need to break two eu-hetero bonds | $\Delta\mathrm{E} = 2E_{EH}$ | $\sim \Gamma_{me} + \Delta\Gamma'_{\mathrm{me}}$ |

Since like-marked nucleosomes interact strongly as compared to unlike marked nucleosomes, $E_{EH} < E_{HH}, E_{EE}$. Thus, methylation occurs more easily, i.e. methylation rate is higher ($\Delta\Gamma_{\mathrm{me}}, \Delta\Gamma'_{me} > 0$) when a bond between unlike nucleosomes is broken than if that between like-marked nucleosome is broken. Similarly, we can have four possible options for $\Gamma_{ac}$. Considering these eight possible options after taking into account the effects of the neighbors, the probability that nucleosome at site $x$ is heterochromatic evolves in time as,

$$\frac{\partial P_H(x,t)}{\partial t} = P_E\left[\underbrace{P_E^+P_E^-\Gamma_{me}}_{EEE} + \underbrace{P_E^+P_H^-(\Gamma_{me}+\Delta\Gamma_{\mathrm{me}})}_{EEH} + \underbrace{P_H^+P_E^-(\Gamma_{me}+\Delta\Gamma_{\mathrm{me}})}_{HEE} + \underbrace{P_H^+P_H^-(\Gamma_{me}+\Delta\Gamma'_{\mathrm{me}})}_{HEH}\right] - P_H\left[\underbrace{P_H^+P_H^-\Gamma_{ac}}_{HHH} + \underbrace{P_H^+P_E^-(\Gamma_{ac}+\Delta\Gamma_{\mathrm{ac}})}_{HHE} + \underbrace{P_E^+P_H^-(\Gamma_{ac}+\Delta\Gamma_{\mathrm{ac}})}_{EHH} + \underbrace{P_E^+P_E^-(\Gamma_{ac}+\Delta\Gamma'_{\mathrm{ac}})}_{EHE}\right] \quad \text{(S25)}$$

Here, $P^+_{H(E)} = P_{H(E)}(x+\delta x)$ is the probability that the right neighbor is heterochromatic (euchromatic). Similarly, $P^-_{H(E)} = P_{H(E)}(x-\delta x)$ is the probability that the left neighbor is heterochromatic (euchromatic). For simplification, we begin with the first bracket term on the right side of Eq (S25):

$$P_E[P_E^+P_E^-\Gamma_{me} + P_E^+P_H^-(\Gamma_{me}+\Delta\Gamma_{\mathrm{me}}) + P_H^+P_E^-(\Gamma_{me}+\Delta\Gamma_{\mathrm{me}}) + P_H^+P_H^-(\Gamma_{me}+\Delta\Gamma'_{\mathrm{me}})]$$

$$= P_E[\Gamma_{me}\{P_E^+P_E^- + P_E^+P_H^- + P_H^+P_E^- + P_H^+P_H^-\} + \Delta\Gamma_{\mathrm{me}}\{P_E^+P_H^- + P_H^+P_E^- + 2P_H^+P_H^-\}]$$

Here, as a simplification, we have assumed that $\Delta\Gamma'_{me} = 2\Delta\Gamma_{me}$. This simplification is only for the conciseness of the present derivation and does not affect the qualitative conclusion from this section. At the end of this section, for more generality, we have also considered the case when $\Delta\Gamma'_{me} = k\Delta\Gamma_{me}$. Next, we use the fact that $P_H + P_E = 1$, $P_H^+ + P_E^+ = 1$ and $P_H^- + P_E^- = 1$.

$$= P_E[\Gamma_{me} + \Delta\Gamma_{\mathrm{me}}\{(1-P_H^+)P_H^- + P_H^+(1-P_H^-) + 2P_H^+P_H^-\}]$$

$$= P_E[\Gamma_{me} + \Delta\Gamma_{\mathrm{me}}\{P_H^- - P_H^-P_H^+ + P_H^+ - P_H^+P_H^- + 2P_H^+P_H^-\}]$$

$$= P_E[\Gamma_{me} + \Delta\Gamma_{\mathrm{me}}\{P_H^+ + P_H^-\}]$$

Similarly, the second bracket term can be simplified, thereby reducing Eq (S25) as,

$$\frac{\partial P_H(x,t)}{\partial t} = -P_H[\Gamma_{ac} + \Delta\Gamma_{\mathrm{ac}}\{2 - P_H^+ - P_H^-\}] + P_E[\Gamma_{me} + \Delta\Gamma_{\mathrm{me}}\{P_H^+ + P_H^-\}] \tag{S26}$$

Note that if the neighborhood effect is switched off (i.e., $\Delta\Gamma_{\mathrm{me}} = \Delta\Gamma_{ac} = 0$), Eq (S26) reduces to Eq (S24) derived above for the case of constant reaction rates. We can simplify Eq (S26) further as,

$$\frac{\partial P_H(x,t)}{\partial t} = -\Gamma_{ac}P_H + \Gamma_{me}P_E - \Delta\Gamma_{\mathrm{ac}}\{2P_H - (r - rP_H + P_H)(P_H^+ + P_H^-)\} \tag{S27}$$

where, $r = \Delta\Gamma_{me}/\Delta\Gamma_{ac}$. Next we can expand the left and right probabilities using Taylor expansion to the second order as $P_H^{\pm} = P_H(x \pm \delta x) = P_H(x) \pm \frac{\partial P_H}{\partial x}\delta x + \frac{\partial^2 P_H}{\partial x^2}\frac{(\delta x)^2}{2}$. After simplification, we get the evolution of heterochromatin marks with time as,

$$\begin{aligned}\frac{\partial P_H(x,t)}{\partial t} &= \underbrace{[-\Gamma_{ac}P_H + \Gamma_{me}P_E - 2(\Delta\Gamma_{\mathrm{me}} - \Delta\Gamma_{\mathrm{ac}})P_E P_H]}_{\text{Reaction term}} \\ &\quad + \underbrace{\underbrace{(\delta x)^2\{\Delta\Gamma_{\mathrm{me}}P_E + \Delta\Gamma_{\mathrm{ac}}P_H\}}_{\text{Effective Diffusivity}}\left\{\frac{\partial^2 P_H}{\partial x^2}\right\}}_{\text{Diffusion-like term}}\end{aligned} \tag{S28}$$

Comparing Eq (S28) above with Eq (S23) used in our model, we immediately see similarity of the reaction and diffusion-like terms in the evolution of epigenetic marks. While the first term in Eq (S28) is non-conservative and follows a first-order reaction-like kinetics, the second term is conservative and obeys diffusion like kinetics. Thus, we have quantitatively, from first principles, shown how epigenetic reaction kinetics, in conjunction with the energetics of chromatin-chromatin interactions can give rise to effective ‘diffusion of epigenetic marks’.

During the above derivation we assumed that $\Delta\Gamma'_{me} = 2\Delta\Gamma_{me}$. However, this is not a strict requirement for the derivation and was done only to simplify the process. In general, we can say that $\Delta\Gamma'_{me} = k\Delta\Gamma_{me}$. Under such scenario Eq (S28) is instead obtained as,

$$\begin{aligned}\frac{\partial P_H(x,t)}{\partial t} &= \left[-\Gamma_{ac}P_H + \Gamma_{me}P_E - \frac{\Delta\Gamma_{\mathrm{me}} + \Delta\Gamma_{\mathrm{ac}}}{2}(k-2)P_E P_H - 2(\Delta\Gamma_{\mathrm{me}} - \Delta\Gamma_{\mathrm{ac}})(k-1)P_E P_H\right] \\ &\quad + \underbrace{(\delta x)^2\{\Delta\Gamma_{\mathrm{me}}P_E + \Delta\Gamma_{\mathrm{ac}}P_H\}\left\{\frac{\partial^2 P_H}{\partial x^2} + (k-2)\left(\frac{\partial P_H}{\partial x}\right)^2\right\}}_{\text{Non-linear Diffusion-like term}}\end{aligned}$$

which again incorporates the reaction and diffusion terms, except this time the diffusion is non-linear, non-Fickian arising out of a possible cooperativity between neighboring E-H bonds which overrides the stoichiometric relationship $\Delta\Gamma'_{me} = 2\Delta\Gamma_{me}$. When $k = 2$, Eq S28 is recovered.

#### *S1.8 Choice of model parameters, initial and boundary conditions*

The parameter choice made for the simulations carried out in this work are motivated here. Broadly, we have the following parameter choices –

*Initial Conditions*: We consider an initially homogenous distribution of chromatin and nucleoplasm within the nucleus. Based on our experimental observations, and prior results [9, 10], we choose the initial overall nucleoplasm content in the nucleus as $\bar{\phi}_n^0 = 0.52$. Unless there is an exchange of water between the nucleoplasm and the cytoplasm due to osmotic perturbations, this value is maintained constant. In accordance with the incompressibility constraint (Eq S13), this corresponds to an initial swelling given as $J^0 = \frac{1}{1-\bar{\phi}_n^0} = 2.08$. This implies that the water content in the nucleus results in a 2.08 times increase in the nuclear volume, with respect to the nucleoplasm-free reference configuration. All future changes in nuclear volume are measured with respect to this initial swelling $J^0$.

Note that the initial choice of the overall volume fractions of acetylated and methylated histones is immaterial since the reactions will overwrite the initial concentration within the reaction-driven timescale $\left(\frac{1}{\Gamma_{ac}+\Gamma_{me}}\right)$ which is much faster than the timescale for HC domain nucleation. Secondly, to mimic the spatial heterogeneities intrinsic to the nucleus, we add a uniform perturbation to the initial homogenous chromatin configuration with range $\sigma_{\phi}^{noise} = 0.01$.

_Boundary Conditions_: Starting from the above-described initial conditions, we allow the chromatin organization to evolve towards a steady state which describes the chromatin organization in a control nucleus – in the absence of any external chemo-mechano-osmotic perturbations. For achieving this steady state, we allow no exchange of water between the nucleus and cytoplasm $\left(\nabla\mu_n.\hat{\boldsymbol{n}}|_{\text{boundary}} = 0\right)$. Further, since all chromatin is confined to within the nucleus, we also enforce $\nabla\mu_d.\hat{\boldsymbol{n}}|_{\text{boundary}} = 0$, which serves as the boundary condition for the chromatin order parameter. These two conditions allow the conservation of epigenetic marks and nucleoplasm. Lastly, we do not yet apply any forces on the nucleus (till the control steady-state is reached), such that the nuclear periphery is traction free $\left(\boldsymbol{s}\,\hat{\boldsymbol{n}}|_{\text{boundary}} = \boldsymbol{0}\right)$, where $\boldsymbol{s}$ denotes the nominal stress defined in Eq S20. These boundary conditions are together applied to the governing conditions to obtain the steady state chromatin organization in the nucleus.

When applying perturbations to epigenetic regulation, osmotic and mechanical forces, we emulate the nucleus within the cytoplasm by allowing the exchange of water between the nucleus and the cytoplasm. We first measure the steady state chemical potential of water in the cytoplasm, which is equal to the steady state chemical potential of water in the nucleus at steady state $\left(\mu_n^{cyt}\right)$. Then we set the boundary condition for water as $\mu_n|_{\text{boundary}} = \mu_n^{cyt}$, while the other boundary conditions remain the same. This ensures that when epigenetic perturbations act, or mechanical forces are applied, a free exchange of water is permitted between the nucleus and the cell. Furthermore, osmotic perturbations are captured by altering the boundary condition for water chemical potential as $\mu_n|_{\text{boundary}} = \mu_n^{cyt} \pm \Delta\mu_n$, with positive change for hypo-osmotic and negative for hyper-osmotic environments.

Lastly, to apply mechanical forces, traction forces (with dimensions of stress) are applied to the nuclear periphery along the outer normal of the surface via the boundary condition $\boldsymbol{s}\cdot\hat{\boldsymbol{n}}|_{\text{boundary}} = \boldsymbol{s^{ext}}$. Here $\boldsymbol{s^{ext}}$ contains the cartesian components of the external force acting on the nucleus. A positive (negative) value denotes outward (inward) traction along the normal direction.

_Model Parameters_: We use the following model parameters in the simulations:

Table 1: Values of parameters used in simulations

| | Parameter | Description | Value | Remark |
|---|---|---|---|---|

| | | | | |
|---|---|---|---|---|
| Initial Conditions | $\phi_n^0$ | Initial water (nucleoplasm) content in the nucleus | 0.52 | Established from experimental H2B density |
| | $\phi_d^0$ | Initial value of order parameter | 0.1 | The choice does not affect the simulation |
| | $\sigma_\phi^{noise}$ | Range of uniform perturbations due to inherent nuclear heterogeneities | 0.01 | |
| Epigenetic Kinetic Parameters | $\Gamma_{me}$ | Rate of histone methylation | 1e-4 $s^{-1}$ | Estimated from reciprocal of timescale of histone methylation reactions in literature [11] |
| | $\Gamma_{ac}/\Gamma_{me}$ | Non-dimensionalised rate of histone acetylation | 1.1 | Estimated from H2A STORM imaging in Fig 2c. Tuned for epigenetic perturbations |
| Chromatin lamina interactions | $\tilde{V}_L$ | Non-dimensionalised strength of chromatin-lamina interactions | 0.005 | Parametric study performed to best approximate the LAD distribution in absence of any perturbations |
| Mechanical constitutive properties of chromatin phases | $\tilde{G}_{EC}$ | Shear modulus of EC phase (non dimensional) | 0.025 | Chosen to reflect the effects of strain energy on chromatin coexistence (see Section S6.3-S7) |
| | $\tilde{G}_{HC}/\tilde{G}_{EC}$ | Ratio of shear modulus of HC phase to that of EC phase | 2 | Tunable |
| | $\tilde{\kappa}$ | Bulk Modulus of chromatin (non-dimensional) | $100 \times \tilde{G}_{HC}$ | High enough to enforce incompressibility constraint |
| | $\tilde{\theta}$ | Viscosity of chromatin (non-dimensional) | $1 \times \tilde{G}_{HC}$ | Tunable |

### S2 Extended experimental methods

#### *S2.1 Cell culture and immunostaining*

Bone marrow-derived hMSCs were purchased from Lonza (Cat. No. #PT-2501). hMSCs were cultured in low-glucose DMEM (Gibco; 11885) supplemented with 10% FBS (Gibco; A52568), 1% Antibiotic-Antimycotic (Gibco; 15240062), and 1% Glutamax (Gibco; 35050) at 37°C with 5% CO2. The cells were trypsinized with TrypLE (Gibco; 12604021) once they reached 70% confluency. The cells were seeded at the density of 2000 cells/cm$^2$. For immunostaining cells were fixed with 4% paraformaldehyde (PFA) in PBS for 15 min at room temperature (RT) and washed thrice with PBS. Cells were then permeabilized with permeabilizing buffer (0.5% Triton X-100 in PBS) for 10 min and blocked with bovine serum albumin (BSA) (4% bovine serum albumin in PBS) for 30 min to minimize nonspecific binding. Anti-H2A (1:50; rabbit; cell signaling,

12349S), primary antibody in 4% bovine serum albumin (BSA) were added to the samples and incubated overnight at 4°C. The primary antibodies were removed, and the samples were rinsed three times with PBS for 5 min each. The samples were then incubated at room temperature with secondary antibodies (1:50, Alexa Fluor 647, AB150079) and Hoechst 33342 (H3570) in 4% BSA for 1h at room temperature. The samples were then rinsed three times with PBS. All immunostained samples were stored in PBS at 4°C until imaging.

#### *S2.2 Nuclear morphometric parameter calculation*

Cells were imaged using a confocal microscope (LSM, Carl Zeiss) at a 63x oil immersion objective. To visualize the vertical cross-section of the nucleus, z-stacks were obtained, and 3D projections were created using Fiji. Z-stacks of confocal fluorescent images were analyzed to calculate the nuclear volume. Gaussian blur filter was applied to the z-stack to smoothen the staining pattern. Furthermore, a 3D object counter function with a fixed threshold was applied for nuclei separation in Fiji. Nuclei were segmented from Z-stacks using the nuclear edge detector function, and the nuclear volume was measured [12].

#### *S2.3 Analysis of chromatin condensation parameter (CCP)*

DAPI-stained nuclei were imaged using a laser scanning confocal microscope (LSM, Carl Zeiss) at 63X oil immersion objective. For analysis, individual nuclei were cropped using FIJI to obtain only one nucleus per image. The CCP was calculated using a MATLAB script, as previously described [13], in brief, a Sobel gradient-based edge detection algorithm was applied to detect local intensity transitions within the DAPI signal, and CCP was calculated as the density of detected chromatin edges within the nuclear area. Increased CCP values correspond to a more condensed chromatin organization, while decreased CCP values reflect a more decondensed chromatin state.

#### *S2.4 Atomic force microscopy*

AFM Measurements were performed on live cells. AFM experiments were carried out on an MFP-3D system (Asylum Research) mounted on an inverted optical microscope in contact mode force. Silicon nitride pyramidal cantilevers (TR800PB, Olympus) with spring constants 0.09 N/m and Resonance frequency ~32 kHz was used. Prior to every measurement, cantilevers were calibrated.

For nuclear stiffness measurements, the AFM tip was positioned above the central region of the nucleus using phase-contrast optics. Indentations were performed at the nuclear center force-indentation curves were acquired at a constant indentation rate of 3–5 μm/s. Measurements were completed within 10–15 min of sample removal from the cell incubator to minimize temperature-induced morphological changes. Force indentation data were analyzed using AtomicJ software [14]. Curves were fit using the Hertzian contact model for a pyramidal indenter. The apparent nuclear stiffness was reported as the Young’s modulus obtained from the fitted force-indentation curves.

#### *S2.5 Super-resolution imaging*

Super-resolution imaging-based visualization of immunostained H2A was utilized to observe the detailed meso-scale organization of chromatin in the cell nuclei. Super-resolution images were acquired using the ONI (Oxford NanoImager) commercial STORM imaging platform. The imaging system was equipped with a ×100, 1.4 numerical aperture oil-immersion objective and a sCMOS

Hamamatsu Orca Flash camera. To ensure optimal photo switching of Alexa Fluor 647, the imaging buffer followed standard guidelines (ref), consisting of 10 mM cysteamine MEA in GLOX Solution: 0.5 mg ml−1 1-glucose oxidase, 40 mg ml−1 1-catalase and 10% glucose in PBS. The 640 nm laser (1,000 mW peak power) was used at a setting of 40% to excite the reporter dye (Alexa Fluor 647). The camera's exposure settings were configured to 15 ms, and a total of 30,000 frames were collected for each image using the ONI software. Additional details can be found in (ref). The subsequent post-analysis of STORM image localization was performed using custom-written MATLAB codes (Ref)

*S2.6 FLIM-FRET imaging of live cell chromatin compaction*

Chromatin compaction was quantified in live HeLa cells expressing the histone FRET pair H2B-eGFP (donor) and H2B-mCherry (acceptor), as previously described [15]. Briefly, cells were plated onto glass-bottom dishes, transfected with H2B-eGFP and H2B-mCherry, and imaged under iso-osmotic (control), hypo-osmotic, or hyper-osmotic conditions. Chromatin decompaction was induced by hypotonic treatment, in which imaging medium was diluted with distilled water to a final ratio of 35:65 (v/v), whereas chromatin compaction was induced by supplementing the imaging medium with 160 mM sucrose. FLIM-FRET measurements were acquired on an Olympus FV3000 laser-scanning microscope coupled to an ISS A320 FastFLIM system using a 60×, 1.2 NA water-immersion objective. Cells were maintained at 37 °C in a humidified atmosphere containing 5% $CO_2$ throughout imaging. H2B-eGFP was excited using a pulsed 488 nm laser (80 MHz), and donor fluorescence was detected through a 520/50 nm emission filter using a Hamamatsu H7422P-40 photomultiplier detector. FLIM images were acquired at 256 × 256-pixel resolution with a pixel dwell time of 20 µs and 20-frame integration. Fluorescence lifetime measurements were calibrated against fluorescein (pH 9.0; lifetime = 4.04 ns). FLIM-FRET data were analysed using the phasor approach to fluorescence lifetime analysis, as previously described [16, 17]. Briefly, fluorescence lifetime maps were corrected for donor-only fluorescence and cellular autofluorescence before conversion to FRET efficiency maps. FRET efficiency maps were exported as greyscale images, with regions of high FRET efficiency corresponding to compact chromatin. Heterochromatin (HC) regions were segmented from the FRET maps by binarizing them and the fraction of nuclear area occupied by HC together with HC domain size distributions were quantified using custom image analysis routines.

*S2.7 Microfabricated static 2D cell confiner method*

To experimentally confine MDA-MB-231 breast cancer cells and IMR90 fibroblasts, cell confinement was performed using the 2D cell confiner adapted for a 6-well glass bottom dish, as previously described (Liu et al., 2015, Nader et al., 2021). The cells were confined with polydimethylsiloxane (PDMS, RT615, GE) microspacers (micropillars) that were attached to 13mm glass coverslips. The glass coverslips were plasma treated and then placed on top of a PDMS mixture (10/1 W/W PDMS A / crosslinker B) on top of wafer molds which were fabricated using standard photolithography procedures to have cylindrical holes of precise and specific depths. The depths of the holes on the wafer molds then defines the height of the micropillar on the coverslips which determines the confined space height between the coverslip and the cell substrate. After baking the coverslips on a 95°C hot plate for 20 minutes, the coverslips with the PDMS pillars were removed from the wafers using a razor blade under isopropanol. The coverslips were then cleaned with isopropanol, dried using an air gun with a filter attachment, and plasma treated. Then the coverslips were treated with 0.5 mg/mL pLL-PEG (SuSoS, PLL(20)-

g[3.5]-PEG(2)) for 1h at room temperature. The coverslips were then rinsed with water and placed in the cell culture media overnight before confining the cells.

The cells were trypsinized, resuspended at a concentration of 1.5 million cells/mL, and stained with Hoechst 33342 (ThermoFisher, R37605). Cells were plated in a 6 well glass-bottom plate. Meanwhile, the modified lid was prepared to confine the cells. Large PDMS pillars were cleaned and attached to the lid. These PDMS pillars were fabricated by pouring a PDMS mixture (A:B = 37:1) into a metallic mold, removed bubbles under a vacuum, baked overnight at 75°C, and removed from the mold with isopropanol. After attaching the confining coverslips to the large PDMS pillars, the lid was taped shut onto 6 well plate. Additional media was added to the plate.

Live cell imaging and Z-stack images were acquired on the Zeiss Airyscan 63x oil immersion super resolution mode. After Z-stacks were obtained, the images were processed with the Zeiss joint deconvolution feature to aid resolution

#### *S2.8 <u>Fabrication of micropatterned PDMS substrates</u>*

A silicon master mold with parallel line features (10 μm width, 10 μm depth, and 10 μm spacing) was custom-fabricated and supplied ready-to-use by [Microfit, Korea]. Polydimethylsiloxane (PDMS; Sylgard 184, Dow Corning) prepolymer and curing agent were mixed at a 10:1 (w/w) ratio, poured onto the master mold, degassed under vacuum to remove air bubbles, and cured at 60 °C overnight. The cured PDMS was peeled from the mold with the aid of ethanol, yielding substrates with 10 μm topographic microgrooves that replicated the master pattern. Flat PDMS control substrates were prepared from the same prepolymer mixture under identical curing conditions.

The prepared PDMS substrates were sterilized with 70% ethanol, rinsed in deionized water, and dried. Surfaces were activated by oxygen plasma treatment (CUTE, Femto Science Inc, Korea) and immediately coated with fibronectin (50 μg/mL) for 1 h at room temperature to promote cell adhesion. The substrates were then rinsed with PBS before cell seeding.

#### *S2.9 <u>Cell culture and seeding on micropatterned PDMS substrates</u>*

Human mesenchymal stem cells (hMSCs) were purchased from the American Type Culture Collection (ATCC, USA) and cultured in α-MEM (Welgene, Gyeongsan, Korea) supplemented with 10% (v/v) fetal bovine serum (FBS; Sigma-Aldrich, St. Louis, MO, USA) and 1% (v/v) penicillin/streptomycin (P/S; Thermo Fisher Scientific). For micropatterning experiments, hMSCs were detached with 0.25% trypsin-EDTA (Thermo Fisher Scientific), and seeded onto fibronectin-coated flat and 10 μm micropatterned PDMS substrates. Cells were allowed to attach for 6 h, after which additional culture medium was added. Cells were then allowed to spread and align along the microgrooves for 24 h before subsequent experiments.

### S3 Role of reaction kinetics in breaking the thermodynamic detailed balance

From a purely energetic perspective, the features of the energy landscape such as the depth of wells and the amount of acetylated and methylated histones available at the initiation of the phase separation process determine the equilibrium abundance of the chromatin species corresponding to the wells. Such energy-driven determination of HC and EC populations is called thermodynamic detailed balance and is always followed under equilibrium conditions. However, since an interconversion between these species is possible, the energetics determined populations of HC and EC can be overridden by the epigenetic regulation. In other words, if the rate of histone methylation $\Gamma_{me}$ increases, the number of methylation histones must increase leading to formation of more HC. Thus, we expect the epigenetic reaction kinetics to override the detailed balance taking the system away from thermodynamic equilibrium.

The evolution of epigenetic marks, i.e. methyl or acetyl marks, on the histones is prominently regulated by (S11), which is reproduced here,

$$\frac{\partial \Phi_d}{\partial t} = \underbrace{\nabla_K \left( M_d J H_{iK} H_{iL} \frac{\partial \mu_d}{\partial X_L} \right)}_{\text{Diffusion-like}} + \underbrace{2J(\Gamma_{me}\phi_a - \Gamma_{ac}\phi_m)}_{\text{Epigenetic Reactions}} \tag{S29}$$

From the above equation, we immediately observe that if methylation rate $\Gamma_{me}$ increases, $\phi_d$ $(= \phi_h - \phi_e)$ will increase with time, i.e. the heterochromatin content in the nucleus increases. Conversely if $\Gamma_{ac}$ increases, $\phi_d$ reduces such that the euchromatin content in the nucleus increases. Thus, qualitatively we can see that the epigenetic reaction rates regulate the amount of heterochromatin and euchromatin in the nucleus.

Next, we would like to quantify the dependence of overall heterochromatin or euchromatin levels in the nucleus on the epigenetic rates. To do so we consider the steady state of the nucleus where the domains of heterochromatin have been formed and do not evolve in time, i.e. $\frac{\partial \Phi_d}{\partial t} = 0$. Since we are interested in the overall average number of histones that are acetylated or methylated, we integrate Eq (S29) over the entire nucleus and divide by the reference volume of nucleus $V_{nuc}^R$. Therefore,

$$0 = \frac{\int_{\mathrm{V}_{nuc}^{\mathrm{R}}} \nabla_K \left( M_d J H_{iK} H_{iL} \frac{\partial \mu_d}{\partial X_L} \right) dV}{\int_{\mathrm{V}_{nuc}^{\mathrm{R}}} dV} + 2\mathrm{J}(\Gamma_{me}\bar{\phi}_a - \Gamma_{ac}\bar{\phi}_m)$$

where $\bar{\phi}_a = \frac{1}{V_{nuc}^R} \int_{V_{nuc}^R} \phi_a dV$ and $\bar{\phi}_m = \frac{1}{V_{nuc}^R} \int_{V_{nuc}^R} \phi_m dV$ are the overall volume fractions of acetylated and methylated histones in the entire nucleus, denoting the average content of acetylated and methylated histones in the nucleus. The first term in the above equation can be simplified to a surface integral of true flux $\boldsymbol{j}_d$ of epigenetic marks along the peripheral surface of the nucleus using divergence theorem. Since there is no flux of epigenetic marks across the nucleus, i.e. all epigenetic marks stay within the nucleus, the first term must be zero. Hence,

$$0 = \Gamma_{me}\bar{\phi}_a - \Gamma_{ac}\bar{\phi}_m$$

As described in Section S1.1 the volume fractions in the current configuration are related as $\phi_a = 1 - \phi_m - \phi_n$. Thus,

$$\bar{\phi}_m \approx \frac{\Gamma_{me}\,(1 - \bar{\phi}_n)}{\Gamma_{me} + \Gamma_{ac}}, \qquad \bar{\phi}_a \approx \frac{\Gamma_{ac}\,(1 - \bar{\phi}_n)}{\Gamma_{me} + \Gamma_{ac}} \tag{S30}$$

Or, in terms of rescaled quantities,

$$\bar{\phi}_m \approx \frac{\tilde{\Gamma}_{me}\,(1 - \bar{\phi}_n)}{\tilde{\Gamma}_{me} + 1}, \qquad \bar{\phi}_a \approx \frac{(1 - \bar{\phi}_n)}{\tilde{\Gamma}_{me} + 1} \tag{S31}$$

Equation (S30) presents the quantitative relation between the epigenetic reaction rates ($\Gamma_{me}$ and $\Gamma_{ac}$) and overall content of methylated and acetylated histones, and thereby the nuclear-wide average heterochromatin and euchromatin content.

Thus, our model predicts that the epigenetic reaction rates, independent of the energetic considerations determine the average heterochromatin and euchromatin level in the nucleus, thus breaking the detail balance of thermodynamic equilibrium.

### S4 Analysis of non-equilibrium phase separation around a single heterochromatin domain

The mathematical model presented in Section 1 captures the spatial reorganization of chromatin across the entire nucleus in response to chemo-mechanical cues from the environment – such as epigenetic dysregulation and applied mechanical forces. This reorganization emerges from a complex interplay between energetic factors, including chromatin-chromatin interactions and mechanical stresses and kinetic factors incorporating nucleoplasm diffusion, diffusion of epigenetic marks and post-translational histone modifications. While the model predicts chromatin organization which matches experimental observations with a high fidelity (Sections 2.2 to 2.6 of results) disentangling the individual mechanistic contributions of each of these components within the full nucleus-scale framework becomes challenging. To better understand these contributions, we introduce a simplified model: a single spherical heterochromatin domain embedded in a larger, water-rich euchromatic region (Fig S4). In this section, we derive the steady-state chromatin distribution and associated nuclear deformation for this simplified geometry of a single heterochromatin domain.

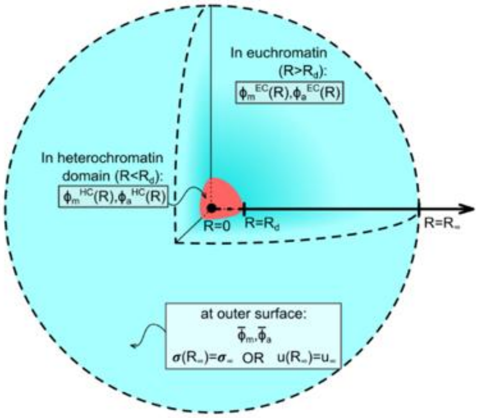

Fig S4: Phase separation of a single spherical domain of heterochromatin

We consider a single spherical heterochromatin domain (red in Fig S4) of radius $R_d$ in a much larger euchromatin background of radius $R_\infty$ (blue in Fig S4) at steady state. Assuming spherical symmetry, we want to evaluate the volume fraction of acetylated and methylated histones $\phi_a(R)$ and $\phi_m(R)$, where $R$ is a radial co-ordinate with origin $(R = 0)$ at the center of the heterochromatin domain. Additionally, to capture the local deformations associated with movement of water and epigenetic marks (see Section S1.1), we denote the radial displacement of material points located in the reference position, $R$ as $u(R)$. The mechanical forces experienced by the nucleus are transduced to chromatin within and are captured by imposing a given level of radial stress or a prescribed displacement (Fig S4) at the outer surface of euchromatin. Note that all the variables ($\phi_a(R)$, $\phi_m(R)$ and $u(R)$) are defined inside the heterochromatin domain ($R < R_d$, using superscript 'HC') as well as outside the domain in the euchromatin phase ($R > R_d$, using superscript 'EC'). Further, due to the movement of acetylated and methylated histones between the heterochromatin and euchromatin phases, the boundary between the phases (i.e., the domain radius $R_d$) is evolving in time. In other words, the steady state domain size $R_d$ itself is not known a priori and hence must be evaluated.

The axisymmetric deformation results in radial, polar and azimuthal stretches which can be written as,

$$\lambda_R = 1 + \frac{\partial u}{\partial R}, \qquad \lambda_\theta = 1 + \frac{u}{R} = \lambda_\phi, \qquad J = \lambda_R \lambda_\theta^2 \tag{S32}$$

where, $J$ is the local swelling, defined as the determinant of the deformation gradient. Note that due to spherical symmetry the polar and azimuthal stretches are the same. The rates of stretches are,

$$\dot{\lambda}_R = \frac{\partial \dot{u}}{\partial R}, \qquad \dot{\lambda}_\theta = \dot{\lambda}_\phi = \frac{\dot{u}}{R}, \qquad \dot{J} = \dot{\lambda}_R \lambda_\theta^2 + 2\lambda_\theta \dot{\lambda}_\theta \lambda_R \tag{S33}$$

Next, we consider the free energy of chromatin in terms of the energetic contributions including the elastic strain energy and the Flory-Huggins type energy of mixing of dissimilar phases (as described in Section S1.4) such that the total free energy per unit reference volume at any point is $W = W_e + W_m + P(J - 1 - \Phi_n)$. Here $P$ is a Lagrange multiplier which enforces the incompressibility constraint (Eq (S13)) at all points and corresponds physically to the osmotic pressure associated with the volumetric deformation at any point. The total free energy of chromatin inside and outside the domain is,

$$\begin{aligned}\Pi = \int_0^{R_d} 4\pi R^2 dR\big[&W_e\big(\lambda_R^{HC}, \lambda_\theta^{HC}\big) + W_m\big(J^{HC}, \Phi_d^{\mathrm{HC}}, \Phi_n^{\mathrm{HC}}\big) - \mu_n \Phi_n^{HC} - \mu_d \Phi_d^{HC} \\ &- P^{HC}\{J^{HC} - 1 - \Phi_n^{HC}\}\big] \\ &+ \int_{R_d}^{R_\infty} 4\pi R^2 dR\big[W_e\big(\lambda_R^{EC}, \lambda_\theta^{EC}\big) + W_m\big(J^{EC}, \Phi_d^{\mathrm{EC}}, \Phi_n^{EC}\big) - \mu_n \Phi_n^{EC} - \mu_d \Phi_d^{\mathrm{EC}} \\ &- P^{EC}\{J^{EC} - 1 - \Phi_n^{EC}\}\big]\end{aligned} \tag{S34}$$

Note that all energies in Eq (S34) are written in reference configuration. From Eq (S34), as the chromatin organization evolves in time the rate of change in free energy is,

$$\begin{aligned}\dot{\Pi} = \int_0^{R_d} 4\pi R^2 \Bigg[&\left(\frac{\partial W_e}{\partial \lambda_R^{HC}}\right)\dot{\lambda}_R^{HC} + 2\left(\frac{\partial W_e}{\partial \lambda_\theta^{HC}}\right)\dot{\lambda}_\theta^{HC} + \left(\frac{\partial W_m}{\partial \Phi_n^{HC}} - \mu_n + P^{HC}\right)\dot{\Phi}_n^{HC} \\ &+ \left(\frac{\partial W_m}{\partial \Phi_d^{HC}} - \mu_d\right)\dot{\Phi}_d^{HC} + \left(\frac{\partial W_m}{\partial J^{HC}} - P^{HC}\right)\dot{J}^{HC}\Bigg] dR \\ &+ \big[W_{total}(R_d^-)4\pi R_d^2 - W_{total}(R_d^+)2\pi R_d\big]\dot{R}_d \\ &+ \int_{R_d}^{R_\infty} 4\pi R^2 \Bigg[\left(\frac{\partial W_e}{\partial \lambda_R^{EC}}\right)\dot{\lambda}_R^{HC} + 2\left(\frac{\partial W_e}{\partial \lambda_\theta^{EC}}\right)\dot{\lambda}_\theta^{HC} + \left(\frac{\partial W_m}{\partial \Phi_n^{EC}} - \mu_d + P^{EC}\right)\dot{\Phi}_n^{EC} \\ &+ \left(\frac{\partial W_m}{\partial \Phi_d^{EC}} - \mu_d\right)\dot{\Phi}_d^{EC} + \left(\frac{\partial W_m}{\partial J^{EC}} - P^{EC}\right)\dot{J}^{EC}\Bigg] dR\end{aligned} \tag{S35}$$

Next, we use the stretch rates from Eq (S33),

$$\dot{\Pi} = \int_0^{R_d} 4\pi R^2 \left[\left\{\frac{\partial W_e}{\partial \lambda_R^{HC}} + \lambda_\theta^{HC^2}\left(\frac{\partial W_m}{\partial J^{HC}} - P^{HC}\right)\right\}\frac{\partial \dot{u}^{HC}}{\partial R} + 2\left\{\frac{\partial W_e}{\partial \lambda_\theta^{HC}} + \lambda_R^{HC}\lambda_\theta^{HC}\left(\frac{\partial W_m}{\partial J^{HC}} - P^{HC}\right)\right\}\frac{\dot{u}^{HC}}{R} + \left(\frac{\partial W_m}{\partial \Phi_n^{HC}} - \mu_n + P^{HC}\right)\dot{\Phi}_n^{HC} + \left(\frac{\partial W_m}{\partial \Phi_d^{HC}} - \mu_d\right)\dot{\Phi}_d^{HC}\right] dR + \left[W_{total}(R_d^-)4\pi R_d^2 - W_{total}(R_d^+)2\pi R_d\right]\dot{R}_d + \int_{R_d}^{R_\infty} 4\pi R^2 \left[\left\{\frac{\partial W_e}{\partial \lambda_R^{EC}} + \lambda_\theta^{EC^2}\left(\frac{\partial W_m}{\partial J^{EC}} - P^{EC}\right)\right\}\frac{\partial \dot{u}^{EC}}{\partial R} + 2\left\{\frac{\partial W_e}{\partial \lambda_\theta^{EC}} + 2\lambda_R^{EC}\lambda_\theta^{EC}\left(\frac{\partial W_m}{\partial J^{EC}} - P^{EC}\right)\right\}\frac{\dot{u}^{EC}}{R} + \left(\frac{\partial W_m}{\partial \Phi_n^{EC}} - \mu_d + P^{EC}\right)\dot{\Phi}_n^{EC} + \left(\frac{\partial W_m}{\partial \Phi_d^{EC}} - \mu_d\right)\dot{\Phi}_d^{EC}\right] dR \qquad \text{(S36)}$$

The derivatives of $W$ with respect to the deformation gradient (and consequently with respect to the stretches considering the spherical symmetry) give the corresponding nominal (first Piola-Kirchoff) stresses such that,

$$s_R^{HC} = \frac{\partial W_e}{\partial \lambda_R^{HC}} + \lambda_\theta^{HC^2}\left(\frac{\partial W_m}{\partial J^{HC}} - P^{HC}\right), \qquad s_\theta^{HC} = \frac{\partial W_e}{\partial \lambda_\theta^{HC}} + 2\lambda_R^{HC}\lambda_\theta^{HC}\left(\frac{\partial W_m}{\partial J^{HC}} - P^{HC}\right) \qquad \text{(S37)}$$

where, the first term is the elastic component of stress, and the next two terms contribute to the osmotic pressure. Thus, the first two terms of energy density (in blue text in Eq(S36)) inside the heterochromatin domain can be simplified using integration by parts:

$$\int_0^{R_d} 4\pi R^2 \left[s_R^{HC}\frac{\partial \dot{u}^{HC}}{\partial R} + 2 s_\theta^{HC}\frac{\dot{u}^{HC}}{R}\right] dR$$

$$= \int_0^{R_d} \left\{\frac{\partial}{\partial R}\left[4\pi R^2 s_R^{HC}\dot{u}^{HC}\right] - 4\pi R^2 \frac{\partial s_R^{HC}}{\partial R}\dot{u}^{HC} - 8\pi R s_R^{HC}\dot{u}^{HC} + 8\pi R s_\theta^{HC}\dot{u}^{HC}\right\} dR$$

(Here we have used: $\frac{\partial}{\partial R}\left[4\pi R^2 s_R^{HC}\dot{u}^{HC}\right] = 4\pi R^2 s_R^{HC}\frac{\partial \dot{u}^{HC}}{\partial R} + 4\pi R^2\frac{\partial s_R^{HC}}{\partial R}\dot{u}^{HC} + 8\pi R s_R^{HC}\dot{u}^{HC}$)

$$= 4\pi R_d^2\left[s_R^{HC}\dot{u}^{HC}\right]\Big|_{R_d} - \int_0^{R_d} 4\pi R^2\left\{\frac{\partial s_R^{HC}}{\partial R} - \frac{2\left(s_R^{HC} - s_\theta^{HC}\right)}{R}\right\}\dot{u}^{HC} dR$$

Using the integration by parts in the HC as well EC phase, we rewrite Eq(S36) as,

$$\dot{\Pi} = 4\pi R_d^2\left[\left\{s_R^{HC}(R_d^-)\dot{u}^{HC}(R_d^-) - s_R^{EC}(R_d^+)\dot{u}^{EC}(R_d^+)\right\} + \left\{W_{total}(R_d^-) - W_{total}(R_d^+)\right\}\dot{R}_d\right] + \int_0^{R_d} 4\pi R^2\left[-\left(\frac{\partial s_R^{HC}}{\partial R} - \frac{2\left(s_R^{HC} - s_\theta^{HC}\right)}{R}\right)\dot{u}^{HC} + \left(\frac{\partial W_m}{\partial \Phi_n^{HC}} - \mu_n + P^{HC}\right)\dot{\Phi}_n^{HC} + \left(\frac{\partial W_m}{\partial \Phi_d^{HC}} - \mu_d\right)\dot{\Phi}_d^{HC}\right] dR + \int_{R_d}^{R_\infty} 4\pi R^2\left[-\left(\frac{\partial s_R^{EC}}{\partial R} - \frac{2\left(s_R^{EC} - s_\theta^{EC}\right)}{R}\right)\dot{u}^{EC} + \left(\frac{\partial W_m}{\partial \Phi_n^{EC}} - \mu_n + P^{EC}\right)\dot{\Phi}_n^{EC} + \left(\frac{\partial W_m}{\partial \Phi_d^{EC}} - \mu_d\right)\dot{\Phi}_d^{EC}\right] dR \qquad \text{(S38)}$$

At steady state, the change in free energy with respect to time is zero such that $\dot{\Pi} = 0$. Thus, from Eq (S38), we obtain the following set of eight equations inside and outside the domain at the steady state:

| **HC phase (inside domain)** | **EC phase (outside domain)** | **Physical Interpretation** |
|---|---|---|
| $\frac{\partial S_R^{HC}}{\partial R} - \frac{2\left(s_R^{HC} - s_\theta^{HC}\right)}{R} = 0$ | $\frac{\partial s_R^{EC}}{\partial R} - \frac{2\left(s_R^{EC} - s_\theta^{EC}\right)}{R} = 0$ | Local balance of mechanical forces |
| $\frac{\partial W_m}{\partial \Phi_d^{HC}} = \mu_d$ | $\frac{\partial W_m}{\partial \Phi_d^{EC}} = \mu_d$ | Driving force for movement of epigenetic marks |
| $\frac{\partial W_m}{\partial \Phi_n^{HC}} + P^{HC} = \mu_n$ | $\frac{\partial W_m}{\partial \Phi_n^{EC}} + P^{EC} = \mu_n$ | Driving force for movement of water |
| $J^{HC} = \frac{1}{\phi_a^{HC} + \phi_m^{HC}}$ | $J^{EC} = \frac{1}{\phi_a^{EC} + \phi_m^{EC}}$ | Molecular incompressibility constraint |

(S39)

The eight equations shown in Eq (S39) correspond to the eight variables being solved for – $\phi_h^{HC}$, $\phi_e^{HC}$, $J^{HC}$, $P^{HC}$, $\phi_h^{EC}$, $\phi_e^{EC}$, $J^{EC}$, and $P^{EC}$. However, as $R_d$, the steady state domain radius, is also an unknown, another equation is needed to fully obtain the steady state chromatin organization.

To obtain the equation governing the size of the spherical heterochromatin domain, we begin by enforcing the following kinematic constraint at the HC/EC interface, i.e. at $R = R_d$,

$$\dot{u}^{EC}(R_d^+) = \dot{u}^{HC}(R_d^-) + \dot{R}_d\left\{\lambda_R^{HC}(R_d^-) - \lambda_R^{EC}(R_d^+)\right\} \tag{S40}$$

Physically, the kinematic constraint set by Eq (S40) implies that a point located just outside the domain moves at a velocity equal to that of a point just inside the domain plus the rate of deformation of the moving HC/EC boundary in the current configuration. Using Eq (S40) in the first term (in blue) in Eq (S38), at steady state $\left(\dot{\Pi} = 0\right)$, we obtain,

$$\left[S_R^{HC}(R_d^-) - s_R^{EC}(R_d^+)\right]\dot{u}^{HC}(R_d^-) + \left[\left\{W_{total}(R_d^-) - W_{total}(R_d^+)\right\} - \left\{\lambda_R^{HC}(R_d^-) - \lambda_R^{EC}(R_d^+)\right\}\right]\dot{R}_d = 0$$

$$\Rightarrow s_R^{HC}(R_d^-) = s_R^{EC}(R_d^+) = \frac{W_{total}(R_d^-) - W_{total}(R_d^+)}{\lambda_R^{HC}(R_d^-) - \lambda_R^{EC}(R_d^+)} \tag{S41}$$

Equation (S41) implies a balance of radial forces at the interface and provides the magnitude of the radial stress at the HC/EC boundary. Thus, Eq (S39) and (S41) offer a set of equations that simultaneously govern the steady state organization of chromatin. However, to solve them we need to specify the elastic and mixing energies such that,

$$W_e = \frac{G}{2}\left[\lambda_R^2 + \lambda_\theta^2 - 2 - 2\ln\lambda_R\lambda_\theta\right], \qquad W_m = J\phi_a^2 + J\phi_m^2(\phi_m^{\max} - \phi_m)^2 \tag{S42}$$

where, $\phi_m^{\max}$ is the volume fraction of methylated histones in the compacted heterochromatin phase and $G$ is the shear modulus which is different for the heterochromatin and euchromatin phases. As discussed in Section S1.4, we choose heterochromatin to be stiffer than euchromatin such that $G^{HC} = 2G^{EC}$. Using Eq (S37) with the specific forms of energy and mixing energies we can get the radial and tangential nominal stresses as,

$$\begin{aligned} s_R &= G\left[\lambda_R - \frac{1}{\lambda_R}\right] + \left(\frac{\partial W_m}{\partial J} - P\right)\lambda_\theta \\ s_\theta &= G\left[\lambda_\theta - \frac{1}{\lambda_\theta}\right] + \left(\frac{\partial W_m}{\partial J} - P\right)\lambda_R, \end{aligned} \tag{S43}$$

respectively. Also, we get the derivatives of $W_m$ as,

$$\frac{\partial W_m}{\partial \Phi_d} = -\phi_a + \phi_m(\phi_m^{\max} - \phi_m)(\phi_m^{\max} - 2\phi_m)$$

$$\frac{\partial W_m}{\partial \Phi_d} = -\phi_a + \phi_m(\phi_m^{\max} - \phi_m)(\phi_m^{\max} - 2\phi_m)$$

We will now individually consider the set of equations in Eq (S39) and (S41) inside and outside the domain.

Inside the heterochromatin domain:

Since the force balance equation $\frac{\partial s_R^{HC}}{\partial R} - \frac{2(s_R^{HC} - s_\theta^{HC})}{R} = 0$ must be solved by a displacement $u^{HC}(R)$ such that $u^{HC}(0) = 0$, its solution must trivially be $u^{HC}(R) = AR$, where $A$ is an unknown constant. Upon obtaining $u^{HC}(R)$, using Eq (S32) it can be immediately seen that $J$, $\lambda_R$, $\lambda_\theta$, $\phi_a$ and $\phi_m$ are independent of $R$, i.e. the chromatin distribution and the associated deformations within the domain are homogenous. Thus, inside the heterochromatin domain we have four variables $(\phi_m, \phi_a, P, A)$ and the following three equations,

$$\begin{aligned} -\phi_a^{HC} + \phi_m^{HC}(\phi_m^{\max} - \phi_m^{HC})(\phi_m^{\max} - 2\phi_m^{HC}) - \mu_d &= 0 \\ -\phi_a^{HC} - \phi_m^{HC}(\phi_m^{\max} - \phi_m^{HC})(\phi_m^{\max} - 2\phi_m^{HC}) + P^{HC} - \mu_n &= 0 \\ (1 + A)^2 - \frac{1}{\phi_a^{HC} + \phi_m^{HC}} &= 0 \end{aligned} \tag{S44}$$

Next, we consider the equations outside the domain.

Outside the heterochromatin domain:

Considering Eq (S39), we have the following four equations to be solved outside the domain $(R > R_d)$ for four variables – $u^o(R)$, $\phi_a^o(R)$, $\phi_m^o(R)$ and $P^o(R)$,

$$\begin{aligned} \frac{\partial s_R^{EC}}{\partial R} - \frac{2\left(s_R^{EC} - s_\theta^{EC}\right)}{R} &= 0 \\ -\phi_a^{EC} - \phi_m^{EC}(\phi_m^{\max} - \phi_m^{EC})(\phi_m^{\max} - 2\phi_m^{EC}) + P^{EC}(R) - \mu_n &= 0 \\ \left(1 + \frac{\partial u^{EC}}{\partial R}\right)\left(1 + \frac{u^{EC}}{R}\right) - \frac{1}{\phi_a^{EC} + \phi_m^{EC}} &= 0 \end{aligned} \tag{S45}$$

The spatial distribution of methyl-marked histones $\phi_m^{EC}(R)$ outside the heterochromatin domain is regulated by the reaction-diffusion equation discussed in Section S1.2 (Eq X), given for the spherically symmetric case at steady state as,

$$M_d \frac{\partial}{\partial R}\left[\frac{\lambda_\theta^{EC^2}}{\lambda_R^{EC}} R^2 \frac{\partial \mu_d}{\partial R}\right] + 2R^2 \lambda_R^{EC} \lambda_\theta^{EC^2} (\Gamma_{me}\phi_a^{EC} - \Gamma_{ac}\phi_m^{EC}) = 0 \quad \text{(S46)}$$

where $\mu_d = \frac{\partial W_m}{\partial \Phi_d}$. Note that the evolution equation Eq X considering the diffusion of nucleoplasm is satisfied at steady state by considering $\nabla \mu_n = 0$ (Eq S8) such that the chemical potential of water is homogenous at all points, inside as well as outside the domain. Since all the variables outside the domain are a function of $R$, here we have a set of PDEs that need to be solved simultaneously. Using Eq (S43) in the first PDE (in Eq (S45)) we obtain a second order PDE in $u^o(R)$ for the force balance at steady state. This PDE requires the following two boundary conditions on $u^o$,

$$u^o(R = R_d) = A \quad \text{(continuity of deformation at the HC/EC interface)}$$

$$\begin{matrix} u^o(R = R_\infty) = u_\infty \\ \text{OR} \\ s_R^o(R = R_\infty) = s \end{matrix} \quad \text{(Mechanical load or displacement specified)} \quad \text{(S47)}$$

As an explanation of the boundary conditions in Eq S49, note that for a second order PDE in $u^o(R)$ we need two boundary conditions. The first boundary condition in Eq S49 is a Dirichlet type boundary condition acting at the interface of the heterochromatin domain, i.e. at $R = R_d$. The second boundary condition indicates the long-range mechanical forces acting at a length scale much larger than the radius of the domain. For a single domain and the euchromatin region around it being considered in this analysis, this involves the mechanical forces generated within the nucleus. It can either involve a hydrostatic pressure $S$ acting on radially symmetrically on the intranuclear region from the surroundings or a specific displacement $u_\infty$, again radially symmetric.

The PDE from Eq (S46) is also a second order PDE in $\phi_a^{HC}(R)$ and $\phi_m^{HC}(R)$. Far away from the domain at steady state the volume fractions of acetylated and methylated histones are determined by reactions such that,

$$\phi_d^o(R_\infty) = \frac{\Gamma_{me} - \Gamma_{ac}}{\Gamma_{me} + \Gamma_{ac}}(1 - \bar{\phi}_n) \quad \text{(S48)}$$

In addition, at the interface of heterochromatin and euchromatin ($R = R_d$), the diffusive flux must be equal to the reaction driven outflux of heterochromatin,

$$4\pi R_d M_d \frac{\partial \mu_d}{\partial R}\bigg|_{R=R_d} = \frac{4\pi R_d^3}{3} \times 2(\Gamma_{ac}\phi_m^{HC} - \Gamma_{me}\phi_a^{HC}) \quad \text{(S49)}$$

Equations (S48) and (S49) together form the boundary conditions for the PDE Eq (S46).

Seven equations coming from Eq (S44), (S45) and (S46), along with boundary conditions Eq (S47), (S48), and (S49) must be solved together for nine unknown variables – $A$, $\phi_a^i$, $\phi_m^i$, $P^i$, $u^o(R)$, $\phi_a^o(R)$, $\phi_m^o(R)$, $P^o(R)$, and $R_d$. We need two more equations which can be obtained from the interfacial constraints in Eq (S41). Firstly, the continuity of radial stress at the interface $R = R_d$ for force balance at steady state gives,

$$G^{HC}\left[1+A-\frac{1}{1+A}\right]-\left[P^{HC}+\frac{\partial W_m}{\partial J}\bigg|_{\phi_a^{HC},\phi_m^{HC}}\right](1+A)$$

$$=G^{EC}\left[\left(1+\frac{\partial u^o}{\partial R}\bigg|_{R_d}\right)-\frac{1}{\left(1+\frac{\partial u^o}{\partial R}\bigg|_{R_d}\right)}\right] \tag{S50}$$

$$-\left[P^{EC}+\frac{\partial W_m}{\partial J}\bigg|_{\phi_a^{EC},\phi_m^{EC}}\right]\left(1+\frac{u^{EC}(R_d)}{R_d}\right)$$

Secondly, from the second part of Eq (S41), the over-arching constraint over the moving boundary value problem requires that,

$$G^{HC}\left[1+A-\frac{1}{1+A}\right]-s\left[P^{HC}+\frac{\partial W_m}{\partial J}\bigg|_{\phi_a^{HC},\phi_m^{HC}}\right](1+A)=\frac{W_{total}(R_d^-)-W_{total}(R_d^+)}{\lambda_R^{HC}(R_d^-)-\lambda_R^{EC}(R_d^+)} \tag{S51}$$

*Numerical solution of axisymmetric governing equations*: Solving the nine equations given by Eq (S44), (S45), (S46), (S50), and (S51) simultaneously, along with boundary conditions Eq (S47), (S48), and (S49), should give us not only the compositions of the coexisting phases, but also the size of the heterochromatin domains coexisting in euchromatin phase. It is possible to solve these equations numerically using moving boundary conditions for the HC-EC interface which determines $R_d$, as shown in Fig S5.

Here, we have indirectly solved the equations via 1D-axisymmetric phase-field system, and then we demonstrate that except at the interface the solution satisfies the governing equation described above. The 1D-axisymmetric distribution of HC and EC phases obtained as a solution is shown in Fig S5(a). Next, we plot the expressions on left hand side of the equations Eq S46 and Eq S47 in Fig S5(b). We observe that the values of these expressions are very close to zero, suggesting that the chromatin organization in Fig S5(a) satisfies the governing equations S46-S47. We note that the expressions corresponding to the incompressibility constraints $\left(J^{HC}-\frac{1}{\phi_a^{HC}+\phi_m^{HC}},\ \text{and}\, J^{HC}-\frac{1}{\phi_a^{HC}+\phi_m^{HC}}\right)$ have a value very close to zero, but not exactly zero. This arises due to the penalization-based incompressibility implemented in the phase-field solution, as opposed to exact equality. As penalization is made stricter, by increasing the bulk modulus $\kappa$, these expressions will approach closer to zero.

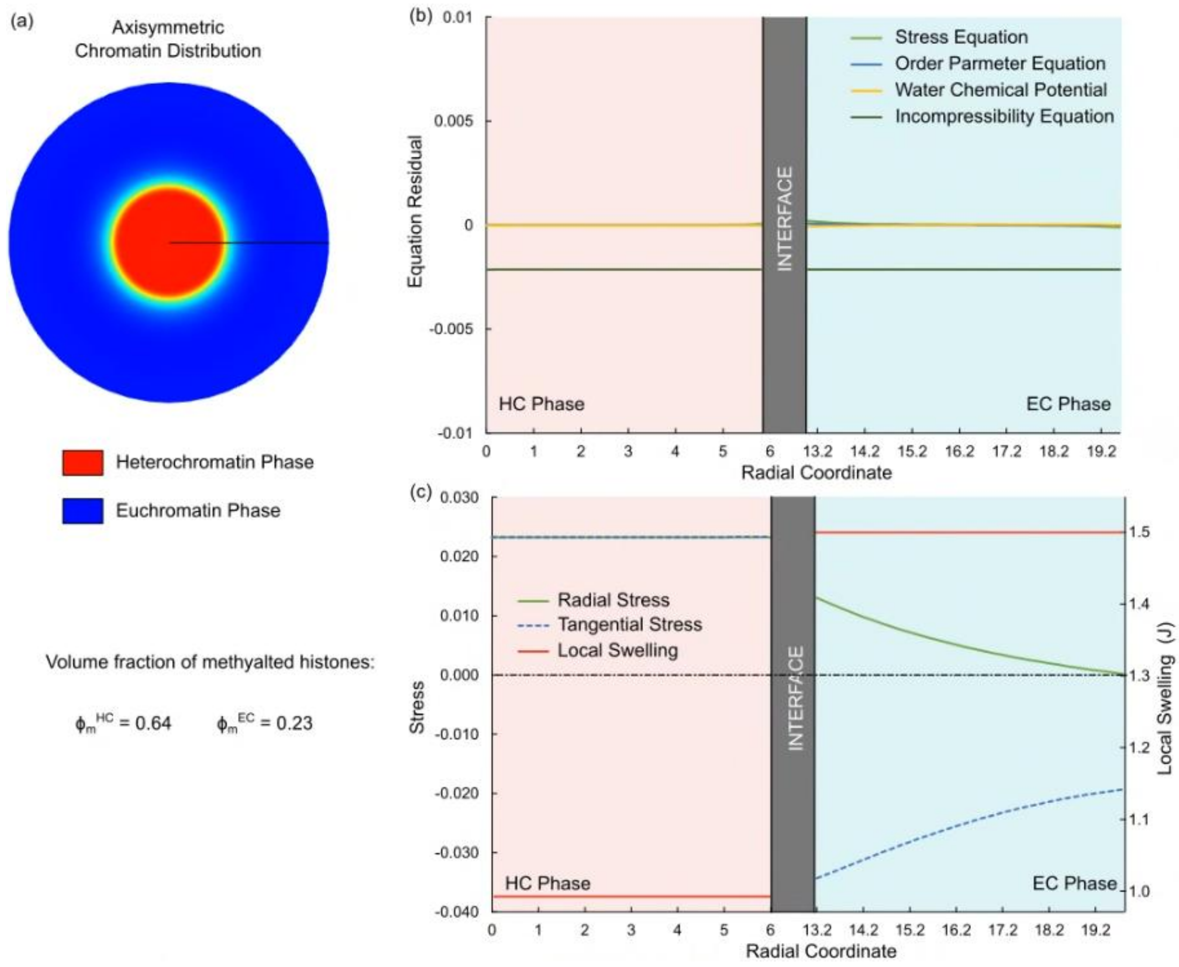

Fig S5: Numerical solution of governing equations Eq S46 – S53. (a) Axisymmetric distribution of $\phi_m$ with high methylated histone (red) region indicating HC domain surrounded by low methylated histone (blue) region indicating EC phase. The solution leads to the formation of an individual HC domain, as described in Section S3, and Fig S4. (b) The solution satisfies the governing equations, as seen by the near-zero values of the residuals at all points. Note that the residual corresponding to incompressibility equation is close to zero, but not exactly zero due to the penalty-based implementation. It will tend closer to zero in the limit $\kappa \to \infty$. (c) Plots of radial distribution of radial nominal stress $(s_{rR})$, tangential nominal stress $(s_{\theta\Theta})$ and the swelling ratio $(J = \det \boldsymbol{F})$.

Secondly, we use the solution obtained above to observe the distribution of nominal normal stress components (radial and tangential components, $s_{rR}$ and $s_{\theta\Theta}$). We note that within the HC domains, the radial and tangential stress components are exactly the same and tensile in nature. This quasi-analytically proves our statement in Section 2.1 of the main manuscript. This isotropic tensile stress exerted on the HC domain resists the expulsion of water from HC. Conversely, we note that in the EC domain the radial nominal stress is positive (tensile) while the tangential stress is negative (compression). This in turn leads to development of deviatoric stresses in the EC phase. This observation is also confirmed by our simulations, as shown in Fig S17.

### S5 Homogenous simplification for solution of the coexisting phases

Since solving the system of equations Eqs (S44) – (S51), governing the formation of a single spherical heterochromatin domain at steady state, can only be solved quasi-analytically, next we try to further decouple the problem into two sub-problems (Fig S6):

1. Thermodynamics dictated phase separation of chromatin into distinct phases – in a spatially homogenous fashion.
2. Reaction kinetics driven regulation of heterochromatin domain size – accounting for non-equilibrium reaction-diffusion driven spatial inhomogeneities.

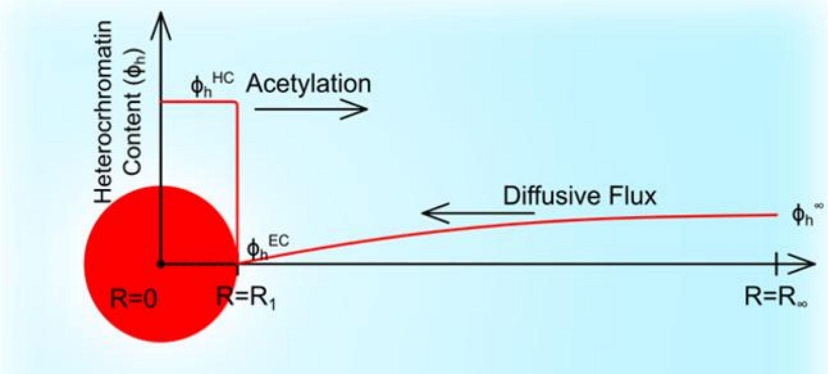

Fig S6: Schematic for a homogenous simplification of the analysis of phase separation under mechanical strain load around a single circular heterochromatin domain (red). In the bottom panel, we plot the local current heterochromatin volume fraction ($\phi_m$) as a function of radial distance from the center of the domain. Around the interface ($R \to R_d$) a sharp drop in $\phi_m$ occurs due to the thermodynamics governed separation of the coexisting phases, whereas far away ($R \to R_\infty$) the heterochromatin content is decided by the steady state reaction kinetics.

#### *S5.1* *Thermodynamics governed phase-separation*

First, we consider the energy-driven phase-separation of heterochromatin (HC) and euchromatin (EC) phases. As in the Section S4, we consider a single circular domain of HC surrounded by EC phase (as shown in Fig S4). Further, as previously, we consider that mechanical loads acting on the nucleus result in a specified radial deformation $u(R_\infty) = u_\infty$ at the outer edge.

At steady state, we are interested in evaluating the volume fractions of acetylated and methylated histones in HC phase $\phi_a^{HC}$ and $\phi_m^{HC}$, and in the EC phase $\phi_a^{HC}$ and $\phi_m^{HC}$, due to the presence of displacement load. At any point in HC or in EC phase, and at steady state, the free energy density can be written as,

$$W = W_e(J) + W_m(J, \Phi_n, \Phi_d) - P[J - 1 - \Phi_n] - \mu_n \Phi_n - \mu_d \Phi_\mathrm{d} \quad \text{(S52)}$$

where, $P$ and $J$ are the unknown, and $\mu_n$ and $\mu_d$ are given chemical potentials of water and epigenetic marks at steady-state. We note that, from the derivation in Section S4, the variables inside the HC domain are spatially homogenous, i.e. independent of $R$. However, those outside the domain vary spatially because of the reaction-diffusion flux of epigenetic marks (Eq (S46)). Since for this sub-problem, we are specifically focused on the coexistent volume fractions immediately within and outside the HC domain, we further assume that all unknown variables are spatially homogenous, i.e. independent of $R$.

Ignoring the spatially varying PDEs in Eq (S44), (S45), (S46), (S50) and (S51) i.e. the non-equilibrium effects of epigenetic reactions, we have the following equations governing the inner and outer volume fractions of acetylated and methylated histones:

| HC phase (inside domain) | EC phase (outside domain) | Physical Interpretation |
|---|---|---|
| $\sigma_p^{HC} = \frac{W^{HC} - W^{EC}}{J^{HC} - J^{EC}}$ | $\sigma_p^{EC} = \frac{W^{HC} - W^{EC}}{J^{HC} - J^{EC}}$ | Local balance of mechanical forces |
| $-\phi_a^{HC} + \phi_m^{HC}(\phi_m^{\max} - \phi_m^{HC})(\phi_m^{\max} - 2\phi_m^{HC}) = \mu_d$ | $-\phi_a^{EC} + \phi_m^{EC}(\phi_m^{\max} - \phi_m^{EC})(\phi_m^{\max} - 2\phi_m^{EC}) = \mu_d$ | Driving force for movement of epigenetic marks |
| $-\phi_a^{HC} - \phi_m^{HC}(\phi_m^{\max} - \phi_m^{HC})(\phi_m^{\max} - 2\phi_m^{HC}) + P^{HC} = \mu_w$ | $-\phi_a^{EC} - \phi_m^{EC}(\phi_m^{\max} - \phi_m^{EC})(\phi_m^{\max} - 2\phi_m^{EC}) + P^{EC} = \mu_w$ | Driving force for movement of water |
| $J^{HC} = \frac{1}{\phi_a^{HC} + \phi_m^{HC}}$ | $J^{EC} = \frac{1}{\phi_a^{EC} + \phi_m^{EC}}$ | Molecular incompressibility constraint |

(S53)

In the above equation the definition of $\sigma_p$, the Cauchy pressure (isotropic component of the force measured per unit current area), emerges as $\sigma_p = \frac{\partial W_e}{\partial J} + \frac{\partial W_m}{\partial J} - P$, analogous to Eq (S43). The Cauchy pressure is the intranuclear mechanical pressure required to result in a deformation equal to $u_\infty$ at the outer surface of the sphere denoting the euchromatin phase (Fig S6). We can solve the eight equations in Eq (S53) to get four variables $\phi_h^{HC}, \phi_e^{HC}, J^{HC}, P^{HC}$ inside the heterochromatin domain and four variables $\phi_h^{EC}, \phi_e^{EC}, J^{EC}, P^{EC}$ in the euchromatin phase.

Note that the equations in Eq (S53) can be also obtained by defining the total free energy of chromatin organization as,

$$F = W^{HC}\frac{4\pi}{3}R_d^3 + W^{EC}\frac{4\pi}{3}\left(R_\infty^3 - R_d^3\right),$$

And following the variation procedure similar to that described in Section S4,

$$\delta F = \left[\left\{\sigma_p^{HC} - \frac{W^{HC} - W^{EC}}{J^{HC} - J^{EC}}\right\}\delta J^{HC} + \left\{\frac{\partial W_m}{\partial \Phi_w^{HC}} - \mu_w + P^{HC}\right\}\delta\Phi_w^{HC} + \left\{\frac{\partial W_m}{\partial \Phi_d^{HC}} - \mu_d\right\}\delta\Phi_d^{HC}\right]\pi R_1^2 + \left[\left\{\sigma_p^{EC} - \frac{W^{HC} - W^{EC}}{J^{HC} - J^{EC}}\right\}\delta J^{EC} + \left\{\frac{\partial W_m}{\partial \Phi_w^{EC}} - \mu_w + P^{EC}\right\}\delta\Phi_w^{EC} + \left\{\frac{\partial W_m}{\partial \Phi_d^{EC}} - \mu_d\right\}\delta\Phi_d^{EC}\right]\pi\left(R_\infty^2 - R_1^2\right)$$

Setting $\delta F = 0$ for steady state we recover the equations in Eq (S53).

#### S5.2 Change in elasticity alters the thermodynamic-driven coexisting volume fractions of acetylated and methylated histones

To capture the effect of shear modulus $G$ on phase-separation, we solve (using Mathematica) Eq (S53) by varying $G$ while keeping the external mechanical strain $u_\infty$ and the chemical potentials $\mu_w$ and $\mu_d$ same. Further, since heterochromatin phase is stiffer than euchromatin (Section S1.4, Eq X), the shear modulus is different in HC and EC phases. Based on [7], we choose $G^{HC} = 2G^{EC}$.

In Fig S7, we plot the change in $\phi_m^{HC}$ and $\phi_m^{EC}$ as predicted by Eq S61 as the shear modulus $G$ changes. At $G \sim 0$ (liquid-liquid phase separation, where the elasticity of chromatin is not considered), the heterochromatin phase is prominently compacted heterochromatin ($\phi_m^{HC} = \phi_m^{max}$) while euchromatin phase has no chromatin compaction at all ($\phi_m^{EC} = 0$). However, as the strain energy contribution increases, $\phi_m^{HC}$ goes down while $\phi_m^{EC}$ increases, making the two distinct phases more and more alike. Beyond $G = 0.34$, it is seen that phase separation is completely suppressed, and the two phases are indistinct (grey hatched region in Fig S7).

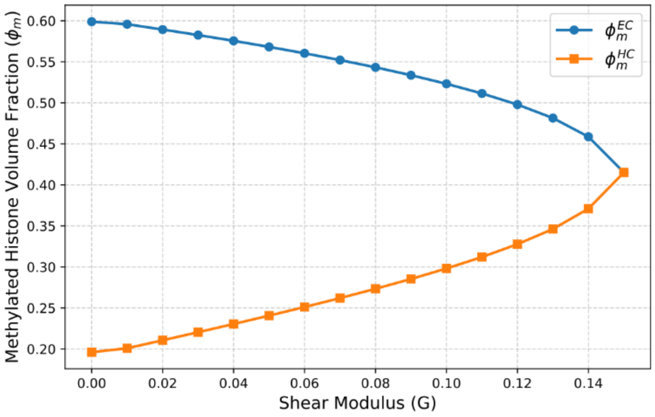

Fig S7: Local current heterochromatin volume fraction ($\phi_h$) in the heterochromatin (blue line) and euchromatin (orange line) phases changes as the effect of elasticity is increased by increasing the shear modulus $G$. Beyond an extent (shaded region) phase separation is completely suppressed.

*S5.3 <u>Reaction kinetics governed far-field composition and size regulation of HC domains</u>*

While the thermodynamic phase separation described in Section S5.1 is essential for choosing the coexisting composition of the chromatin phases, it alone is insufficient to determine the spatial aspects of chromatin organization. The spatial distribution of acetylated and methylated histones is governed by the reaction-diffusion equation (Eq (S46)) discussed in Section S4, which has not been considered due to the homogenous assumption made in Section S5.1. As discussed previously, the average amount of acetylated and methylated histones in the nucleus is determined purely by the epigenetic reactions (Eq (S30)) thereby breaking the detailed balance condition wherein at thermodynamic equilibrium the total amounts of HC and EC phase is determined by the free energies of the corresponding phases. Thus, due to the presence of epigenetic reactions the organization of chromatin is a non-equilibrium phenomenon (see Section S3) which we account for next.

Consider a circular domain of heterochromatin which phase-separates as shown in Fig S6. The volume fractions of methylated histones within the domain and immediately outside the domain are determined as $\phi_m^{HC}$ and $\phi_m^{EC}$. These values are determined by the solution Eq (S53) in Section S5.1 under a homogeneous simplification. However, far from the condensed heterochromatin domain (see Fig S6), the reaction determined average chromatin composition stays undisturbed as,

$$\phi_m^\infty = \frac{\Gamma_{me}}{\Gamma_{me} + \Gamma_{ac}}(1 - \bar{\phi}_n) \tag{S54}$$

Under the assumption of a dilute limit, where the heterochromatin domains are small and sparse enough to not interact with each other, the heterochromatin distribution plotted in Fig S6 is axisymmetric. Using a polar coordinate system with origin at the domain center, we can determine radial distribution of the heterochromatin volume fraction $\phi_m(R)$ as,

$$\phi_m(R) = \begin{cases} \phi_m^{HC} & R < R_d \quad \text{(driven by phase-separation)} \\ \phi_m^{+} = \phi_m^{EC} + \dfrac{\eta}{R} & R = R_d^{+} \quad \text{(driven by interfacial surface tension)} \\ \phi_m^{\infty} & R = \infty \quad \text{(region far from the domain stays undisturbed)} \end{cases} \tag{S55}$$

where $R_d$ is the radius of the heterochromatin domain, while $\eta$ measures the interfacial energy. Fig S6 shows the distribution of heterochromatin volume fraction $\phi_h(R)$ around a single spherical domain of heterochromatin (red) of radius $R_d$ surrounded by euchromatin phase (in blue). The gradient in the methylated histone volume fraction outside the domain will drive an inward flux of heterochromatin into the domain. The volume fraction field outside the domain at steady state must follow the reaction-diffusion equation,

$$0 = D_m \nabla^2 \phi_m - \Gamma_{ac}\phi_m + \Gamma_{me}\phi_a$$

with boundary conditions $\phi_m|_{R_d^+} = \phi_m^+$ and $\phi_m|_\infty = \phi_m^\infty$, and thus must have the form,

$$\phi_m^o(R) = \phi_m^\infty + (\phi_m^+ - \phi_m^\infty)\frac{R_d}{R} e^{\frac{R_d - R}{\ell_{RD}}} \tag{S56}$$

where $\ell_{RD}$ is the reaction-diffusion length scale defined in Section S1.6. Thus,

$$\frac{\partial \phi_m^o}{\partial R} = (\phi_m^\infty - \phi_m^+)\frac{R_d(\ell_{RD} + R)}{\ell_{RD} R^2} e^{\frac{R_d - R}{\ell_{RD}}}$$

When the heterochromatin domain is much smaller as compared to the reaction-diffusion length scale ($R_d \ll \ell_{RD}$), the we obtain,

$$\left.\frac{\partial \phi_m^o}{\partial R}\right|_{R \to R_d^+} = \frac{\phi_m^\infty - \phi_m^{EC}}{R_d} - \frac{\eta}{R_d^2}$$

The reaction-diffusion driven influx drives the growth of the heterochromatin domains such that the resulting diffusive influx of methylated histones into the domain can be written as,

$$\mathcal{J}^{in} = 4\pi R_d^2 D_m \left.\frac{\partial \phi_m}{\partial R}\right|_{R_d^+} = 4\pi R_d^2 D_m \left(\frac{\phi_m^\infty - \phi_m^{EC}}{R_d} - \frac{\eta}{R_d^2}\right)$$

Inside the domain, the methylated histone can be acetylated via epigenetic reactions. The acetylated histones are driven out due to their energetic preference for interacting with other acetylated histones. This outward flux opposes the reaction-diffusion-driven inward flux, resulting in the rate of change of volume of the HC domain, which can be written as,

$$\frac{dV_d}{dt} = \mathcal{J}^{in} - \frac{4\pi R_d^3}{3}(\Gamma_{ac}\phi_m^{HC} - \Gamma_{me}\phi_a^{HC})$$

Simplifying this, we obtain,

$$4\pi R_d^2 \frac{dR_d}{dt} = 4\pi R_d^2 D_m \left(\frac{\phi_m^\infty - \phi_m^{EC}}{R_d} - \frac{\eta}{R_d^2}\right) - \frac{4\pi R_d^3}{3}(\Gamma_{ac}\phi_m^{HC} - \Gamma_{me}\phi_a^{HC})$$

$$\Rightarrow \frac{dR_d}{dt} = D_m\left(\frac{\phi_m^{\infty} - \phi_m^{EC}}{R_d} - \frac{\eta}{R_d^2}\right) - \frac{R_d}{3}\left(\Gamma_{ac}\phi_m^{HC} - \Gamma_{me}\phi_a^{HC}\right)$$

At steady state, we set $\frac{dR_d}{dt} = 0$. Moreover, we have previously shown [10] that the effect of interfacial term $\eta$ is important in the nucleation of HC domains, but negligible at the steady state. Using these in the above equation, at the steady state, we obtain,

$$D_m\left(\frac{\phi_m^{\infty} - \phi_m^{EC}}{R_d} - \frac{\eta}{R_d^2}\right) = \frac{R_d}{3}\left(\Gamma_{ac}\phi_m^{HC} - \Gamma_{me}\phi_a^{HC}\right)$$

$$R_d = \sqrt{\frac{3D_m(\phi_m^{\infty} - \phi_m^{EC})}{(\Gamma_{ac}\phi_m^{HC} - \Gamma_{me}\phi_a^{HC})}} = \sqrt{\frac{3D_m}{(\Gamma_{ac}\phi_m^{HC} - \Gamma_{me}\phi_a^{HC})}\left\{\frac{\Gamma_{me}(1-\bar{\phi}_n)}{\Gamma_{me}+\Gamma_{ac}} - \phi_m^{EC}\right\}}$$

If the heterochromatin volume fraction immediately outside the domain is very small,

$$R_d = \sqrt{\frac{3D_m\Gamma_{me}}{\Gamma_{me}+\Gamma_{ac}}\frac{(1-\bar{\phi}_n)}{(\Gamma_{ac}\phi_m^{HC} - \Gamma_{me}\phi_a^{HC})}} \quad \text{(S57)}$$

Note however, that the heterochromatin volume fraction immediately outside the domain i.e. $\phi_m^{EC}$ may not always be negligible.

### S6 Determination of spinodal energy-well compositions from energy landscape

#### *S6.1 Triphasic liquid-like phase-separation:*

In terms of the composition of the nucleus given by $(\phi_n, \phi_d)$, the energy density arising out of chromatin-chromatin interactions is defined as,

$$W = \phi_m^2 + \phi_m^2(\phi_m^{max} - \phi_m)^2 \quad \text{(S58)}$$

where, $\phi_a = \frac{1-\phi_n-\phi_d}{2}$, and $\phi_m = \frac{1-\phi_n+\phi_d}{2}$ denote the volume fractions of acetylated and methylated histones respectively. The contour plot of $W$ in terms of $\phi_n$ and $\phi_d = \phi_m - \phi_a$ is shown in Fig S8. Note that the minima of $W$ lie at points corresponding to $(\phi_n \sim \phi_m^{max}, \phi_m = 0)$ which is the high water euchromatin phase, and $(\phi_n = 0, \phi_m = \phi_m^{max})$ which is the low water heterochromatin phase.

As discussed in Section S3, *the average contents of each phase in the overall nucleus is determined solely by the reaction kinetics*. Let the reactions choose an average heterochromatin and euchromatin composition as,

$$\bar{\phi}_m = \frac{\Gamma_{me}(1-\bar{\phi}_n)}{\Gamma_{ac}+\Gamma_{me}}, \qquad \bar{\phi}_a = \frac{\Gamma_{ac}(1-\bar{\phi}_n)}{\Gamma_{ac}+\Gamma_{me}} \quad \text{(S59)}$$

Note that these compositions depend on the initial water content $\bar{\phi}_n$ in the nucleus. Linear stability analysis can determine if any minor fluctuations from this original uniform composition will result in phase separation or not. Let there be spatially inhomogeneous random perturbations about this average composition such that,

$$\phi_a(\boldsymbol{x}) = \bar{\phi}_a + \delta\phi_a \sin(\boldsymbol{\kappa} \times \boldsymbol{x}) \quad \text{(S60)}$$

$$\phi_m(\boldsymbol{x}) = \bar{\phi}_m + \delta\phi_m \sin(\boldsymbol{\kappa} \times \boldsymbol{x})$$

The energy density $W$ gets perturbed such that the total change in the energy of a system (say, extending over a multiple of period of the sin perturbation i.e., $2\pi/\kappa$) is given as,

$$\delta\Pi = \int_0^{2\pi/\kappa} \delta W = \int_0^{2\pi/\kappa} \frac{\partial W}{\partial \phi_i}\delta\phi_i + \frac{\partial^2 W}{\partial \phi_i^2}(\delta\phi_i)^2 + \frac{\partial^3 W}{\partial \phi_i^3}(\delta\phi_i)^3 + \frac{\partial^4 W}{\partial \phi_i^4}(\delta\phi_i)^4 + 0$$

for $i = a, m$. Noting that odd powers of perturbations are negated over the entire geometry, and even orders beyond the fourth order are zero due to the definition of W (Eq (S79)), we have the perturbation in the energy as,

$$\delta\Pi = \int_0^{2\pi/\kappa} 2\delta\phi_a^2 + 2\left(6\bar{\phi}_m^2 - 6\bar{\phi}_m\phi_m^{max} + \phi_m^{max\,2}\right)\delta\phi_m^2 \tag{S61}$$

In matrix form,

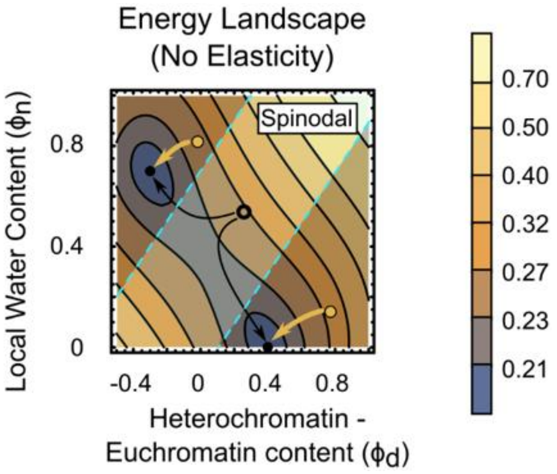

Fig S8: Two-dimensional contour plot of the free energy density $W(\phi_n, \phi_d)$. The figure also shows the spinodal region according to Eq S70. All reactions which give rise to average compositions beyond the spinodal (orange circles) do not phase separate.

$$\delta\Pi = [\delta\phi_a \quad \delta\phi_m]\begin{bmatrix}1 & 0 \\ 0 & 1\left(6\bar{\phi}_m^2 - 6\bar{\phi}_m\phi_m^{max} + \phi_m^{max\,2}\right)\end{bmatrix}\begin{bmatrix}\delta\phi_a \\ \delta\phi_m\end{bmatrix} \tag{S62}$$

Negative eigenvalues of the matrix in Eq S69 determine the unstable spinodal region, which can be written as,

$$\left(\frac{3-\sqrt{3}}{6}\right)\phi_m^{max} < \frac{1-\phi_n+\phi_d}{2} < \left(\frac{3+\sqrt{3}}{6}\right)\phi_m^{max} \tag{S63}$$

The spinodal region predicted by Eq S70 is highlighted in cyan between the dashed cyan lines in Fig S8. If the reactions determine the average composition to be outside this range (orange circles), the combined effect of kinetics and energetics would minimize the energy of the system by driving it towards the nearest minima resulting in a homogenous euchromatin or heterochromatin phase.

*S6.2 <u>Triphasic solid-like phase-separation:</u>*

Lastly, since chromatin polymer in the nucleus behaves like a solid, we extend our understanding to a solid-solid phase separation of chromatin in the nucleus. As previously, the reaction kinetics driven choice for the average euchromatin and heterochromatin compositions are determined solely by the reaction kinetics (Eq S66).

However, the energy landscape would get transformed due to the solid-like behavior of chromatin giving rise to strain energy due to any phase-transitions or evolution. The strain energy arises from the flux of volume-altering water in the distinct water-rich euchromatin phase (higher volume, $J \gg 1$) and water poor heterochromatin phase (low volume, $J \sim 1$), where $J$ denotes the change in the local volume with respect to a water-free reference state of chromatin (a water free-reference state is tightly packed heterochromatin state with no water and $J = 1$). Also note that since the local volume changes, the definitions of volume fractions change depending on whether they are calculated with respect to current (altered) or reference (original) local volumes. In this section, all volume fractions denoted via small $\phi$ are in the current configurations while all denoted as capital $\Phi$ are in reference configuration.

The reference free energy density can be written as,

$$W = J\phi_m^2 + J\phi_m^2(\phi_m^{max} - \phi_m)^2 + \frac{G}{2}(\lambda_1^2 + \lambda_2^2 - 2 - 2\ln J) \quad \text{(S64)}$$

Where, $G$ is the shear modulus and $\lambda_1$ and $\lambda_2$ are the mechanical stretches along the principal directions. Under an isotropic loading simplification, $\lambda_1 = \lambda_2 = \sqrt{J}$. Further, due to molecular incompressibility (Section S1.3), $J = \frac{1}{\phi_a + \phi_m}$.

Consider a nucleus with homogenous initial compositions $\left((\bar{\phi}_n, \bar{\phi}_d) \equiv (\bar{\phi}_a, \bar{\phi}_m)\right)$ as determined by the rates of acetylation and methylation (Eq S28). Let there be spatially inhomogeneous random perturbations about this average composition such that,

$$\begin{aligned} \phi_a(\boldsymbol{X}) &= \bar{\phi}_a + \delta\phi_a \sin(\boldsymbol{\kappa} \times \boldsymbol{X}) \\ \phi_m(\boldsymbol{X}) &= \bar{\phi}_m + \delta\phi_m \sin(\boldsymbol{\kappa} \times \boldsymbol{X}) \end{aligned} \quad \text{(S65)}$$

The resulting perturbation in the volume $J$ is $\delta J = -J^2(\delta\phi_a + \delta\phi_m)$. Thus, the alteration in the free energy density is,

$$\delta\Pi = \int_0^{2\pi/\kappa} \delta W = \int_0^{2\pi/\kappa} \left[\frac{\partial^2 W}{\partial\phi_a^2}\delta\phi_a^2 + \frac{\partial^2 W}{\partial\phi_m^2}\delta\phi_m^2 + \frac{\partial^2 W}{\partial J^2}\delta J^2\right] \quad \text{(S66)}$$

As previously, odd powered terms vanish due to periodicity as do the even powered terms beyond 4 due to the biquadratic energy density definition.

$$\delta\Pi = \int_0^{2\pi/\kappa} \left[2J\delta\phi_a^2 + 2J\left(6\bar{\phi}_m^2 - 6\bar{\phi}_m\phi_m^{max} + \phi_m^{max2}\right)\delta\phi_m^2 + \frac{G}{J^2}\delta J^2\right] \quad \text{(S67)}$$

Substituting $\delta J$,

$$\delta\Pi = \int_0^{2\pi/\kappa} \Big[2J\delta\phi_a^2 + 2J\big(6\bar{\phi}_m^2 - 6\bar{\phi}_m\phi_m^{max} + \phi_m^{max\,2}\big)\delta\phi_m^2 + J^2G\delta\phi_a^2 + J^2G\delta\phi_m^2 + 2J^2G\delta\phi_a\delta\phi_m\Big] \tag{S68}$$

Or,

$$\delta\Pi = \begin{bmatrix}\delta\phi_a & \delta\phi_m\end{bmatrix}\begin{bmatrix} J + J^2G & J^2G \\ J^2G & J\big(6\bar{\phi}_m^2 - 6\bar{\phi}_m\phi_m^{max} + \phi_m^{max\,2}\big) + J^2G\end{bmatrix}\begin{bmatrix}\delta\phi_a \\ \delta\phi_m\end{bmatrix} \tag{S69}$$

The region within which phase-separation is seen (negative eigenvalues) is given as,

$$J\big(1 + \phi_m^{max\,2} - 6\phi_m^{max}\phi_m + 6\phi_m^2\big) + 2GJ^2 - \sqrt{\big(1 - \phi_m^{max\,2} + 6\phi_m^{max}\phi_m - 6\phi_m^2\big)J^2 + 4G^2J^4} \le 0 \tag{S70}$$

where, $J = 1/(\phi_h + \phi_e)$ . On the contour plot of $W(\phi_n, \phi_d)$ in Fig S9, we have also highlighted in cyan the region dictated by Eq S77, which is identified as the spinodal region. If the reactions choose the average chromatin composition at a point within this region, we will see a phase separation. Outside the spinodal there is no phase separation.

#### *S6.3 Validation of numerical simulation using the analytically predicted spinodal*

In Section S6.2, we analytically predicted the spinodal region wherein a triphasic solid-like phase-separation of chromatin in the nucleus occurs. The analytical exercise can be used to validate the

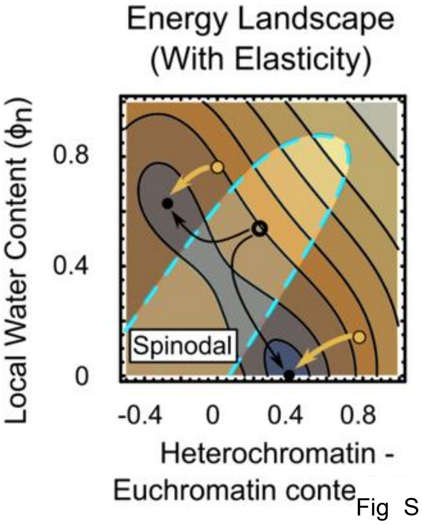

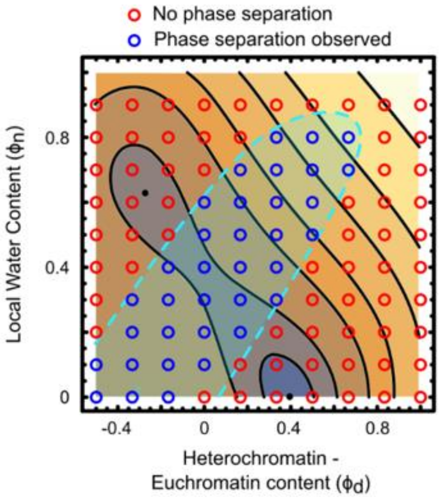

Fig S9: Two-dimensional contour plot of the free energy dens region according to Eq S77. Note that the shape of the spinod plot is created for the shear modulus $\tilde{G} = 0.04$.

Fig S10: Validation of the numerical simulations by comparison of numerically predicted (circles) and analytically predicted (contour plot) spinodal regions. The shear modulus is chosen as $\tilde{G} = 0.4$.

numerical solution of the solid phase-separation model by observing the range of average compositions over which phase-separation occurs. We numerically solve Eq S4, S8, S9, S18 and S19 over a square geometry denoting a nucleus along with boundary conditions Eq S5. In order to fix the average water content in the nucleus, we apply no-flux boundary condition for water along all the boundaries. We parametrically vary the average nucleus composition over the $(\phi_d, \phi_n)$ space.

In the $(\phi_d, \phi_n)$ space, each numerical simulation performed is denoted as a color-coded circle in Fig S10. The spinodal region predicted analytically is followed very closely by the numerical simulations as well, thus validating the numerical accuracy of our simulations with strain energy incorporated.

### S7 Effect of strain energy on phase separation and coexistence

In Sections S4-S5 of the Supplementary Information, we discussed the effects of the combined roles of chromatin-interaction energetics and strain energy in determining the phase-separation of the system. While the equations derived for this interplay were solved analytically, the energy landscapes developed in Section S6 allow us to visually interpret the individual contributions of strain energy over and above the energetics arising from chromatin-chromatin interactions. Specifically, we found that the solid-like behavior of chromatin resulted in a modified spinodal region on the chromatin energy landscape (Fig S9). Here we explore in more detail the role of mechanical strain energy in the phase separation process. Notably, the visualizations demonstrated in this section recapitulate the results emerging naturally from the equations derived in Section S4. However, these visualizations should only be interpreted qualitatively because any analysis based on energy landscapes alone lacks the non-equilibrium reaction kinetics considerations which were quantitatively incorporated in Section S4. Despite this, we still go ahead with the energy landscape analysis because of ease of visual demonstration of the interplay between mechanical strain energy and chromatin-chromatin interactions.

#### *S7.1 Higher strain energy favors heterochromatin well*

The free energy density in the reference configuration at any point in the nucleus can be written as,

$$\widetilde{W}(\phi_d, \phi_n) = \frac{\tilde{G}}{2}[2J - 2 - \log J] + J\phi_e^2 + J\phi_h^2(0.7 - \phi_h)^2 \qquad \text{(S71)}$$

Note that here we have chosen $\phi_m^{max} = 0.7$. The contour plot for the bivariable function $\widetilde{W}$ in Eq (S71) for the case if elasticity is not incorporated (i.e., set $\tilde{G} = 0$) is plotted in Fig S8. We reproduce this below in Fig S11c (leftmost panel). We can see that the energy density is a double well function which has two minima at $(-0.3, 0.7)$ and $(0.4, 0)$. As can be readily seen, the first of these minima corresponds to the water-rich euchromatin phase ($\phi_n \sim 0.7$), while the other corresponds to water-poor heterochromatin phase ($\phi_n = 0$). The co-ordinates of the two minima on the contour plot describe the constitution of the coexisting phases. With strain energy absent there is no reason why one phase should be favored over the other energetically, thus we see that both the energy wells are equally deep.

However, upon incorporating strain energy ($\tilde{G} \neq 0$) we see from Eq (S71) that the energy of each phase is different due to the presence of different amounts of nucleoplasm in each phase. A highly compact phase (the heterochromatin phase, with $\phi_n \to 0$) will have a very low swelling ($J \to 1$) as compared to the nucleoplasm free reference state. The elastic strain energy due to the low volumetric expansion is negligible. However, the euchromatin phase which has a high

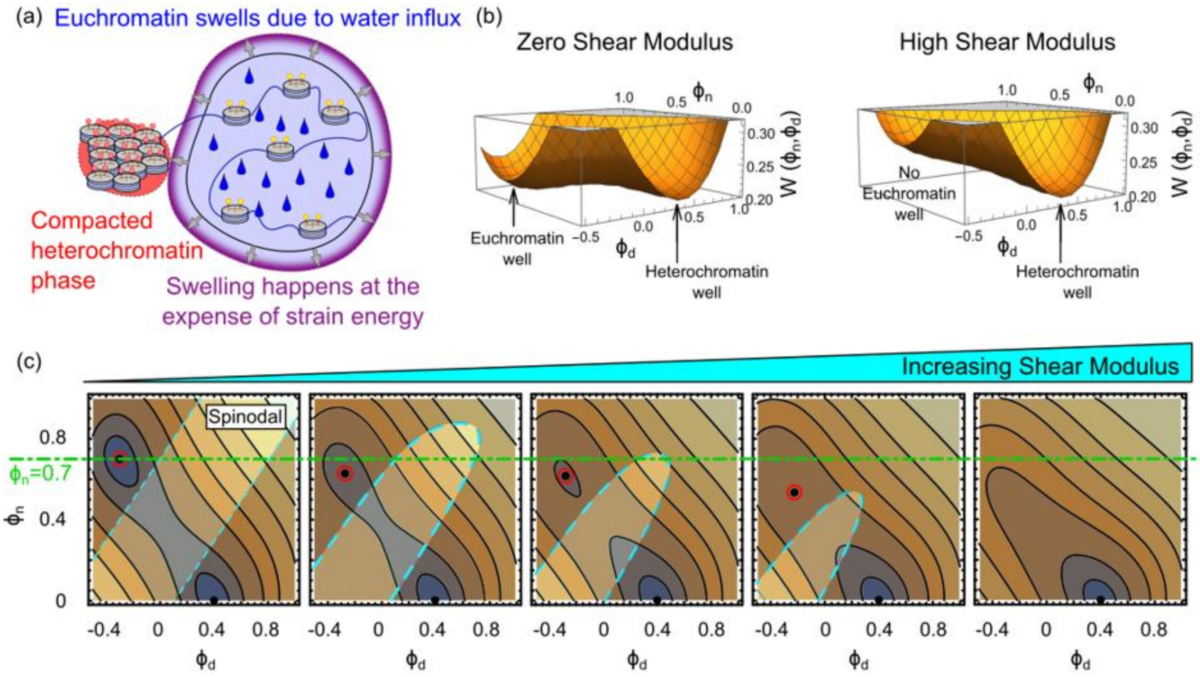

Fig S11: (a) A schematic representing the swelling of the euchromatin phase due to the presence of water. The resulting swelling causes an increase in the strain energy which makes the euchromatin phase energetically unfavorable. (b) A comparison of the 3D plots of energy density functional $\tilde{W}$ over the parametric space $(\phi_n, \phi_d)$ When shear modulus is high, the euchromatin well disappears. (c) A comparison of energy density functionals $W(\phi_n, \phi_d)$ as the effect of strain energy is increased by increasing the shear modulus of euchromatin $\tilde{G}_e$ from 0 (no effects of elasticity – liquid-liquid phase separation) to 0.2. We observe three effects – (i) the euchromatin well (encircled in red) becomes shallower until it disappears and is no longer an energetically stable state, (ii) The euchromatin well tends to approach the heterochromatin well, i.e. its composition changes as it becomes more compact ($\phi_n$ for the well decreases, while $\phi_h$ increases), and lastly, (iii) the spinodal region shrinks and gradually disappears.

nucleoplasm composition ($\phi_n \to 0.7$) will have a very high swelling due to the incompressibility constraint (Eq S10) $J = \frac{1}{1-\phi_n}$ (refer Fig S11a). Such high swelling results in an associated increase in the strain energy contribution corresponding to the first term in Eq (S71). Thus, we see that the euchromatin phase is energetically distinct from the heterochromatin phase.

As the strain energy contribution is increased (via increasing the shear modulus $\tilde{G}$), we expect the euchromatin well to increase in energy, thereby becoming less favorable as compared to the heterochromatin well. Indeed, the 3D function plot of the energy density (Fig S11b) over the parametric space $(\phi_n, \phi_d)$ show that the euchromatin well becomes shallower as the shear modulus increases. This can also be seen from the contour plots of energy density $\tilde{W}(\phi_n, \phi_d)$ as $\tilde{G}$ increases. At high values of the shear the well can completely disappear leading to loss of any phase separation behavior.

*S7.2 <u>Strain energy changes the composition of the euchromatin and heterochromatin wells:</u>*

It follows from the previous discussion that the swelling of the euchromatic phase becomes energetically unfavorable as the strain energy increases. Therefore, with increasing strain energy contribution, we would expect that the euchromatic phase swells lesser. From the incompressibility constraint (Eq S13) we know that low swelling corresponds to a smaller water content. Thus, it can be concluded that as the contribution of strain energy increases, all else being the same, the water content ($\phi_n$) corresponding to the euchromatin phase decreases.

Physically, we can imagine that there exists an energetic compensation of the expenses between the two individual contributors of bulk free energy – the strain energy ($W_e$) and the double well energy arising from the intermixing of the chromatin phases ($W_m$). In the presence of a relatively high shear modulus, the strain energy increases. However, a corresponding increase in the total energy can be avoided by adjusting the composition of the euchromatin well to be slightly distant from the euchromatic well of $W_m(\phi_n, \phi_d)$. Thus. the location of the well of the total energy density shifts on the $(\phi_n, \phi_d)$ parametric space.

This is evident on the contour plots of the energy density $\widetilde{W}(\phi_n, \phi_d)$ plotted in Fig S11c. The location of the euchromatin well on the $(\phi_n, \phi_d)$ parametric space is encircled in red. The green horizontal line shows the baseline water content of $\phi_n = 0.7$ which would correspond to the energy well in absence of elasticity $(\tilde{G} = 0)$. We see that as the shear modulus is increased, not only does the euchromatin become shallower, but it also shifts downwards, i.e. its water content $\phi_n$ decreases. Thus, the strain energy also forces a translation of the energy wells, altering the composition of the energy minimizing phases. This explains the mechanical force driven changes in the compositions of the coexisting phases as observed from our single-domain analysis in Section S4.

### S8 Simulation procedure for measuring nuclear stiffness after epigenetic perturbations

For measuring the stiffness of the nucleus, we apply a uniaxial load and measure the resulting deformation. Effective stiffness is taken as slope of the nominal stress v/s nominal strain plot under the uniaxial loading. To capture the effect of chromatin alone, we model the interior of the nucleus as shown in Fig S12a, and apply a uniaxial load as shown in Fig S12a. We apply a symmetry boundary condition for the deformations on the left and bottom boundaries ($\boldsymbol{u}.\hat{\boldsymbol{n}} = 0$).

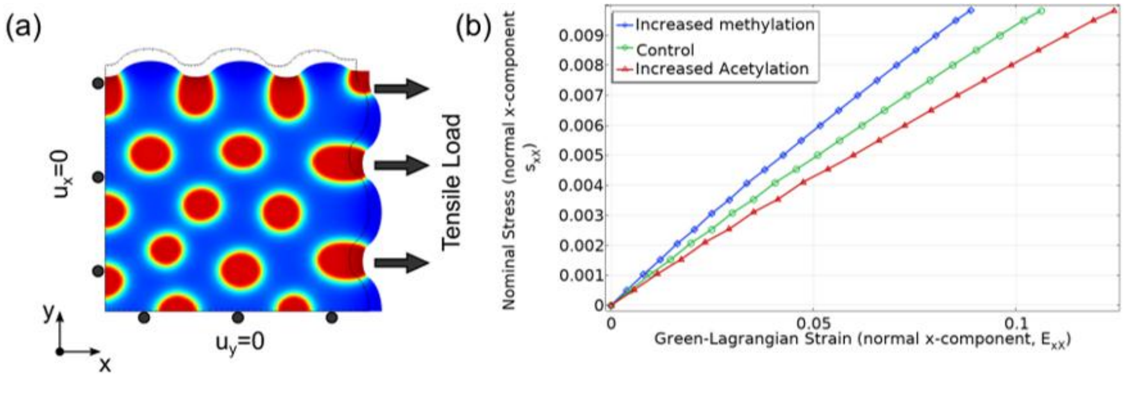

Fig S12: (a) Uniaxial tensile loads are applied to a nucleus at steady-state (black outline) and the new deformed nucleus, we reorganized chromatin is obtained. To ensure that uniaxial loads act, the left and bottom boundaries of the nucleus are treated as a roller/symmetry boundary with deformation $\boldsymbol{u}$ constrained as $\boldsymbol{u}.\hat{\boldsymbol{n}} = 0$, where $\hat{\boldsymbol{n}}$ is the outward normal to the boundary. (b) The xx-component of the nominal stress and the xx component of the Green-Lagrange strain are plotted with the slope of the curve indicating the stiffness of the nucleus.

For control nuclei, as well as nuclei under increased acetylation and methylation conditions, we plot the nominal stress (in x-direction. i.e. xx component) versus the Green-Lagrange strain (xx component, defined as $\boldsymbol{E} = 0.5 \times (\boldsymbol{F}^T\boldsymbol{F} - \boldsymbol{I})$). The effective stiffness of the nucleus is the slope of this nominal stress v/s strain curve, plotted in Fig S12b. We note that the increase in methylation (acetylation) increases (decreases) the slope of the curve, indicating an increase (decrease) in the effective mechanical stiffness of the nucleus – taking into consideration the role of chromatin organization.

### S9 Change in chromatin energy landscape in presence of mechanical loads

The change in coexistence of phases due to the presence of strain energy, which we looked at in the previous section was in absence of any external mechanical forces. We noticed that inclusion of strain energy term, via Eq (S16) alters the energy landscape of chromatin organization changing (i) depth (corresponding to energetic favorability), and (ii) location (local volume fraction of water, acetylated and methylated histones) of the heterochromatic and euchromatic phases. However, as discussed in the manuscript (Sections 2.5-2.6) application of mechanical tensile, compressive and shear forces further attenuate these changes. In this section we discuss how the energy landscape and chromatin organization is altered due to application of volumetric and isochoric (volume-conserving) shear forces.

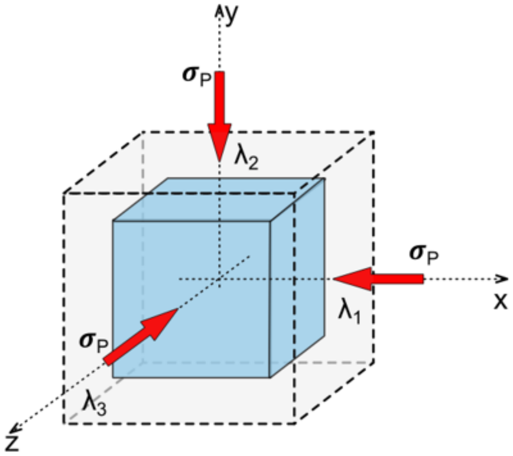

Fig S13: Volumetric load on an infinitesimal region within the nucleus.

#### *S9.1 <u>Application of volumetric loading changes the energy landscape</u>*

Consider an infinitesimal region within the nucleus, as shown in Fig S13, which experiences a triaxial mechanical stretch that is equal in all directions, i.e., $\lambda_1 = \lambda_2 = \lambda_3 = \bar{\lambda}$. Note that such is purely volumetric and has no shear component. The resulting change in the volume of this infinitesimal region is given as $J = \bar{\lambda}^3$. Say the local volume fraction of nucleoplasm, acetylated and methylated histones are $\Phi_n$, $\Phi_a$ and $\Phi_m$ with respect to the reference configuration. In terms of volume fractions of the nuclear constituents and the volumetric deformation $J$, the free energy per unit infinitesimal volume of the region can be written as,

$$W = \underbrace{\frac{G}{2}[2J - 2 - 2\ln J]}_{\text{elastic strain energy}} + \underbrace{J[\phi_a^2 + \phi_m^2(0.7 - \phi_m)^2]}_{\text{chromatin-chromatin interactions}} - \underbrace{\mu_d\Phi_d - \mu_n\Phi_n}_{\substack{\text{work done by}\\ \text{chemical potentials}}} - \underbrace{\sigma_p(J-1)}_{\substack{\text{work done}\\ \text{mechanical stress}}} - \underbrace{P(J - 1 - \Phi_n)}_{\substack{\text{Penalty enforcing}\\ \text{incompressibility constraint}}} \quad \text{(S72)}$$

Here, $\sigma_p$ is the mechanical stress acting at the boundaries of the infinitesimal region being considered, which results in the stretch $\bar{\lambda}$. The free energy density can be rewritten in terms of two independent variables - $\phi_n$ and $\phi_d$ as,

$$W = \frac{G}{2}\left[\frac{2}{1-\phi_n} - 2 - 2\ln\left(\frac{1}{1-\phi_n}\right)\right] + \frac{1}{1-\phi_n}\left[\left(\frac{1-\phi_n-\phi_d}{2}\right)^2 + \left(\frac{1-\phi_n+\phi_d}{2}\right)\left(0.7 - \frac{1-\phi_n+\phi_d}{2}\right)^2\right] - \frac{\mu_d\phi_d}{1-\phi_n} - \frac{\mu_n\phi_n}{1-\phi_n} - \frac{\sigma_p\phi_n}{1-\phi_n} \quad \text{(S73)}$$

where, we have used the incompressibility constraint $J = 1/(1-\phi_n)$ (See Section S1.3) and redefined all variables in terms of $\phi_n$ and $\phi_d$. Thus, Eq (S73) predicts how the local volume fraction of the local nuclear constituents change due to the presence of a volumetric triaxial stress $\sigma_p$. We use Eq (S73) to plot the energy landscape in presence of varying tensile and compressive triaxial stresses (Fig S14). As discussed in Section 2.5 of the main manuscript, under tensile loads the volume of the nucleus will increase, which is accompanied by a water influx. The higher presence of water will then lead to a preference for the water-rich loosely packed euchromatin phase. Thus, we see from Fig S14 that under tensile loads the euchromatin well (denoted as blue dots) become deeper (more blue) and the model predicts a higher favorability towards euchromatic phase. On the other hand, compressive loads result in a shallower euchromatin well (less blue), favoring the existence of more compacted heterochromatin phase.

In addition to changing the energy landscape visualized as the contours in Fig S14, we also see

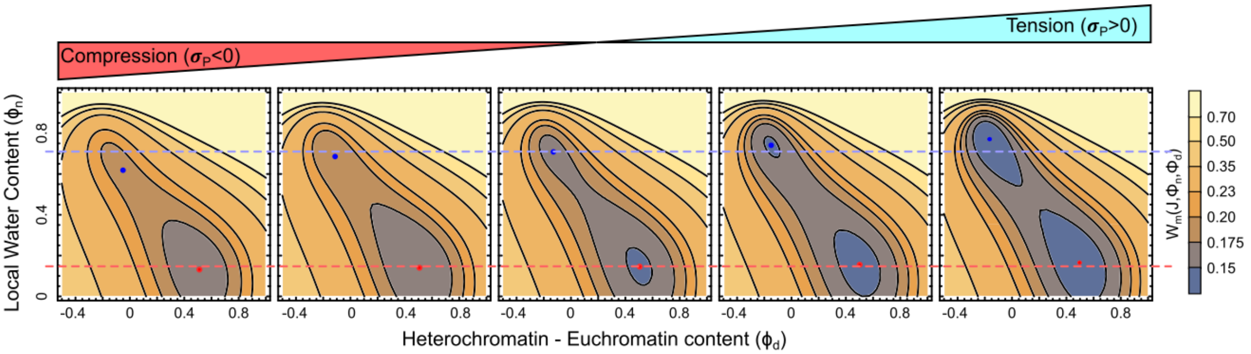

Fig S14: Change in the energy landscape of chromatin organization upon application of tensile and compressive volumetric forces.

that the locations of the energy wells change. Physically this means that the local volume fractions of water, acetylated and methylated histones $(\phi_n, \phi_a, \phi_h)$ that correspond to the energy-minima stable phase of chromatin are also determined by the application of forces. Compressive forces lead to nucleus squeezing and water efflux. As a result, the volume fraction of nucleoplasm $\phi_n$ in the euchromatin phase decreases – making the euchromatin phase relatively more chromatin-rich. On the other hand, under tensile loads water influx into the nucleus results in the euchromatin phase with a higher volume fraction of water – making it more water-rich. Note that while the euchromatin well (blue dots) translates significantly, the heterochromatin well (red dot) does not move much upon application of volumetric loads. This is because the heterochromatin phase already has a very less water $(\phi_n \ll \phi_h, \phi_e)$, so change in water content in the nucleus does not significantly perturb this well.

In conclusion, our theory demonstrates mechanistically how application of tensile and compressive loads can alter the energy landscape, thereby driving a chromatin reorganization.

Although the local volume fractions of water, acetylated and methylated histones in the two stable phases (locations of the energy wells in Fig S14) demonstrate the qualitative trends, their quantitative values are not accurate. This is because when calculating the location of the energy wells, we are assuming that a state of thermodynamic minima is reached. In other words, the wells denote the composition if energy minimum is attained at each specific point in the nucleus. However, as discussed in Section S3, chromatin organization is a far from equilibrium process where diffusion-reaction kinetics plays a significant role. Thus, the chromatin evolution occurs so as to reduce the overall free energy of the entire nucleus rather than every individual point.

*S9.2 <u>Chromatin reorganization driven by application of shear loads</u>*

In the previous subsection we have derived how volume altering mechanical forces change (i) the composition of the coexisting phases, and (ii) their steady-state sizes. In this section we study the role of volume-conserving simple shear stress.

As previously, we consider a single domain of heterochromatin with the neighboring euchromatin region. For analytical simplicity with shear loading, we consider a geometry as shown in Fig S15.

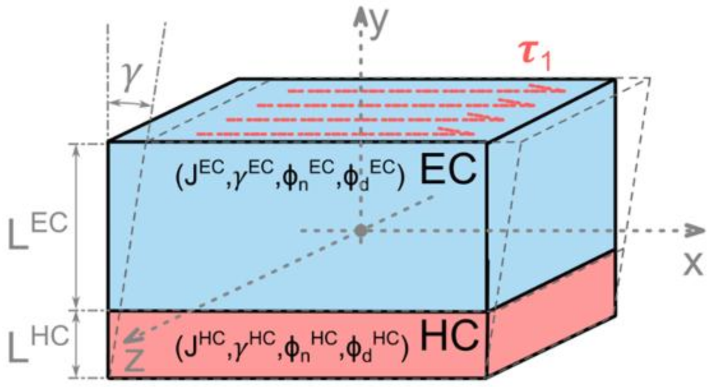

Fig S15: The two phases of chromatin under a simple shear load.

We assume that no external pressure (volumetric stress) acts on the geometry, such that it is free to swell or shrink isotropically in the three dimensions. Let us assume that the volume fraction of nucleoplasm, acetylated and methylated histones in the EC and HC phase are respectively: $(\phi_n^{EC}, \phi_d^{EC} = \phi_m^{EC} - \phi_a^{EC})$ and $(\phi_n^{HC}, \phi_d^{HC} = \phi_m^{HC} - \phi_a^{HC})$. The volumetric deformation of each phase is $J^{EC}$ and $J^{HC}$, such that the EC phase contains more water and swells more than the HC phase. At the interface of the two phases, we are ignoring the non-homogeneity that arises due to the mismatched volumetric deformation. Under such a free swelling condition, in absence of any external mechanical load, the sizes of the EC and HC phases are $L^{EC}$ and $L^{HC}$, respectively, such that the total size of the geometry $L = L^{HC} + L^{EC}$.

Next, we apply a simple shear stress $\tau_1$ along the x-axis such that the overall shear strain is $\gamma$. However, the unlike phases may not deform in a similar manner. Thus, the shear strain in the EC and HC region are $\gamma^{EC}$ and $\gamma^{HC}$ respectively. The deformed size of the geometry is,

$$\gamma^{EC} L^{EC} + \gamma^{HC}(L - L^{EC}) = \gamma L \quad \text{(S74)}$$

Assuming that all these variables are spatially homogeneous in their respective phases, and ignoring any heterogeneities at the interface, the free energy density at any point is,

$$W = W_e(J, \gamma) + W_m(J, \Phi_n, \Phi_d) - P(J - 1 - \Phi_n) - \mu_n \Phi_n - \mu_d \Phi_d \quad \text{(S75)}$$

The total free energy of chromatin in the entire geometry can be written as,

$$\Pi = AL^{EC} W^{EC} + AL^{HC} W^{HC} \quad \text{(S76)}$$

where, $W^{HC}$ is the energy density according to Eq S57 in the heterochromatin domain, while $W^{EC}$is that in the euchromatin region.

As a result of application of shear force $\tau$, let us say that the composition and volume of each phase changes. The resulting change in the free energy density in each phase is:

$$\delta \mathrm{W}^{\mathrm{EC}} = \left(\frac{\partial W_e}{\partial J^{EC}} + \frac{\partial W_m}{\partial J^{EC}} - P^{EC}\right)\delta J^{EC} + \left(\frac{\partial W_m}{\partial \Phi_n^{EC}} - \mu_n + P^{EC}\right)\delta \Phi_n^{EC} + \left(\frac{\partial W_m}{\partial \Phi_d^{EC}} - \mu_d\right)\delta \Phi_{\mathrm{d}}^{\mathrm{EC}} + \frac{\partial W_e}{\partial \gamma^{EC}}\delta \gamma^{EC}$$

$$\delta \mathrm{W}^{\mathrm{HC}} = \left(\frac{\partial W_e}{\partial J^{HC}} + \frac{\partial W_m}{\partial J^{HC}} - P^{HC}\right)\delta J^{HC} + \left(\frac{\partial W_m}{\partial \Phi_n^{HC}} - \mu_n + P^{HC}\right)\delta \Phi_n^{HC} + \left(\frac{\partial W_m}{\partial \Phi_d^{HC}} - \mu_d\right)\delta \Phi_{\mathrm{d}}^{\mathrm{HC}} + \frac{\partial W_e}{\partial \gamma^{HC}}\delta \gamma^{HC} \quad \text{(S77)}$$

Similarly, from Eq (S74), we obtain,

$$\delta \gamma^{EC} L^{EC} + \delta \gamma^{HC} L^{HC} + \delta L^{EC}(\gamma^{EC} - \gamma^{HC}) = 0$$

$$\Rightarrow \delta L^{EC} = \frac{-\delta \gamma^{EC} L^{EC} - \delta \gamma^{HC} L^{HC}}{(\gamma^{EC} - \gamma^{HC})} = -\delta L^{HC} \quad \text{(S78)}$$

The change in the total free energy due to the shear deformation is obtained from the variation of Eq (S76) as,

$$\frac{\delta \Pi}{\mathrm{A}} = \delta L^{EC} W^{EC} + L^{EC}\delta W^{EC} + \delta L^{HC} W^{HC} + L^{HC}\delta W^{HC}$$

$$\Rightarrow \frac{\delta \Pi}{\mathrm{A}} = \delta L^{EC}(W^{EC} - W^{HC}) + L^{EC}\delta W^{EC} + L^{HC}\delta W^{HC} \quad \text{(S79)}$$

We use Eq (S77) and (S78) in Eq (S79) to obtain,

$$\frac{\delta \Pi}{A} = \left(\overbrace{\frac{\partial W_e}{\partial \gamma^{EC}}}^{\tau^{EC}} - \frac{W^{EC} - W^{HC}}{\gamma^{EC} - \gamma^{HC}}\right)\delta \gamma^{EC} L^{EC} + \left(\overbrace{\frac{\partial W_e}{\partial \gamma^{HC}}}^{\tau^{HC}} - \frac{W^{EC} - W^{HC}}{\gamma^{EC} - \gamma^{HC}}\right)\delta \gamma^{HC} L^{HC} + L^{EC}\left[\left(\frac{\partial W_e}{\partial J^{EC}} + \frac{\partial W_m}{\partial J^{EC}} - P^{EC}\right)\delta J^{EC} + \left(\frac{\partial W_m}{\partial \Phi_n^{EC}} - \mu_n + P^{EC}\right)\delta \Phi_n^{EC} + \left(\frac{\partial W_m}{\partial \Phi_d^{EC}} - \mu_d\right)\delta \Phi_{\mathrm{d}}^{\mathrm{EC}}\right] + L^{HC}\left[\left(\frac{\partial W_e}{\partial J^{HC}} + \frac{\partial W_m}{\partial J^{HC}} - P^{HC}\right)\delta J^{HC} + \left(\frac{\partial W_m}{\partial \Phi_n^{HC}} - \mu_n + P^{HC}\right)\delta \Phi_n^{HC} + \left(\frac{\partial W_m}{\partial \Phi_d^{HC}} - \mu_d\right)\delta \Phi_{\mathrm{d}}^{\mathrm{HC}}\right]$$

Note that the terms in red in the above equation define the Cauchy shear stress in the EC and HC phase such that $\tau^{EC} = \partial W_e/\partial \gamma^{EC}$, and $\tau^{HC} = \partial W_e/\partial \gamma^{HC}$. At steady state, $\delta \Pi = 0$, giving the following equations to be solved simultaneously to obtain the steady state compositions and stresses in the EC and HC phase,

| EC Phase | HC Phase |
|---|---|
| $\frac{\partial W_e}{\partial J^{EC}} + \frac{\partial W_m}{\partial J^{EC}} - P^{EC} = 0$ | $\frac{\partial W_e}{\partial J^{HC}} + \frac{\partial W_m}{\partial J^{HC}} - P^{HC} = 0$ |
| $\frac{\partial W_m}{\partial \Phi_n^{EC}} - \mu_n + P^{EC} = 0$ | $\frac{\partial W_m}{\partial \Phi_n^{HC}} - \mu_n + P^{HC} = 0$ |
| $\frac{\partial W_m}{\partial \Phi_d^{EC}} - \mu_d = 0$ | $\frac{\partial W_m}{\partial \Phi_d^{HC}} - \mu_d = 0$ |
| $J^{EC} = 1 + \Phi_{\mathrm{n}}^{\mathrm{EC}}$ | $J^{HC} = 1 + \Phi_n^{HC}$ |
| $\tau = \tau^{HC} = \tau^{EC} = \frac{W^{EC} - W^{HC}}{\gamma^{EC} - \gamma^{HC}}$ | |

(S80)

The last equation in Eq (S80) gives the force balance between the two phases such that the Cauchy stress in the two phases are $\tau^{EC} = \tau^{HC} = \frac{W^{EC}-W^{HC}}{\gamma^{EC}-\gamma^{HC}}$. If a shear strain $\gamma$ on the geometry is known, the shear strain in each phase is obtained in terms of the Cauchy stress as $\gamma^{HC} = \tau/G^{HC}$ and $\gamma^{EC} = \tau/G^{EC}$. Substituting these in the last equation (in red) in Eq (S80), we obtain two equations which can be solved simultaneously for two variables - $\gamma^{EC}$ and $\gamma^{HC}$.

Note that the last equation in red in Eq (S80) is decoupled from the other eight equations. Thus the first eight equations in Eq (S80) can be solved simultaneously to obtain the following eight variables - $(\phi_h^{EC}, \phi_e^{EC}, J^{EC}, P^{EC})$ and $(\phi_h^{HC}, \phi_e^{HC}, J^{HC}, P^{HC})$. Note that these eight equations do not involve the shear stress $\tau_1$ or the shear strain $\gamma$ applied to the geometry. Thus, this simplified analysis suggests that the application of volume conserving shear loads does not alter the volume $J$ of either phase, and therefore do not influence the composition of the two phases. This result is consistent with our numerical observations from the nucleus-wide model where application of simple shear as well as pure shear does not alter the volume fractions of nucleoplasm, acetylated or methylated histones in both the heterochromatin and the euchromatin phases (Section 2.6 in the main manuscript). Thus, we predict that while shear forces give rise to a distortional deformation (i.e. change in the shape of HC domains), there is no dilational deformation (i.e. no change in volumes of HC, EC phases or that of the entire nucleus.)

### S10 Experimental validation of how substrate stiffness regulates chromatin condensation and nuclear volume via epigenetic control

In Section 2.2 of the manuscript, we showed how epigenetic modifications induced pharmacologically in bone marrow-derived human mesenchymal stem cells (hMSCs) alter HC domain sizes, thereby quantitatively validating our model predictions. Beyond pharmacological perturbations, epigenetic activity can also be indirectly modulated biophysically through changes in cell contractility, which influence the nucleo-cytoplasmic shuttling of epigenetic regulators such as HDAC3 and EZH2 [18, 19]. For example, in stiff tissue environments with an elevated cell contractility [20], HDAC3 is exported from the nucleus to the cytosol [21]. Conversely, when contractility is pharmacologically inhibited, HDAC3 accumulates in the nucleus [21, 22]. To further test our model's predictions with such upstream biophysical perturbations, we cultured hMSCs on soft (3 kPa) and stiff (30 kPa) polyacrylamide (PAA) hydrogel substrates and used DAPI staining with confocal imaging to quantify chromatin condensation and nuclear volume (Fig S16a). Cells on soft substrates exhibited greater chromatin condensation, with chromatin compaction parameter (CCP) approximately 1.4 times higher than that of cells on stiff substrates (Fig S16b), in agreement with model predictions (Fig 2a) that increased histone methylation on soft substrates enhances chromatin condensation. Additionally, cells on stiff substrates displayed an average nuclear volume 1.6 times larger than those on soft substrates (Fig S16c), further supporting our prediction that chromatin-phase transitions, driven by epigenetic regulation, mediate nuclear volume changes. While previous studies have attributed nucleus volume changes to compressive forces exerted by perinuclear actin filaments [22], studies have estimated that in non-muscle mammalian cells cytoskeletal stress exerted on the nucleus is at most of the order of 1 kPa [23]. On the other hand, our model identifies an alternative cell-signaling driven nucleus volume regulation – epigenetically driven chromatin reorganization alters osmotic pressure within the nucleus, which in turn regulates water exchange with the cytosol and nuclear volume. Indeed, osmotic pressures have been shown to result in stresses of the order of 100 kPa to 1 MPa [23] in the nuclear envelope which is enough to drive a net water flux and hence nuclear volume changes.

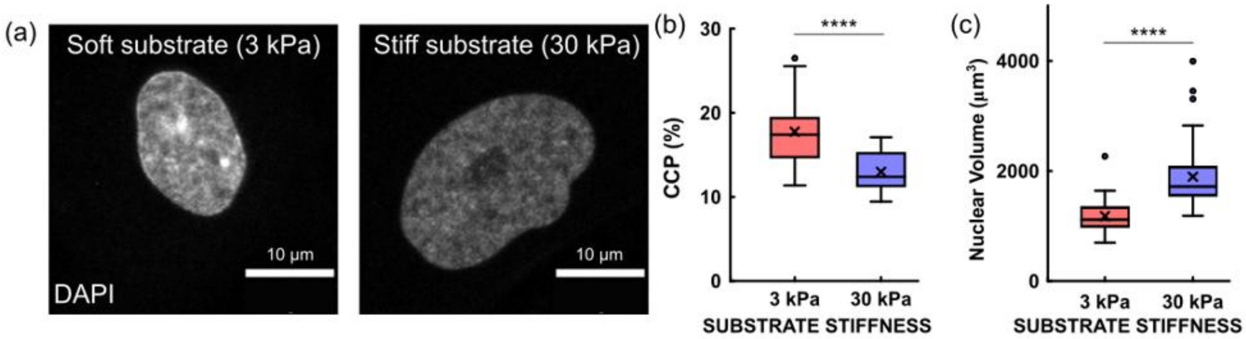

Fig S16: (a) Confocal images of nuclei in cells cultured on soft (3 kPa) and stiff (30 kPa) substrates (n=35 nuclei for each condition). All scale bars: 10 $\mu$m. (b) Quantification of chromatin condensation parameter (CCP) normalized to the mean CCP on stiff condition (n= 35 nuclei, unpaired two-tailed t-test, $p=4.4\times10^{-8}$). (c) Quantification of nuclear volume normalized to the mean value on soft substrate (n=35 nuclei, unpaired two-tailed t-test, $p=1.2\times10^{-8}$).

### S11 Additional Figures

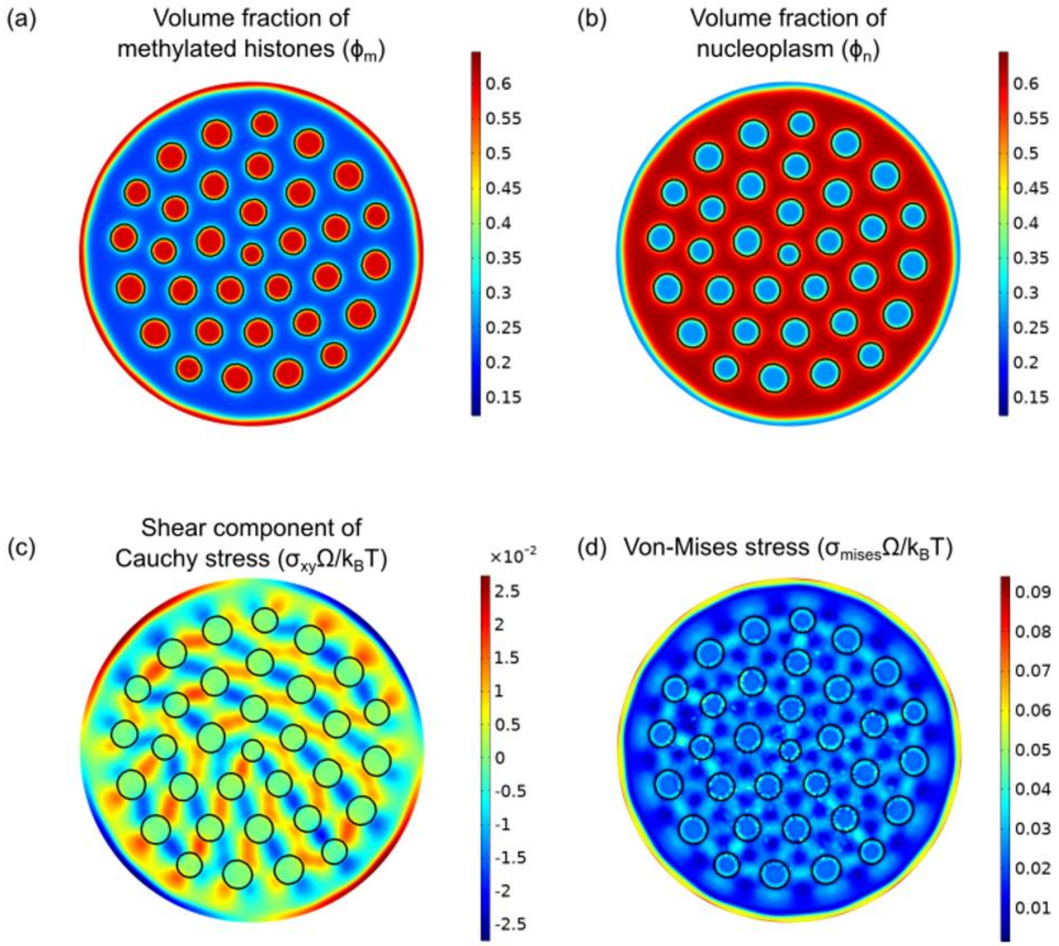

Fig S17: Simulation figures visualizing the steady-state spatial distribution of (a) the volume fraction of methylated histones $\phi_m$, (b) volume fraction of water $\phi_n$, (c) Normalized Cauchy shear stress, and (d) normalized Von-mises stress. The red regions in (a) denote the heterochromatin domains, which are also outlined in black in all the figures. Figure (b) shows that the EC phase is rich in water. Using figures (c) and (d), we visualize the presence of distortional stresses within the EC phase, especially close to HC domains.

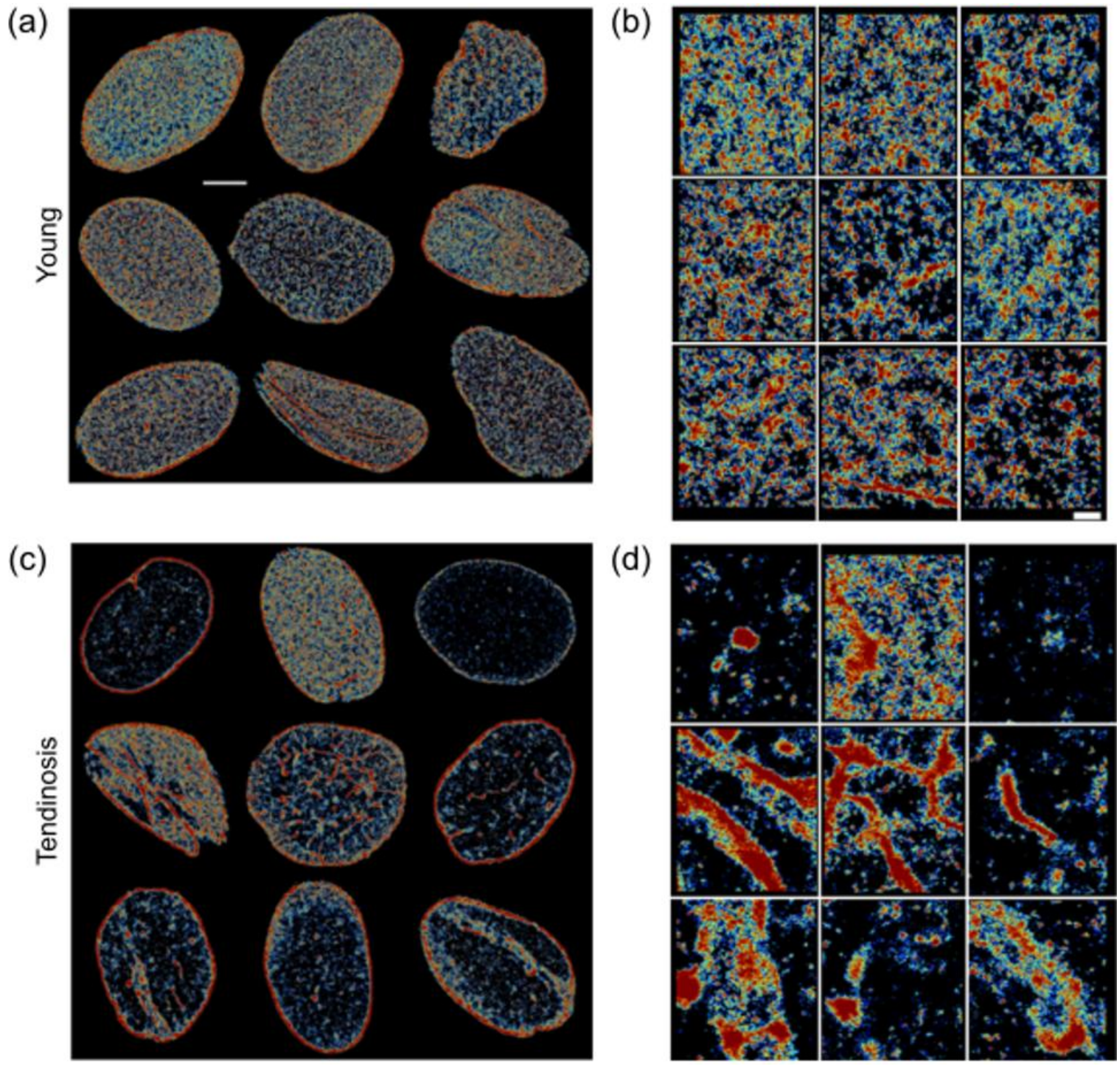

Fig S18: Mosaic representations of STORM super-resolution images reproduced from Supplementary Fig 2 in Heo, et al. (2022) [24]. Whole nucleus (left column), and zoomed in interior regions (right column) of histone H2B in human tenocytes (derived from young, healthy (top row), and young tendinosis (bottom row) patients) rendered as a Voronoi density map. Histone methyltransferase EZH2 (enhancer of zeste homolog 2), which is known to be over expressed in the nucleus in a mechanical soft tissue environment [24], leads to increased chromatin condensation in diseased nuclei, particularly resulting in the formation of long lamellar heterochromatin domains, as predicted by the simulation (hypermethylation condition in Fig 2a, main manuscript).

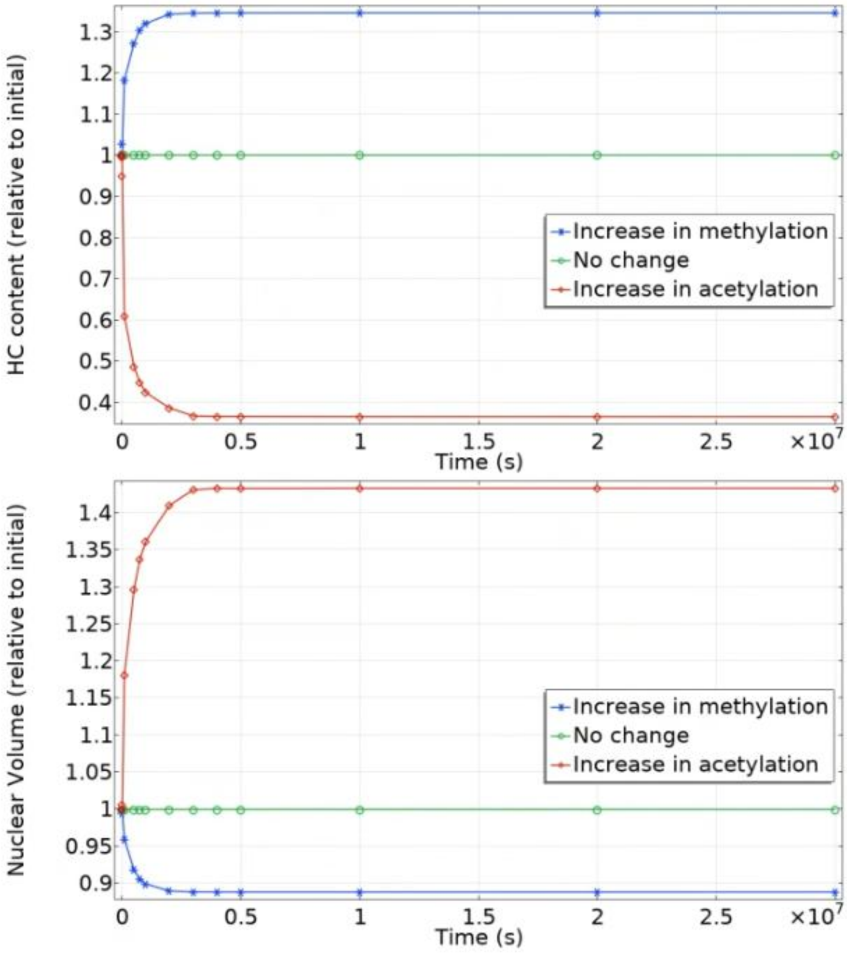

Fig S19: Change in HC content (defined as the percent of nuclear area occupied by HC) and the change in the nuclear volume over time as the rate of overall methylation or acetylation is increased. All quantities are normalized to their initial values.

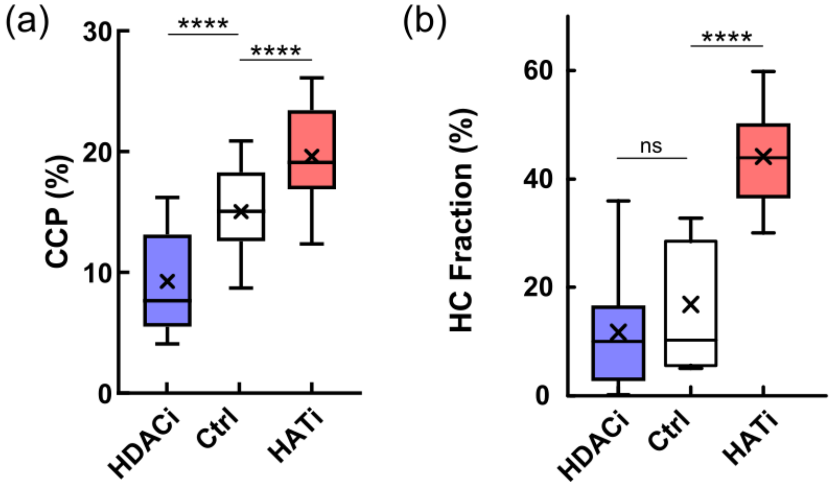

Fig S20: (Refers to Fig 2e,f in the main manuscript) Quantification of chromatin reorganization in hMSC nuclei subjected to HAT inhibition (anacardic acid), control and HDAC inhibition (valproic acid) treatments. The reorganization is quantified through (a) chromatin condensation parameter (CCP), and (b) percentage of volume of nucleus occupied by the heterochromatin phase.

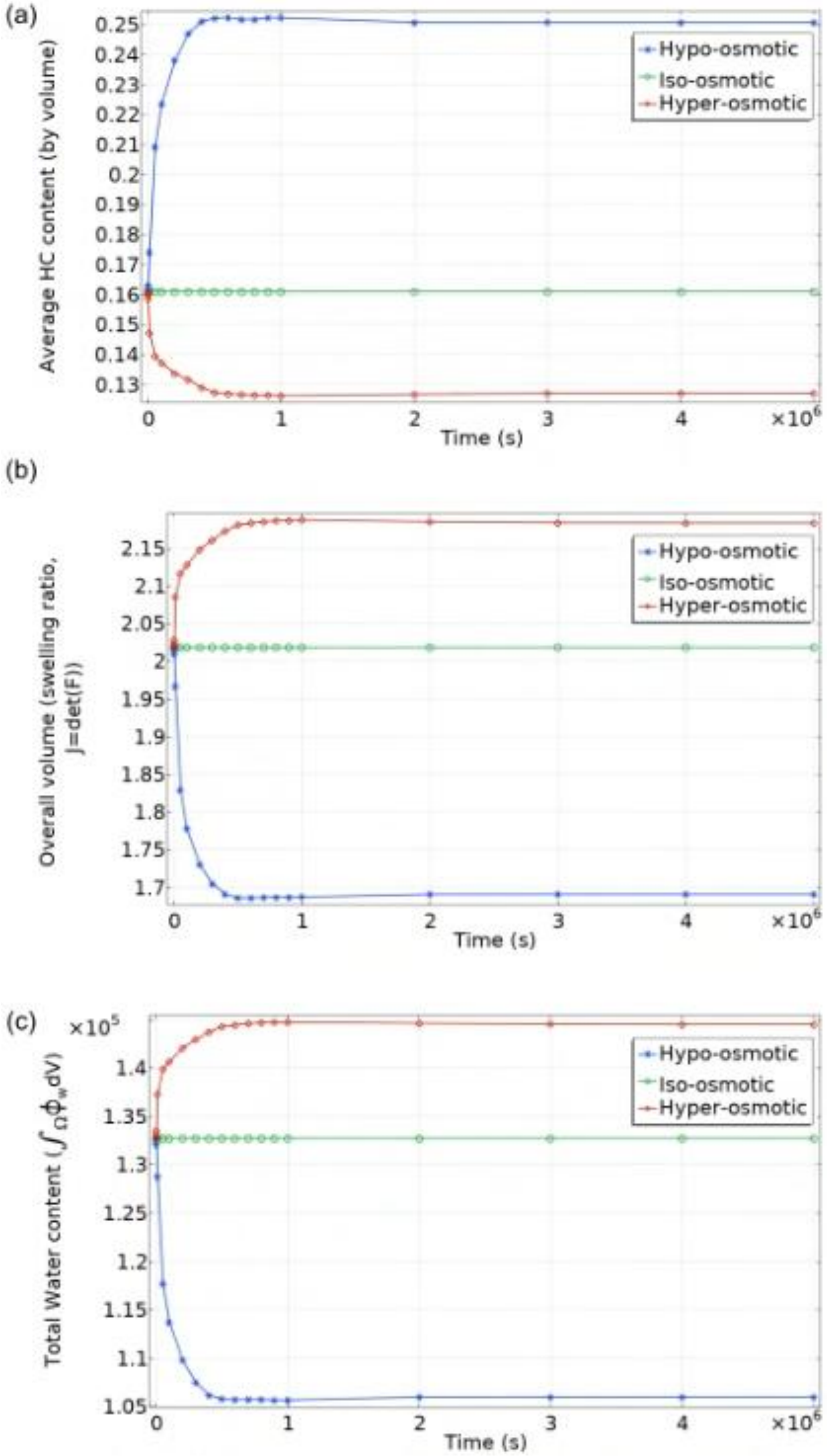

Fig S21: Simulation predicted changes in the (a) overall HC content (by percent volume) in the nucleus, (b) Total volume of the nucleus, and (c) total water content in the nucleus after hyper-, iso-, and hypo-osmotic loading.

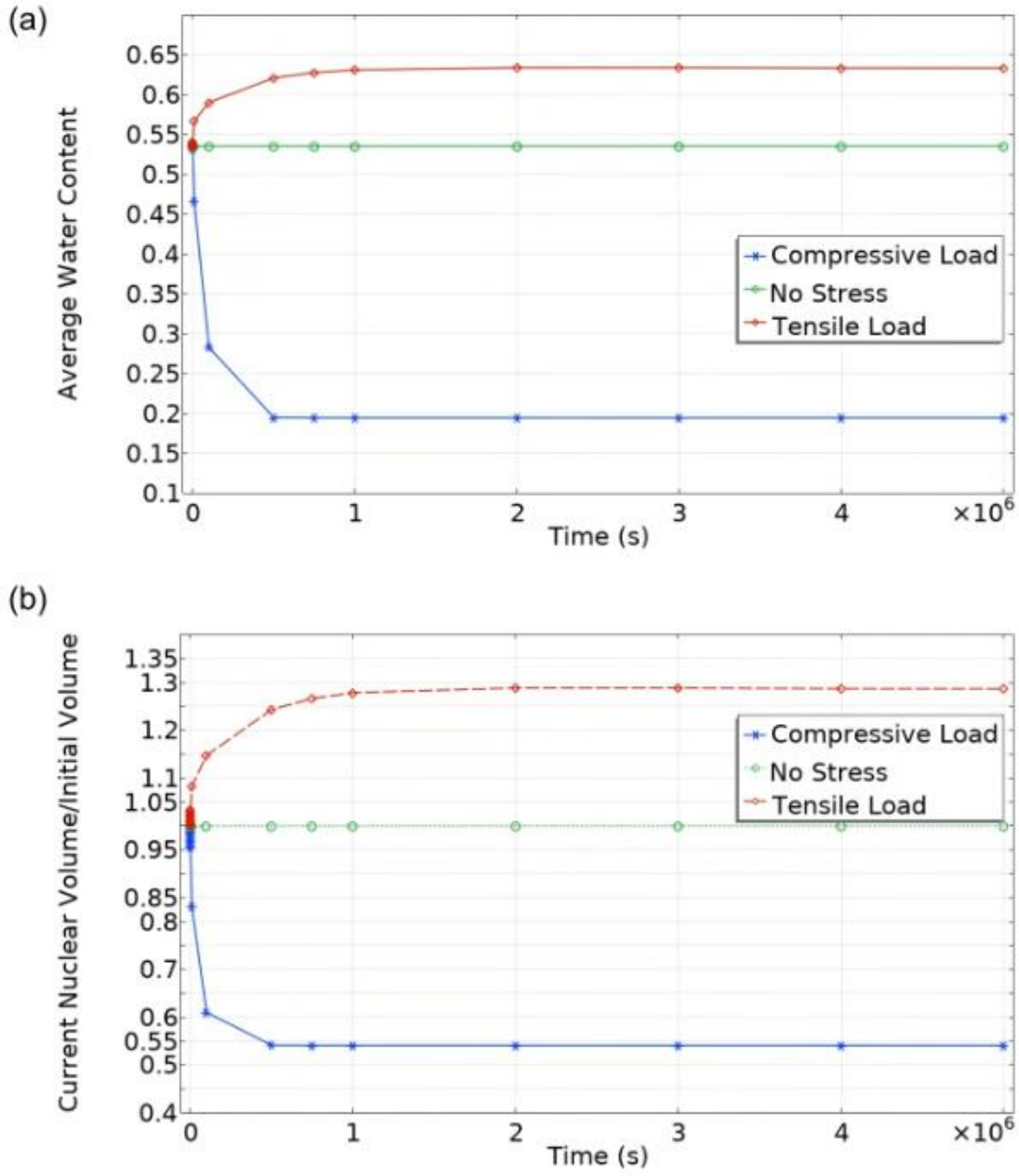

Fig S22: Simulation predicted changes in the (a) total water content in the nucleus, (b) Total volume of the nucleus, after tensile and compressive loads.

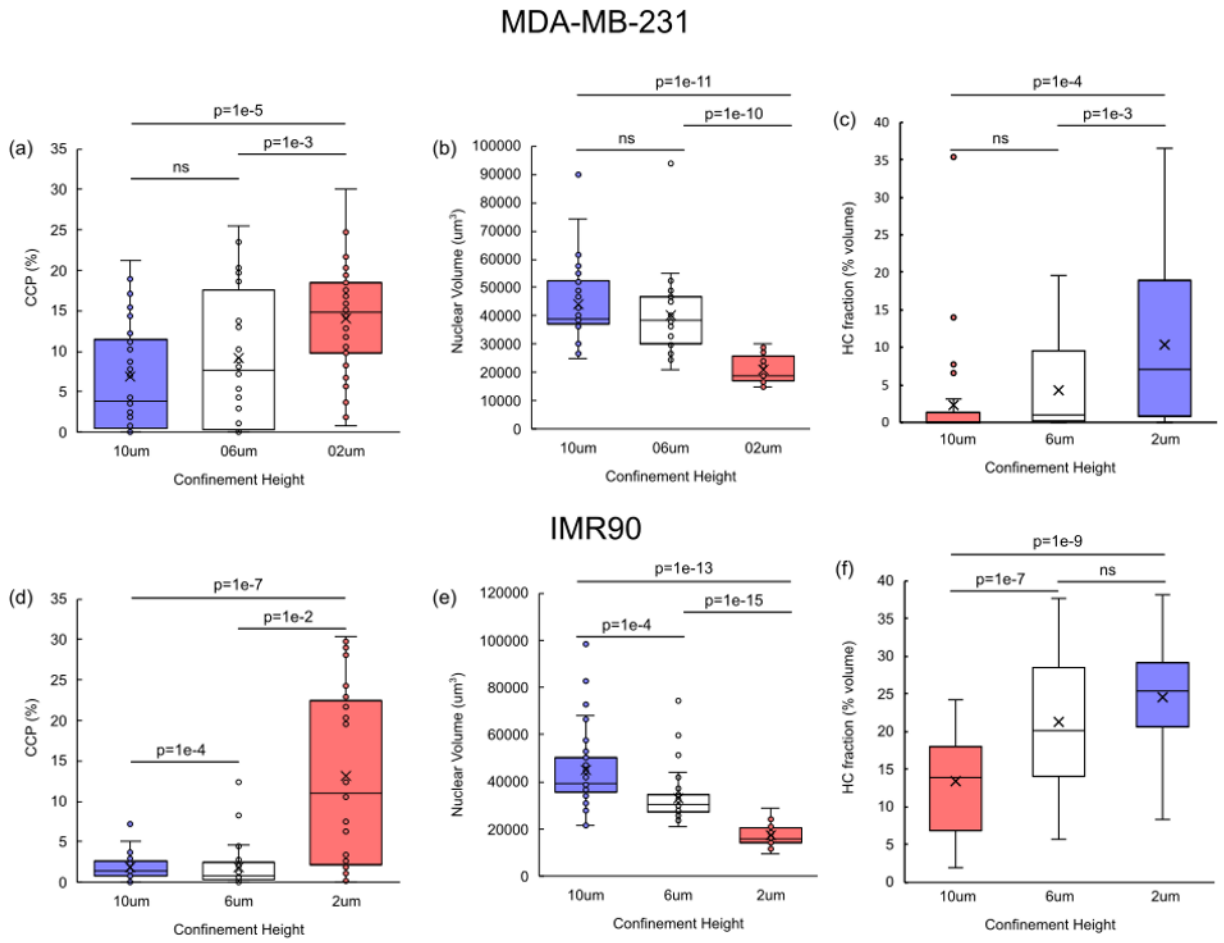

Fig S23: (Refers to Fig 4c-f in the main manuscript) Quantification of MDA-MB-231 (top row) and IMR (bottom row) nuclei in confinements of 10 $\mu m$, 6$\mu m$ and 2 $\mu m$. Progressively increasing the nuclear compression results in increase in chromatin compaction parameter (CCP) and fraction of nucleus (by volume) occupied by HC, while decreasing the nuclear volume. All significance values obtained via unpaired two-tailed t-test. (Number of MDA-MB-231 nuclei: 34, 36 and 35 nuclei respectively for 10$\mu m$, 6$\mu m$ and 2$\mu m$ cases; Number of IMR90 nuclei: 39, 44 and 38 nuclei respectively for 10$\mu m$, 6$\mu m$ and 2$\mu m$ cases)

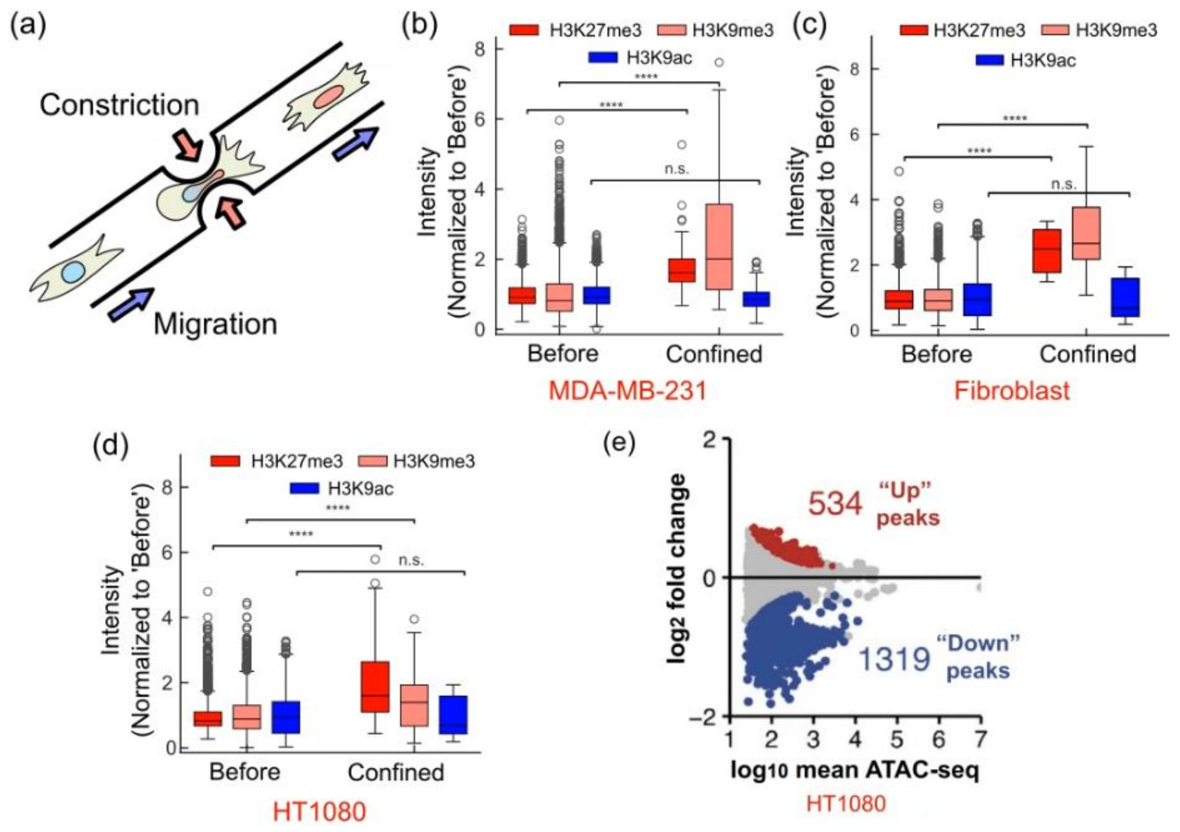

Fig S24: (Refers to Section 2.5): Nuclear compression due to confined migration of cells. (a) A representative schematic demonstrating the confined migration of cells through constrictive pores. The confined migration results in increased HC markers (H3K27me3 and H3K9me3) while H3K9ac levels stay unchanged. These changes are quantified in (b) MDA-MB-231 cells, (c) SV40 human fibroblasts, and (d) HT1080 cells demonstrating that these observations are not restricted to a single cell type, suggesting a cell-independent mechanism, as predicted by our simulations. All p-values in (b)-(d) are obtained via unpaired two-tailed t-test. **** indicates p-values <1e-10, and n.s. indicates p>0.5. (e) Confined migration of HT1080 through collagen matrices with smaller pore size results in reduced chromatin accessibility, consistent with model predictions of increased HC content (reproduced from [25]).

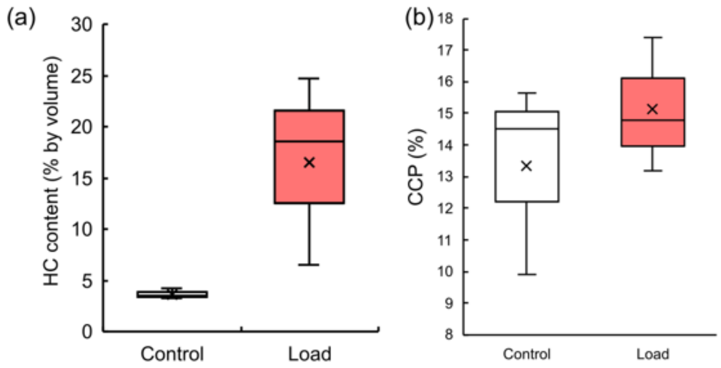

Fig S25: (Refers to Fig 5b,c in the main manuscript) Quantification of chromatin reorganization upon application of compressive load on NIH3T3 nuclei through (a) increase in percent volume of nucleus occupied by the heterochromatin phase, and (b) chromatin condensation parameter (CCP).

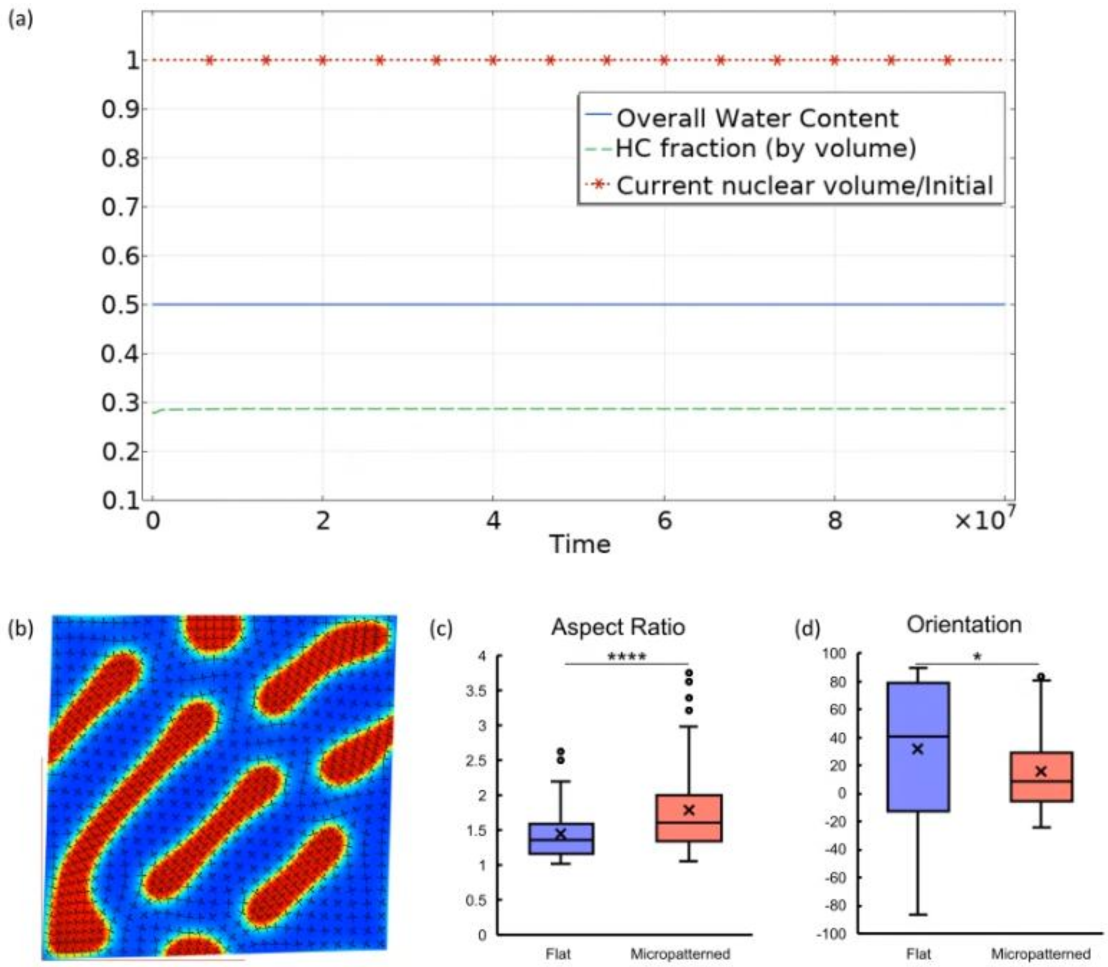

Fig S26: (Refers to Fig 6a-f in the main manuscript) (a) Application of shear loads does not alter overall water content in the nucleus, the nuclear volume and the fraction of total area occupied by the HC phase confirming that these loads are volume preserving, and thus the reorganization of chromatin is not due to nucleoplasm influx or efflux from the nucleus. (b) The chromatin organization under high shear shown in Fig 6g, right panel in the main manuscript, with the principal directions overlaid as black arrows. Note that the orientation of the HC domains exactly matches the first principal direction (with second principal direction always perpendicular to the first principal direction). (c-d) Additional quantification of HC domain reorientation in response to anisotropic uniaxial compression, in hMSCs cultured on flat and micropatterned glass substrates. (c) The aspect ratio of HC domains, measured as ratio of major to minor axes of circumscribing ellipse (n = 29 and 30 nuclei on flat and micropatterned substrates respectively; unpaired, two-tailed t-test, p=1e-6). (d) Orientation of the HC domains. A narrower range of orientation on micropatterned glass substrate indicates more coherent alignment between the HC domains – as predicted by our model.

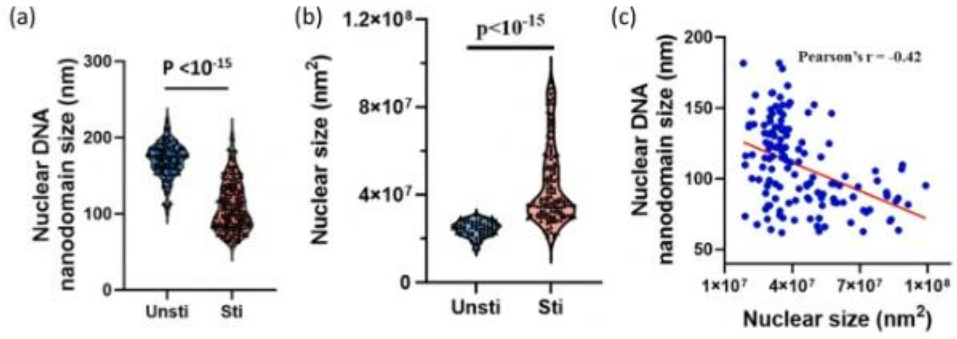

Fig S27: (Reproduced from Xu, et al. [26]) Chromatin nanodomains and nuclear size changes in T cells in quiescent (Unsti) and activated (Sti) states quantified after super-resolution dSTORM imaging of DNA labeled with small-molecule fluorescent probes. (a) The average nuclear DNA nanodomain size showed a reduction upon activation suggesting chromatin decondensation, and (b) was accompanied by an increase in nuclear size. (c) The DNA nanodomain size and nuclear size are inversely correlated. This observation agrees with our model's prediction that chromatin decondensation (due to epigenetic, osmotic or mechanical perturbations) is accompanied by increase in nuclear volume and vice-versa.
